# From chromosome-scale assembly to the mappable genome of rubber dandelion: chromosome reorganization, rubber-pathway coordination, and domestication traits

**DOI:** 10.64898/2026.10.23.761435

**Authors:** Pablo Vargas-Mejía, Camila Gutierrez-Manriquez, Elizabeth S. Anderson, Daniel R. Williams, Julio Vega-Arreguín, Jonathan Fresnedo-Ramírez

## Abstract

Natural rubber supply depends almost entirely on one tropical tree, motivating development of temperate alternatives. Rubber dandelion (*Taraxacum kok-saghyz* Rodin) is the leading candidate, with comparatively well characterized rubber biosynthesis pathways, but its large, repeat-rich genome has left breeding genomics far behind pathway biology. In response, we assembled OSU TarKok 3.0, a 1.19-Gb nuclear reference in which eight chromosome-scale scaffolds contain 98% of the sequence. Continuity within chromosomes improved markedly, with contig N50 rising from 1.3 to 37.8 Mb, and conserved gene-space recovery improved accordingly. Macrosynteny showed strong chromosome conservation within Taraxacum and, when compared to lettuce, displayed four nearly intact and four composite chromosomes, a signature compatible with descending dysploidy accompanied by rearrangement rather than a single end-to-end fusion. Across nine Cichorieae genomes, none of six surveyed rubber-related gene families was uniquely enlarged in rubber dandelion. Instead, a *cis*-prenyltransferase locus, a rubber transferase activator, and three small rubber particle protein genes were enriched in underground tissues and grouped with rubber elongation factor in a latex-associated co-expression module, favoring coordinated pathway deployment over species-specific family expansion. Intersecting uniquely mappable and nonrepetitive sequence defined the mappable genome, approximately 210 Mb or 18% of the assembly, within which markers can be interpreted allelically. Mapping in an independent F₂ family of 83 plants identified a chromosome 1 locus for filled-seed production under restricted cross-pollination, whereas herbicide response yielded no significant locus in this design. OSU TarKok 3.0 links chromosome reorganization, pathway deployment, and domestication traits into one framework for improving an outcrossing crop.

**Core Ideas:**

1. OSU TarKok 3.0 markedly improves within-chromosome continuity and conserved gene-space recovery.
2. Repeat-aware defines a ∼210 Mb short-read mappable space (18% of the assembly), enabling reliable marker analysis.
3. Four chromosomes retain conserved relationships, supporting dysploidy with rearrangement beyond simple fusion.
4. No rubber-related gene family is uniquely expanded; rubber production reflects pathway coordination, not dosage.
5. A chromosome 1 QTL for selfing-related filled-seed production overlaps the positional candidate *g7174*.

**Plain Language Summary:** As the global supply of natural rubber is dependent on one tropical tree, rubber dandelion has been proposed as a temperate alternative. Although scientists already understand many of the genes involved in its rubber production, breeding has been slowed by its large, highly repetitive genome. In response, we built OSU TarKok 3.0, a much more continuous genome that places 98% of the sequence on eight chromosomes. Comparison with related dandelions revealed broad chromosome conservation, while comparison with lettuce was consistent with rearrangements during the reduction from nine to eight chromosomes. Across nine members of the sunflower family, none of six rubber-related gene families were uniquely enlarged in rubber dandelion. Instead, genes involved in building rubber chains and particles were active together in underground tissues and latex-associated samples, suggesting that coordinated gene activity matters more than extra gene copies. We also identified ∼210 million base pairs, 18% of the genome, where short-read genetic markers can be interpreted reliably. Within that space, we mapped a chromosome 1 region associated with producing filled seed when cross-pollination was restricted. This region overlaps a candidate gene from a protein family involved in reproductive development in other plants, providing a testable target for improving seed production.

## Introduction

Natural rubber combines elasticity, resilience, and fatigue resistance in ways that synthetic polymers still do not fully reproduce, and it remains essential to tens of thousands of products, including those used in applications in which material failure cannot be tolerated (Cornish, 2017; Cherian et al., 2019). Nearly all natural rubber is harvested from *Hevea brasiliensis* (Willd. ex A.Juss.) Müll.Arg., which is cultivated from a narrow clonal base in a small number of tropical production regions. This concentration exposes a strategic industrial material to disease, land-use change, and geopolitical disruption (van Beilen & Poirier, 2007; Cornish, 2017). Rubber dandelion (*Taraxacum kok-saghyz* Rodin; Asteraceae, tribe Cichorieae) is a promising temperate alternative. It accumulates high-molecular-weight natural rubber in root laticifers, can be propagated by seed, and can be cultivated as an annual crop under temperate field conditions.

The biochemistry of rubber synthesis in this species is increasingly well characterized (Luo et al., 2017). *Cis*-prenyltransferases (CPTs) elongate the polyisoprene chain within a membrane-associated complex with *cis*-prenyltransferase-like/rubber transferase activator proteins (CPTL/RTAs), while small rubber particle proteins (SRPPs) and rubber elongation factor (REF) contribute to rubber-particle formation, stability, and accumulation (Hillebrand et al., 2012; Epping et al., 2015; Niephaus et al., 2019). Recent work has resolved individual components of that system. TkCPT1 and TkCPT2 form complexes with TkCPTL1 that produce extra-long-chain polyisoprenes, whereas other family members show different tissue distributions and products (Müller et al., 2025). CRISPR disruption of *TkCPT1* reduces rubber production, and simultaneous disruption of *TkCPT1* and *TkCPT2* blocks both rubber biosynthesis and rubber-particle formation (Xu et al., 2025). Jasmonate-responsive and direct transcriptional regulation of rubber-particle genes has also been demonstrated (Wu et al., 2024; Fantini et al., 2025; Fang et al., 2026). Together, these studies provide a functional basis for testing how gene-family dosage and coordinated regulation contribute to rubber production.

That mechanistic progress has not been matched by equivalent advances in breeding genomics, and the resulting asymmetry now limits the crop more than any remaining gap in pathway biology. Unimproved rubber dandelion establishes slowly, competes poorly with weeds, and can suffer substantial yield loss under weed pressure, while sensitivity to many broadleaf herbicides restricts selective chemical control during the vulnerable establishment window (Iaffaldano et al., 2018; Pannwitt et al., 2025). The species is also predominantly outcrossing (Kirschner et al., 2013), and microscopy of modern material has demonstrated a sporophytic self-incompatibility response that restricts self-pollen progression (Wollenweber et al., 2021), a mating system shared with other members of Asteraceae in which *S*-locus diversity and dominance interactions constrain crossing design (Hiscock et al., 2003; Brennan et al., 2011). Outcrossing preserves genetic diversity, but it complicates the fixation of favorable allele combinations, the development of uniform lines, and the multiplication of seed without synchronized compatible parents. Germplasm surveys nevertheless indicate that the capacity to set selfed seed varies among individuals (Ponciano et al., 2026), and cytological analysis confirms predominantly regular sexual sporogenesis and gametogenesis (Schuchovski et al., 2026). These observations make selfing-related reproductive capacity a plausible target for genetic analysis and selection. Domesticating a predominantly outcrossing species is, therefore, a distinct genetic problem from improving an established inbred crop, and one in which reproductive predictability and field establishment, not product chemistry, are frequently the limiting steps (Van Tassel et al., 2020).

Genomics-assisted breeding is further complicated by a repeat-rich nuclear genome of approximately 1.2 Gb. Repeats fragment de novo assemblies, and obscure recent paralogs can allow short reads originating from related copies to be mistaken for allelic variation, producing spurious heterozygosity and unstable marker behavior (McKinney et al., 2017). The TK1151 assembly established chromosome-scale sequence for the species (Lin et al., 2022), but chromosome-scale organization and within-chromosome continuity are not the same property. A scaffold assembled to chromosome length may still contain many internal breaks that divide gene neighborhoods, recent paralogs, syntenic blocks, and candidate intervals. The practical consequence is rarely stated explicitly: in a repeat-rich genome, only part of the sequence can be genotyped reliably from short reads, and that fraction is usually left implicit in marker studies. We refer to that fraction of the sequence that is both uniquely mappable and nonrepetitive, and therefore interpretable at the allelic level, as the short-read mappable space or genome. Delimiting it identifies where markers can be trusted and assays can be designed, bounds the interpretation of negative association results, and highlights regions that long-read genotyping or a multi-accession pangenome will need to resolve (Pockrandt et al., 2020; Della Coletta et al., 2021).

Greater continuity also makes three questions tractable that are difficult to address with a fragmented reference. At chromosome scale, *Taraxacum* and *Lactuca* exemplify the eight-and nine-chromosome base numbers found within Cichorieae, respectively. These differences occur within a deeper Asteraceae history of paleopolyploidy, fractionation, and rearrangement (Feng et al., 2026). Cytogenetic and phylogenomic evidence from *Reichardia* supports descending dysploidy within the tribe, making an ancestral nine-to-eight transition plausible but leaving its realization in *Taraxacum* unresolved. Evaluating this hypothesis benefits from a reference that preserves syntenic blocks and the interstitial sequence between them. At pathway scale, whether rubber production reflects lineage-specific gene dosage or coordinated deployment of a broadly conserved system cannot be settled by pairwise comparison. Earlier work reported relative expansion of CPT and SRPP families in TK1151 relative to *T. mongolicum* Hand.-Mazz. (Lin et al., 2022), but a family can be larger than in one comparator without being unusual across the tribe. Distinguishing these possibilities requires uniform criteria applied to many genomes together with tissue and co-expression evidence. At breeding scale, the recombination landscape determines where a locus can be resolved efficiently and where linked variation will remain difficult to separate, a constraint that shapes selection strategy in every crop (Taagen et al., 2020; Brazier & Glémin, 2022). While previous studies have produced the first linkage map and, more recently, a high-density SNP map with rubber-content QTLs in rubber dandelion (Arias et al., 2016; Yang et al., 2023), integrating new trait mapping with a more continuous physical reference can improve the interpretation of markers, recombination, and candidate intervals.

We therefore developed OSU TarKok 3.0 as a common coordinate system for the biological and breeding questions above. Our objectives were to (i) quantify gains in assembly continuity and gene-space completeness relative to available references; (ii) resolve chromosome correspondence across Cichorieae and test whether each rubber dandelion chromosome corresponds predominantly to one comparator ancestral chromosome or combines segments with different chromosomal affinities; (iii) compare six rubber-related gene families across nine Cichorieae genomes under uniform criteria and integrate copy number with tissue expression and co-expression to distinguish family expansion from coordinated deployment; (iv) define the short-read mappable genome and characterize the recombination landscape within it; and (v) demonstrate the utility of that framework by mapping two domestication-relevant endpoints in an independent F₂ family: filled-seed production from bagged capitula treated as a selfing-related operational phenotype and survival following a single pre-emergence application of the protoporphyrinogen oxidase (PPO)-inhibiting herbicide (Dayan et al., 2018). Because the available cohorts are modest, comprising a 181-member association panel and 83 independent F₂ plants, the mapping analyses are proof-of-utility demonstrations bounded by cohort size, relatedness, ascertainment, and phenotype design rather than definitive locus-discovery experiments. By connecting chromosome structure, pathway deployment, and segregating domestication traits within one explicitly bounded reference, this study moves rubber dandelion genomics from description toward a platform for sustainable genetic improvement of an outcrossing crop.

## Materials and Methods

### Plant materials, cohorts, and phenotyping

The study included 187 sequenced *T. kok-saghyz* individuals. Complete consensus genotypes were obtained for 186, and the 181 individuals with matched genotype and trait records conformed to the final population panel. The panel comprised one family founder, 83 independent F2, 77 F3, and 20 additional accessions. Because the F3 are descendants of selected F2 plants and are therefore not independent of the F2 generation, linkage scans were restricted to the 83 F2. Filled-seed scores were available for 166 panel members, and herbicide scores were available for 177 (145 tolerant and 32 susceptible; four unscored). The herbicide imbalance reflects survival selection before genotyping because susceptible plants were less likely to be advanced and sampled. Cohort membership for every analysis is given in Supplemental Table S1.

The filled-seed endpoint recorded the presence or absence of filled seed on bagged capitula without a directed cross. Among the 181-member panel, 166 individuals were scored: 84 had filled seed and 82 did not. This binary observation was analyzed as an operational endpoint and not as a reproductive mechanism. Herbicide response was scored after a single pre-emergence application of sulfentrazone [*N*-[2,4-dichloro-5-[4-(difluoromethyl)-3-methyl-5-oxo-1,2,4-triazol-1-yl]phenyl]methanesulfonamide] at 280 g active ingredient ha−1. Sulfentrazone inhibits protoporphyrinogen oxidase, causing light-dependent membrane peroxidation in emerging seedlings (Dayan et al., 2018); the pre-emergence timing therefore places the injury window during crop establishment. Emergence and survival through the first true leaf defined tolerance, whereas failure to emerge or death before that stage defined susceptibility. All 83 independent F₂ plants were scored for both endpoints: 48 produced filled seed and 35 did not, whereas 53 were tolerant and 30 were susceptible. The two scans therefore used the same individuals, genotype data, marker set, and family structure and differed only in the phenotype tested. OH-1004 (listed as OH_1004 in the association panel), was one of the 20 non-family accessions and was scored for both trait endpoints.

Because the association panel (n = 181) and independent F2 family (n = 83) were the available cohorts for this genomic-resource study, these analyses were designed as proof-of-utility demonstrations rather than definitive locus-discovery experiments; inference was therefore restricted to prespecified thresholds and explicitly bounded by cohort size, relatedness, ascertainment, and phenotype design.

### Genome sequencing, assembly, and evaluation

High-molecular-weight DNA was extracted from OH-1004, an inbred line that is susceptible to sulfentrazone and that served as the susceptible control in every herbicide screen and was sequenced with PacBio HiFi chemistry at approximately 36-fold coverage. OSU TarKok 3.0 was generated within the same accession-specific assembly project as OSU TarKok 2.0, and all bioinformatic analyses were performed under the infrastructure of the Ohio Supercomputer Center (1987). HiFi reads were assembled with Verkko (Rautiainen et al., 2023), and the resulting contigs were scaffolded with RagTag using the Omni-C-supported chromosome organization of OSU TarKok 2.0 (Alonge et al., 2022). Remaining unplaced contigs were reconciled with RagTag using OSU TarKok 2.0 as the primary guide, and remaining gaps were patched with sequence from an independent hifiasm assembly and OSU TarKok 2.0 (Cheng et al., 2021). No new long-range library was generated for OSU TarKok 3.0. Chromosome order therefore inherits long-range evidence from the same accession, while read depth, telomeric arrays, and independent synteny provide complementary checks. Full software versions and parameters are in Supplemental Methods.

Organellar sequences were separated before nuclear-assembly statistics and annotation counts were finalized. Assembly length, scaffold and maximal N-split contig Nx, gaps, GC content, read depth, and telomeric arrays were evaluated on the final 658-sequence organellar-filtered nuclear assembly. Repeat occupancy, transposable-element occupancy, and LTR Assembly Index (LAI) were calculated on the 857-sequence primary assembly before organellar partitioning. Assembly completeness was evaluated with BUSCO v6.0.0 using the eudicotyledons_odb12 lineage dataset (n = 2,805 (Manni et al., 2021)). Genome-mode BUSCO was evaluated for OSU TarKok 3.0 and TK1151 and is reported in Table 1. Predicted-proteome completeness was evaluated separately in protein mode using the final OSU TarKok 3.0 nuclear representative proteome of 34,637 proteins, and the published TK1151 proteome of 47,427 protein sequences; these results are also reported in Table 1.

**Table 1.** Assembly continuity and conserved gene-space completeness of OSU TarKok 3.0 and TK1151. Contigs are maximal non-N sequence segments obtained by splitting scaffolds at N-runs; anchored fraction is the percentage of total assembly length contained in chromosome-scale scaffolds.

| Assembly characteristic | OSU TarKok 3.0 | TK1151 |
| --- | --- | --- |
| Assembly length (bp) | 1,192,836,847 | 1,101,786,192 |
| FASTA sequences/scaffolds | 658 | 9 |
| Contigs (sequences split at N-runs) | 746 | 1,926 |
| Chromosome-scale scaffolds | 8 | 8 |
| Length in chromosome-scale scaffolds (bp) | 1,168,509,718 | 1,073,873,677 |
| Fraction anchored | 98.0% | 97.5% |
| Scaffold N50 / N90 (Mb) | 152.9 / 117.9 | 131.6 / 111.4 |
| Contig N50 / N90 (Mb) | 37.8 / 8.6 | 1.3 / 0.3 |
| Largest scaffold / contig (Mb) | 178.2 / 117.7 | 162.5 / 6.0 |
| Gaps (N-runs) | 88 | 1,917 |
| N bases | 8,636 | 17,324,794 |
| N bases per 100 kb | 0.72 | 1,572.43 |
| GC content (%) | 38.0 | 36.9 |
| Genome-mode BUSCO complete (single, duplicated) | 98.4% (91.6, 6.8) | 94.2% (77.6, 16.6) |
| Protein sequences evaluated by BUSCO | 34,637 | 47,427 |
| Protein-mode BUSCO complete (single-copy, duplicated) fragmented, missing | 97.5% (92.1, 5.4) 1.3, 1.2 | 73.5% (45.2, 28.2) 8.8, 17.7 |

### Annotation, comparative genomics, and gene-family evolution

Repeats were modeled with RepeatModeler2, classified with TEsorter, and masked with RepeatMasker; structural transposable elements and the LTR Assembly Index were evaluated independently. BRAKER3 integrated RNA-seq and protein evidence for nuclear gene prediction (Flynn et al., 2020; Zhang et al., 2022; Gabriel et al., 2024). Comparative chromosome structure was evaluated with ntSynt against TK1151, *Taraxacum mongolicum* Hand.-Mazz., and lettuce (*Lactuca sativa* L.) (Coombe et al., 2025). A nine-genome phylogeny used 1,480 shared complete single-copy BUSCO proteins aligned with MAFFT, trimmed with trimAl, and analyzed with IQ-TREE with branch support assessed using 1,000 ultrafast bootstrap replicates (Capella-Gutiérrez et al., 2009; Katoh & Standley, 2013; Hoang et al., 2018; Wong et al., 2026). Nucleotide comparisons retained best alignments from 5-kb query fragments only when the aligned span was at least 2 kb.

Candidate biosynthetic gene clusters were screened with plantiSMASH v2.0.4 (Del Pup et al., 2026). Rubber-pathway homologs were projected uniformly to nine assemblies with Miniprot using fixed identity and coverage criteria (Li, 2023). Projected homology clusters were kept distinct from curated expression loci and global orthogroups. OrthoFinder analyzed eight one-protein-per-gene proteomes and CAFE5 tested family-size evolution under the selected single-global-rate, no-error model (Emms & Kelly, 2019; Mendes et al., 2020). Family P values were adjusted by the Benjamini–Hochberg method (Benjamini & Hochberg, 1995) across all 24,025 tested families; branch probabilities were retained only as unadjusted descriptive model outputs. Genome and proteome accessions, input versions, and the analyses performed for each taxon are listed in Supplemental Table S2.

### RNA sequencing and co-expression

RNA was obtained from leaf, flower, crown, root, and pooled floral and seed material. The in-house experiment comprised 12 Illumina libraries: two each from leaf, flower, crown, and root and four pooled floral/seed libraries. featureCounts produced gene counts; TPM was used for display, and DESeq2 tested underground (crown and root; n = 4) versus aerial (leaf, flower, and pooled floral/seed; n = 8) expression with false-discovery control (Liao et al., 2014; Love et al., 2014). Enrichment required an underground:aerial ratio greater than one and adjusted P < 0.05.

The primary weighted co-expression analysis used the combined 36-library dataset; the separately fitted 12-library network was treated as a sensitivity analysis (Langfelder & Horvath, 2008). Primary module– tissue P values were adjusted across all 259 tests comprising 37 modules and seven tissue categories. The displayed 36-library network used undirected topological overlap measure (TOM) edges selected as the two strongest links per displayed node plus core-locus pairs with TOM ≥ 0.27, after deterministic tie, missing-value, and duplicate handling. All latex libraries were public and therefore confounded with study, genotype, and protocol. Full edge rules and motif tests are given in Supplemental Methods.

### Single-copy genotyping, association, and F2 mapping

Variants generated from seqWell-prepared genomic libraries were called independently with GATK HaplotypeCaller and BCFtools, and exact allele agreement defined the caller consensus (McKenna et al., 2010; Danecek et al., 2021). GenMap-defined uniquely mappable sequence was intersected with non-repeat sequence to establish the conservative mappable genome (Pockrandt et al., 2020). Consensus variants were filtered for allele frequency, missingness, excess heterozygosity, depth, and Mendelian behavior in selfed families. Population association used a GEMMA linear mixed model (Zhou & Stephens, 2012), GCTA MLMA-LOCO (Yang et al., 2011), and PLINK2 logistic regression with Firth fallback (Firth, 1993; Chang et al., 2015). All three models included the first five principal components as covariates. GEMMA and GCTA used a Bonferroni threshold of P = 0.05/1,099,707, whereas PLINK2 used phenotype-specific 5% genome-wide thresholds obtained from 1,000 phenotype permutations. An exact chromosome:position:reference:alternate allele record was considered cross-model supported only when it passed its model-specific threshold in at least two of the three models. A locus was required to meet the prespecified genome-wide support in at least two models. Principal component analysis (PCA) used 181 individuals and FST used the 166 filled-seed-scored individuals.

The independent-F2 scan used additive logistic likelihood-ratio tests at founder-heterozygous, single-copy markers. LOD was the full-minus-null log likelihood divided by ln(10), and experiment-wise thresholds were the 95th percentiles of maximum LOD from 1,000 phenotype permutations. Connected 1.5-LOD support bounds were interpolated linearly between adjacent marker scores on each side of the peak. Lep-MAP3 constructed a sex-averaged linkage map with a Kosambi mapping function (Rastas, 2017); Marey curves used chromosome-dominant markers and isotonic regression. The same 83 F2 and marker framework were used for both operational endpoints.

## Results

### A chromosome-scale nuclear resource substantially improves continuity within chromosomes

The HiFi-read 21-mer spectrum showed a principal homozygous peak near 35× coverage. Because the direct GenomeScope2 fit did not capture the extensive high-copy repeat tail, genome size was estimated from total k-mer abundance and peak coverage at approximately 1.08 Gb, about 112 Mb smaller than the final nuclear assembly (Supplemental Figure S1). OSU TarKok 3.0 comprises 1,192,836,847 bp in 658 nuclear sequences, with eight chromosome-scale scaffolds accounting for 1,168,509,718 bp, or 98.0% of the assembly. Because OSU TarKok 3.0 and the published TK1151 are both organized at chromosome scale, their scaffold N50 values are relatively similar (152.9 and 131.6 Mb, respectively). The larger improvement is evident within chromosomes: contig N50 increased from 1.3 Mb in TK1151 to 37.8 Mb in OSU TarKok 3.0, contig N90 increased from 0.3 to 8.6 Mb, and the largest contig increased from 6.0 to 117.7 Mb (Table 1; Figure 1A). At chromosome level, contig N50 values for Chr1 through Chr8 were 26.9, 42.0, 37.8, 17.8, 31.0, 51.5, 65.8, and 117.7 Mb, respectively, and the corresponding scaffolds contained 12, 9, 8, 15, 7, 26, 7, and 4 gaps. Across TK1151 ChrA1 through ChrA8, contig N50 values ranged from 1.1 to 1.7 Mb and individual pseudomolecules contained 128 to 262 gaps; the separate 27.9-Mb ChrA0 sequence bin contained an additional 401 gaps.

**Figure 1.**
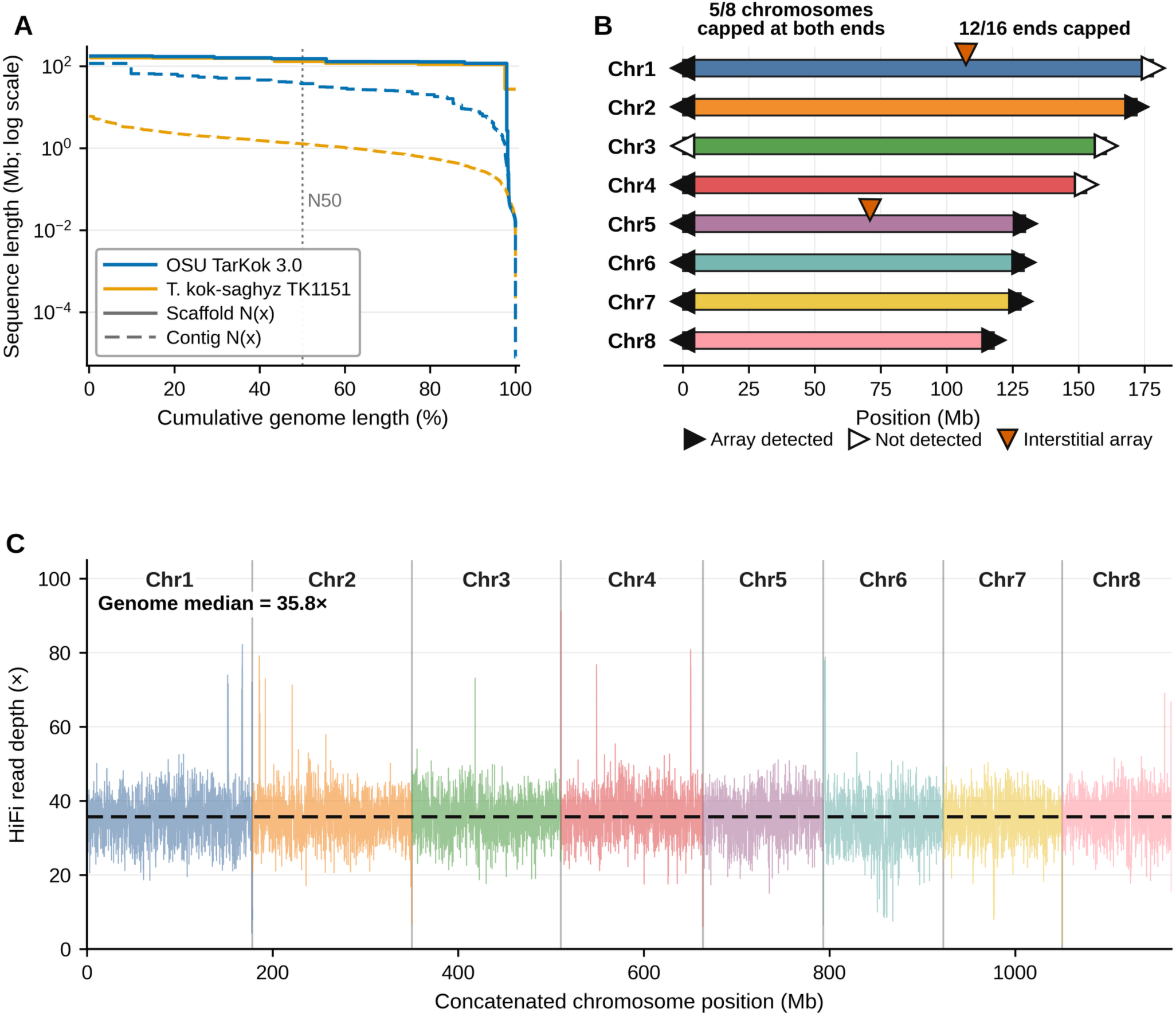
Assembly continuity and chromosome-scale support for OSU TarKok 3.0. (A) Scaffold and contig Nx profiles for OSU TarKok 3.0 and TK1151. The OSU curves were calculated from the nuclear assembly and its 746 maximal N-split contigs; TK1151 was evaluated under the same definition. (B) Terminal and interstitial telomeric arrays on the eight OSU chromosome scaffolds; arrays were detected at both ends of five chromosomes and at 12 of 16 ends. (C) PacBio HiFi read depth in 100-kb windows.

The reduction in unresolved sequence was equally pronounced. Splitting scaffolds at gaps produced 746 contigs in OSU TarKok 3.0, compared with 1,926 in TK1151. OSU TarKok 3.0 contains 88 gap regions and 0.72 N bases per 100 kb, whereas TK1151 contains 1,917 gaps and 1,572.43 N bases per 100 kb (8,636 and 17,324,794 N bases in total, respectively; Table 1). Telomeric-repeat arrays were detected at 12 of 16 chromosome ends, including both ends of five chromosomes, and interstitial arrays were present on Chr1 and Chr5 (Figure 1B; Supplemental Figure S2). HiFi depth was broadly uniform across the eight chromosome scaffolds (Figure 1C). Together, telomeric-repeat recovery and uniform read depth provide complementary support for the continuity and structural organization of the chromosome-scale assembly.

GC-versus-coverage profiling distinguished the eight nuclear chromosome scaffolds, centered near 38% GC and 36× coverage, from high-copy organellar sequences. The 198 sequences assigned to the organellar set were excluded before nuclear assembly statistics and annotation were finalized. Final assembly quality control also removed one 25,466-bp unplaced sequence classified as ascomycete contamination and lacking annotated protein-coding genes (Supplemental Figure S3). A separate analysis of plastid-recruited HiFi reads produced a 151,316-bp reference-guided chloroplast consensus comprising an 83,900-bp large single-copy region, an 18,538-bp small single-copy region, and two 24,439-bp inverted repeats (Supplemental Figure S4).

The 857-sequence primary assembly evaluated before organellar partitioning was highly repetitive: repetitive sequence occupied 77.5% of the assembly, and 59.8% was classified as transposable-element sequence. LTR retrotransposons were the predominant components, led by Gypsy and Copia elements, which occupied 20.5% and 16.9% of the assembly, respectively; the length-weighted LTR Assembly Index was 41.6 (Supplemental Figure S5).

BRAKER3 integration of the 12-library RNA-seq evidence with a eudicot protein reference set produced a final nuclear annotation of 34,637 protein-coding genes represented by 39,326 mRNAs, a mean of 1.14 transcripts per gene. Of these genes, 34,610 (99.92%) lie on the eight chromosome-scale scaffolds, whereas 27 (0.08%) lie on unplaced sequences. Gene models averaged 2,129 bp, with a mean of 4.67 coding exons per transcript and a median intron length of 94 bp. Of the predicted transcripts, 25.28% were monoexonic and 43.87% contained both annotated 5′ and 3′ UTRs. Functional evidence supported most of the representative protein set. InterPro annotation was recovered for 80.94%, at least one Gene Ontology term for 69.67%, and an eggNOG orthologous-group assignment for 89.22%. Across InterPro, Pfam, Gene Ontology, eggNOG, and KEGG evidence, 89.87% of representative proteins received at least one functional assignment, leaving 10.13% without one (Supplemental Table S3). Non-coding RNA annotation retained 698 non-pseudogene tRNA predictions and 1,502 additional structured RNA predictions, comprising 175 snRNA, 920 snoRNA, 98 miRNA, and 309 other Rfam-family predictions. The much larger set of rRNA-model calls was retained as uncurated predictions rather than interpreted as a gene count because its abundance indicated extensive repeat-associated matches (Supplemental Table S3).

Genome-mode BUSCO recovered 98.4% complete orthologs from OSU TarKok 3.0, including 91.6% single-copy and 6.8% duplicated orthologs compared with 94.2% from TK1151. The contrast was larger in protein mode: BUSCO recovered 97.5% complete orthologs from the OSU TarKok 3.0 nuclear proteome, including 92.1% as single-copy, compared with 73.5% complete and 45.2% single-copy orthologs in the TK1151 proteome. Fragmented and missing orthologs together accounted for 2.5% of the OSU TarKok 3.0 proteome and 26.5% of the TK1151 proteome (Table 1). Thus, the improvement in sequence continuity is accompanied by substantially more complete recovery of conserved eudicot gene space.

Genome and proteome completeness were evaluated with BUSCO v6.0.0 against eudicotyledons_odb12 (n = 2,805). Genome mode was applied to each assembly and protein mode to the corresponding proteome FASTA; the number of protein sequences evaluated is reported.

### Chromosome correspondence reveals conserved and multicomponent relationships

Synteny blocks shared among the three *Taraxacum* assemblies anchor 490.8 Mb, equivalent to 42.0% of the combined length of the eight OSU chromosome scaffolds. For each of seven OSU chromosomes, 94.2–96.6% of the anchored length maps to a single dominant TK1151 chromosome-scale pseudomolecule. OSU Chr4 shows the only substantial two-component relationship, with 73.3% of its 58.6-Mb anchored span mapping to TK1151 Chr5 and 23.9% to Chr8 (Figure 2A). The 27.9-Mb TK1151 ChrA0 unplaced sequence bin receives 8.8 Mb of dispersed OSU-source blocks but is not the predominant counterpart of any OSU chromosome. Thus, most of the chromosome-scale organization is conserved between the two *T. kok-saghyz* assemblies, while OSU Chr4 captures a more complex correspondence.

**Figure 2.**
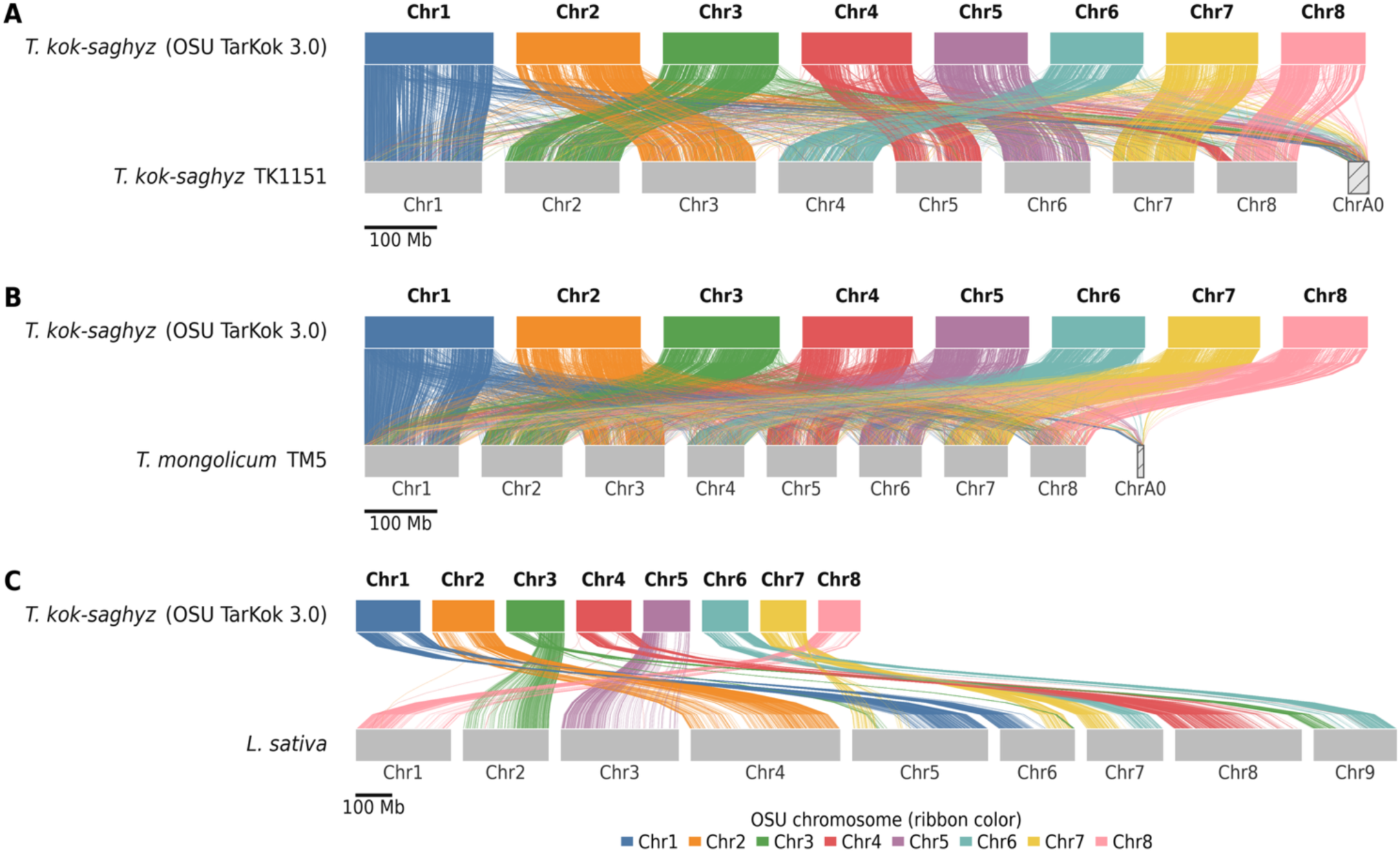
Macrosynteny of the eight OSU TarKok 3.0 chromosome scaffolds with three Asteraceae references. Ribbons show retained ntSynt blocks and are colored by OSU chromosome. Panels (A) and (B) show blocks shared among OSU TarKok 3.0, *T. kok-saghyz* TK1151, and *T. mongolicum* TM5. TK1151 and TM5 each have eight chromosome-scale pseudomolecules shown as Chr1–Chr8. Retained blocks involving ChrA0 are displayed separately as an unplaced sequence bin. Panel (C) shows the four-genome blocks also shared with lettuce (*Lactuca sativa*) and contains the nine nuclear lettuce chromosomes (Chr1–Chr9). Dominant chromosome correspondences are evident, with minor off-chromosome blocks retained. The multichromosomal pattern is compatible with rearrangement or fusion but does not identify a unique history.

The comparison with the independently assembled *T. mongolicum* TM5 genome shows similarly strong chromosome-scale conservation. Each OSU chromosome places 96.3–97.5% of its shared anchored length on one dominant TM5 chromosome-scale pseudomolecule (Figure 2B). This correspondence is notable because the *T. mongolicum* genome is substantially smaller and less extensively alignable to OSU TarKok 3.0 than TK1151. Together, the two comparisons show that predominant chromosome relationships remain recognizable across *Taraxacum* despite differences in genome size, sequence divergence, and unplaced sequence content.

The phylogenomic analysis places these structural comparisons in an evolutionary framework. A maximum-likelihood tree constructed from 1,480 complete single-copy BUSCO proteins recovered *Taraxacum*, *Cichorium*, and *Lactuca* as distinct groups, with all reported internal support values reaching 100% (Figure 3A). The two *T. kok-saghyz* assemblies formed a clade, whereas *T. officinale* grouped with *T. mongolicum*. The two *Cichorium* species formed the sister group to *Taraxacum*, and the three *Lactuca* species constituted the outgroup. Within *Lactuca*, *L. sativa* grouped with *L. saligna*, while *L. virosa* branched earlier.

**Figure 3.**
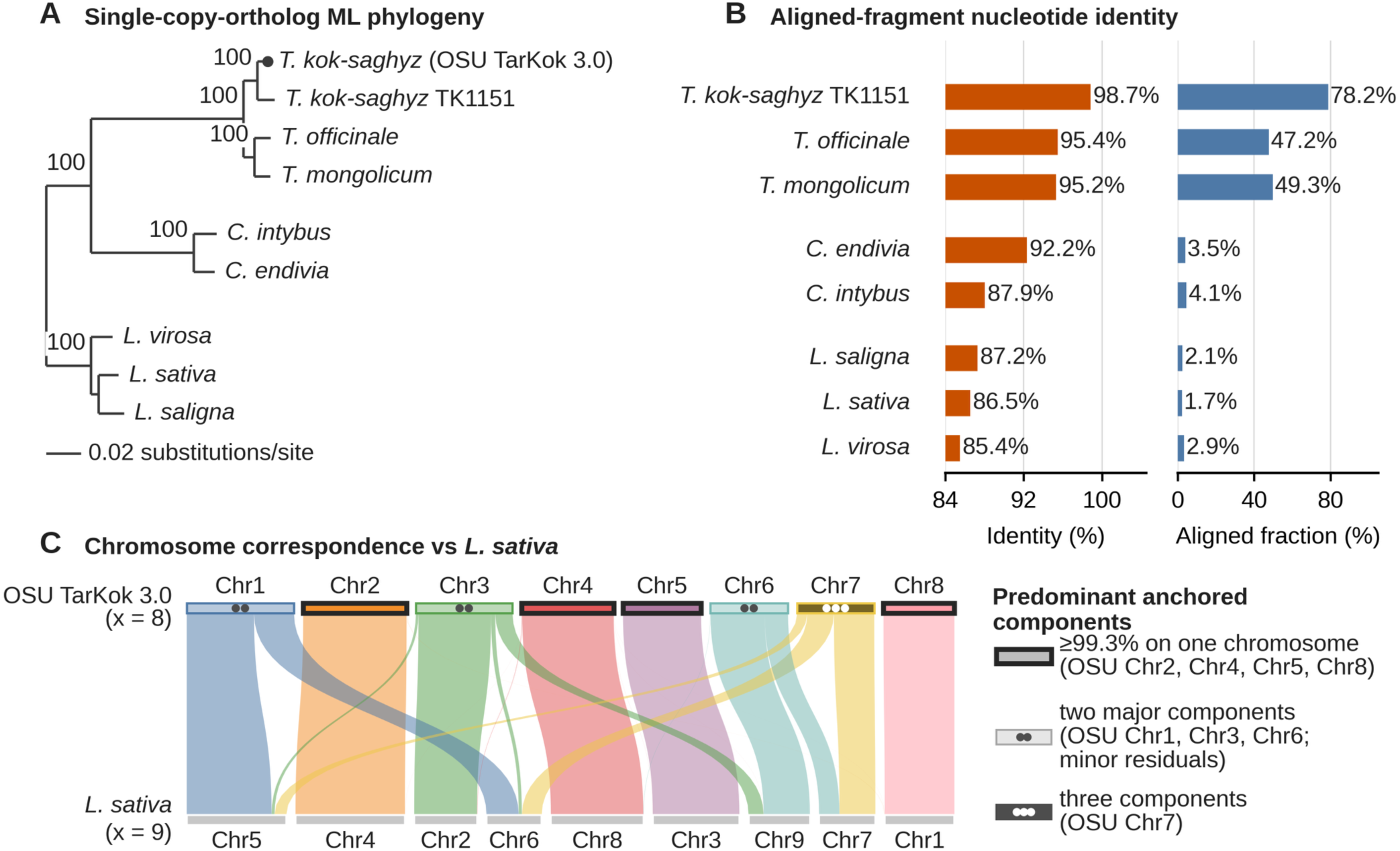
Phylogenomic placement and chromosome correspondence of *Taraxacum kok-saghyz*. (A) Maximum-likelihood phylogeny from 1,480 complete single-copy BUSCO proteins shared by nine Cichorieae genomes; branch support is shown where present in the retained tree. (B) Length-weighted nucleotide identity among best alignments retained from 5-kb comparator fragments with aligned span at least 2 kb, together with the aligned fraction of each comparator query assembly. Identity is conditional on retained alignments and is not whole-genome ANI. (C) Predominant lettuce-chromosome components of anchored syntenic length: four OSU chromosomes place at least 99.3% on one lettuce chromosome, Chr1 has two components, Chr3 and Chr6 have two major components plus minor residuals, and Chr7 has three components. These relationships do not establish the direction, mechanism, or order of chromosome-number change.

Sequence comparisons revealed a progressive decline in both identity and genomic coverage, although the two measures changed at different rates. Among retained best alignments from 5-kb comparator fragments with aligned spans of at least 2 kb, TK1151 had 98.74% length-weighted nucleotide identity across 78.20% of its query assembly. Identity remained similar for *T. officinale* and *T. mongolicum* (95.36% and 95.22%, respectively) but their aligned fractions declined to 47.17% and 49.28% (Figure 3B; Supplemental Figure S6; Supplemental Table S4). Outside *Taraxacum*, retained identity ranged from 87.9% to 92.2% for *Cichorium* and from 85.4% to 87.2% for *Lactuca*, while aligned fractions fell to only 3.5–4.1% and 1.7–2.9%, respectively. The sharp reduction in aligned fraction shows that the identity estimates for these more distant genera describe small retained portions of their genomes rather than genome-wide nucleotide similarity.

The lettuce comparison places the otherwise conserved *Taraxacum* chromosome architecture in the context of descending dysploidy. Synteny retained across all four genomes anchors 222.3 Mb, equivalent to 19.0% of the combined length of the eight OSU chromosome scaffolds. Four OSU chromosomes show nearly exclusive correspondence with one lettuce chromosome: Chr2 maps predominantly to lettuce Chr4, Chr4 to lettuce Chr8, Chr5 to lettuce Chr3, and Chr8 to lettuce Chr1, with at least 99.3% of their anchored length assigned to those respective chromosomes (Figures 2C and 3C; Supplemental Table S5). The remaining chromosomes have multicomponent relationships. OSU Chr1 comprises contributions from lettuce Chr5 and Chr6 (62.7% and 37.3%); Chr3 maps primarily to Chr2 and Chr9 (75.1% and 17.9%), with smaller contributions from Chr6 and Chr5; Chr6 maps principally to Chr9 and Chr7 (69.6% and 30.1%); and Chr7 contains contributions from Chr7, Chr6, and Chr5 (52.4%, 34.5%, and 13.2%). Across the eight OSU chromosomes, the predominant lettuce component accounts for an average of 82.4% of anchored length, ranging from 52.4% for Chr7 to 100% for Chr5 and Chr8.

This combination of conserved and multicomponent relationships provides a chromosome-level signature compatible with descending dysploidy from an x = 9-like ancestral organization to the x = 8 *Taraxacum* karyotype. The four nearly single-chromosome correspondences indicate that substantial chromosome content remained intact, whereas the composite structures of Chr1, Chr3, Chr6, and Chr7 identify where chromosome-number reduction and subsequent rearrangement may have redistributed ancestral segments. The pattern argues against treating the transition as a single, simple end-to-end fusion: fusion accompanied by translocation (Mandáková & Lysak, 2018), or a more complex series of rearrangements, would better accommodate the participation of several chromosomes. Interstitial telomeric arrays on OSU Chr1 and Chr5 provide additional structural landmarks, but only Chr1 also has a multicomponent lettuce relationship. Consequently, the telomeric and syntenic patterns together support a role for chromosome fusion in the x = 9-to-x = 8 transition without identifying a unique breakpoint, event order, or mechanism.

### Rubber-pathway family sizes overlap across Cichorieae, while focal loci show coordinated expression

We evaluated two alternatives: lineage-specific enlargement of rubber-related gene families and coordinated expression of a broadly shared gene set. If rubber production depended on lineage-specific gene dosage, the surveyed families should be enlarged in *T. kok-saghyz* relative to non-rubber-producing Cichorieae under uniform criteria; if it depended on coordinated deployment, family sizes should overlap while focal loci show tissue-restricted and coordinated expression.

The curated nuclear inventory contains six cis-prenyltransferase (CPT) loci, two (CPTL/RTA) genes, seven small rubber particle protein (SRPP) loci, one rubber elongation factor (REF) locus, five citrate-binding protein (CitBP) loci, and 15 3-hydroxy-3-methylglutaryl-CoA reductase (HMGR) loci. Full-length protein identity supports *g5630* as *TkCPTL1*/*TbRTA*, while *g9503* remains a probable divergent CPTL/RTA paralog. The structural prediction at *g12582* contains both *TkCPT1*-like and *TkCPT2*-like isoforms. Gene counts and expression for this prediction therefore describe the combined *g12582* locus and cannot be assigned separately to the two closely related paralogs. The rubber-pathway genes are distributed across the genome rather than organized as a single biosynthetic cluster. plantiSMASH identified 42 candidate biosynthetic gene clusters comprising 686 genes, including terpene and terpene-saccharide clusters. Locus-level expression values and differential-expression statistics are provided in Supplemental Table S6. Only one rubber-related locus, *g12582,* overlapped a predicted cluster, and that overlap occurred within a lignan cluster rather than a coordinated rubber-biosynthesis region. The remaining rubber-pathway loci were dispersed, and the SRPP and REF rubber-particle genes were not recovered in any predicted cluster (Supplemental Figure S7; Supplemental Table S7).

Uniform protein-to-genome projections across nine Cichorieae assemblies showed that none of the six surveyed families was uniquely enlarged in the OSU TarKok 3.0 assembly. The projected CPT count was higher in each of the three lettuce assemblies than in *T. kok-saghyz*, *Cichorium intybus* had the largest SRPP count, and *T. officinale* had the largest projected HMGR count. REF was represented by one projected homology cluster in each of the nine assemblies, while CPTL/RTA and CitBP counts varied only within narrow ranges. For HMGR, the projections recovered 15 homology clusters in OSU TarKok 3.0, compared with 12 in TK1151, 13 in *T. mongolicum*, 18 in *T. officinale*, 16 in each *Cichorium* assembly, and seven in each lettuce assembly. These broad projections differ from the curated expression inventory because they were designed to apply the same homology criteria to every genome (Supplemental Figure S8; Supplemental Table S8, Panel A).

The gene-family model reached the same conclusion for the focal HMGR test. The selected single-rate, no-error model in Computational Analysis of gene Family Evolution 5 (CAFE5) analysis tested 24,025 families, of which 891 remained significant after family-level Benjamini-Hochberg (BH) correction. HMGR-containing OG0000471 showed nominal evidence of family-size change, with raw P = 0.008 and an inferred terminal-branch change of +4 in OSU TarKok 3.0. The result did not remain significant after correction, however, because its family-level BH q value was 0.160. Thus, neither the cross-genome projections nor the corrected CAFE5 test supports a significant HMGR expansion specific to the OSU TarKok 3.0 lineage (Supplemental Figure S9; Supplemental Table S8, Panel B).

In contrast to the broad overlap in projected family sizes across genomes, focal rubber-pathway loci showed marked differences in expression among tissues. The 12-library experiment compared underground tissues comprising crown and root (n = 4) with aerial tissues comprising leaf, flower, and pooled floral and seed samples (n = 8). The combined *g12582* CPT-family locus was enriched in unde ground tissues, with an underground-to-aerial ratio of 6.95 and an adjusted P value of 0.00971. *TkCPTL1*-supported *g5630* also met the underground-enrichment criterion (underground-to-aerial ratio 5.32; adjusted P = 0.0244), as did CPT *g21319* (1.31; 0.0128) and HMGR *g12555* (3.23; 0.0133). Three adjacent SRPP loci showed still stronger enrichment: *g260* had a ratio of 8.93, *g261* a ratio of 42.20, and *g262* a ratio of 66.59. All three passed the prespecified expression-ratio and adjusted-P criteria, with adjusted P values of 9.7 × 10−4, 9.6 × 10−8, and 0.0274, respectively. By contrast, *g259* had an underground-to-aerial ratio of 3.02 but did not pass the multiple-testing threshold because its adjusted P value was 0.171 (Figure 4; Supplemental Table S6). Other CPT and HMGR loci showed heterogeneous expression across tissues, indicating that family membership alone does not predict underground enrichment. Because these measurements came from bulk crown and root samples, they establish enrichment in underground tissues but do not distinguish root cell types or demonstrate laticifer specificity.

**Figure 4.**
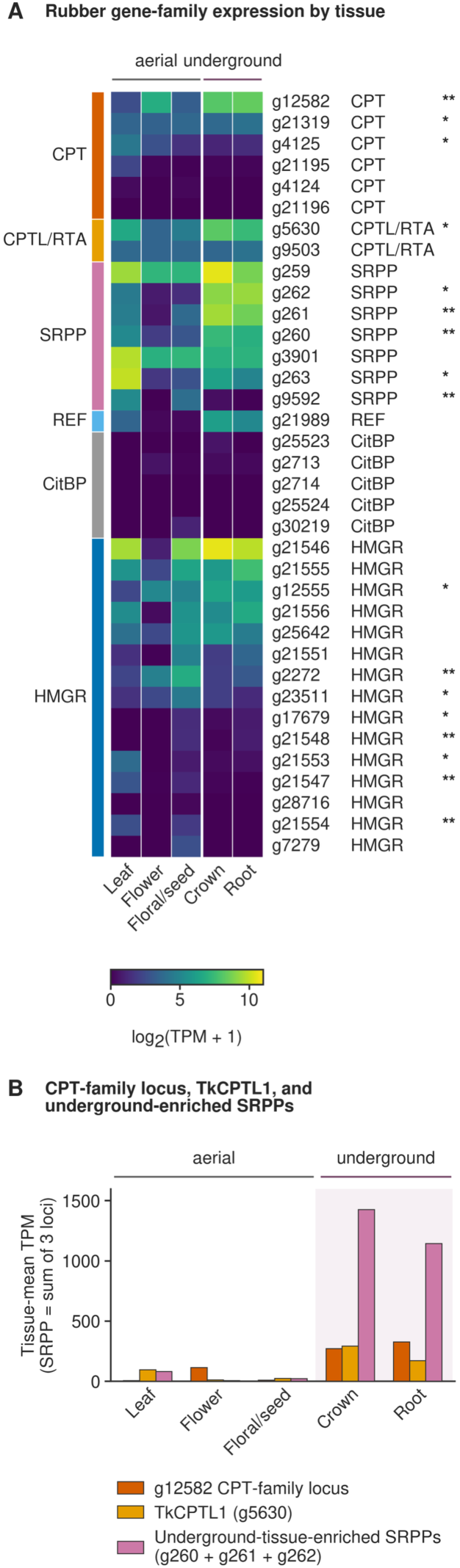
Rubber-pathway expression in the 12-library in-house dataset. (A) Locus-level expression, shown as log₂(TPM + 1), across five tissue categories. Asterisks indicate significance in the underground-versus-aerial DESeq2 contrast (* adjusted P < 0.05; ** adjusted P < 0.01). The underground group comprised crown and root libraries, whereas the aerial group comprised leaf, flower, and pooled floral/seed libraries. Expression of *g12582* is reported only at the locus level because its structural prediction contains *TkCPT1*-like and *TkCPT2*-like isoforms. (B) Expression profiles of *g12582*, *TkCPTL1*, and the summed TPM of the three underground-enriched SRPP loci *g260*, *g261*, and *g262*. *g259* was excluded from the SRPP sum because it did not meet the adjusted P < 0.05 criterion.

The broader 36-library analysis connected these tissue-level patterns to latex-containing samples. The network contained 37 biological modules evaluated across seven mutually exclusive tissue categories. The turquoise module was strongly associated with latex after correction across all 259 module-tissue tests (r = 0.9185, q = 3.81 × 10^−13^). It contained all six focal rubber-pathway loci displayed in the network: the *g12582* CPT-family locus, *TkCPTL1 g5630*, SRPP genes *g260*, *g261*, and *g262*, and REF gene *g21989* (Figure 5A). Their placement in one module links the CPT and CPTL/RTA components of the rubber-transferase complex with the rubber-particle proteins identified in the underground-tissue comparison.

**Figure 5.**
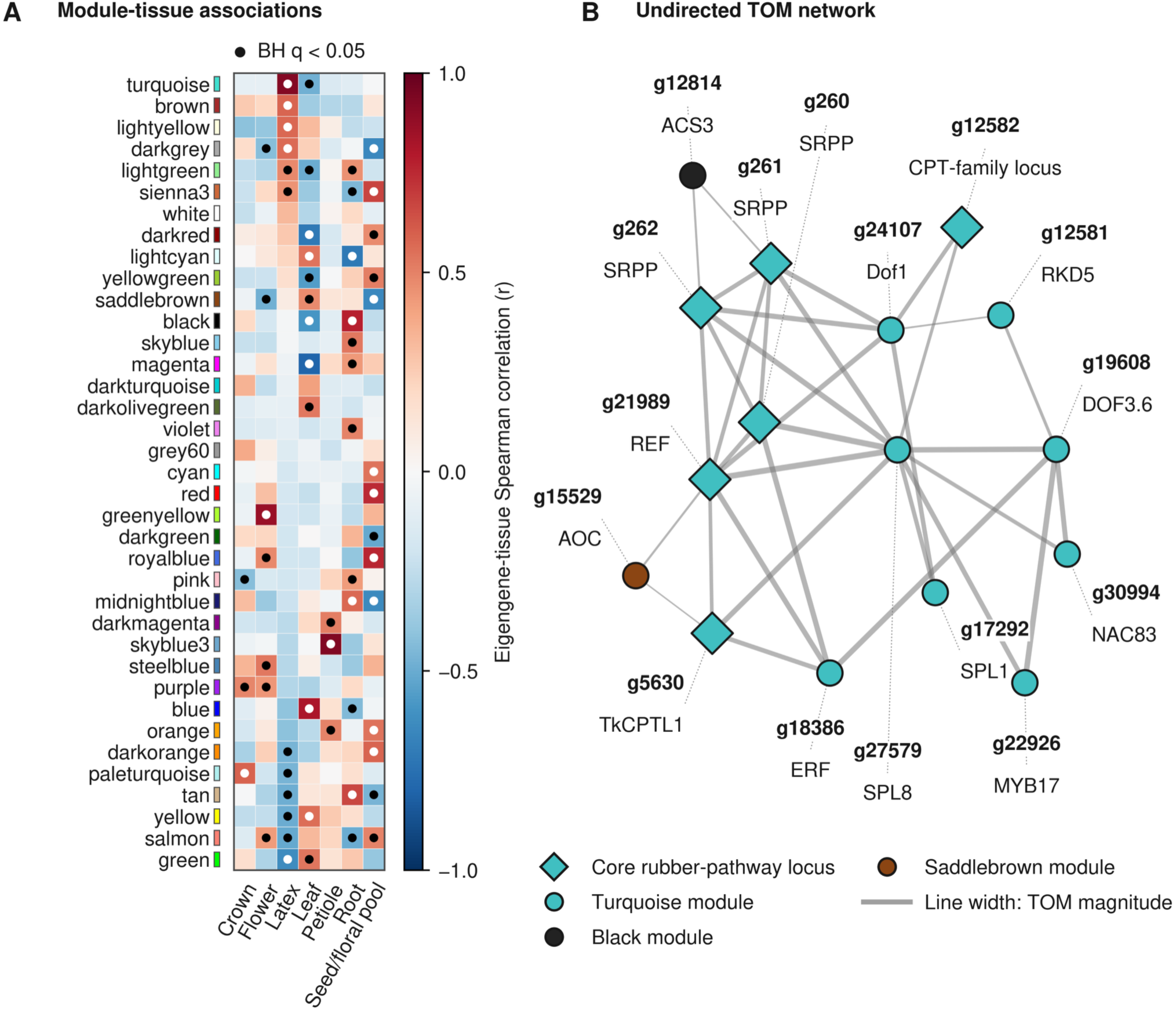
Rubber-pathway co-expression in the combined 36-library network. (A) Module–tissue correlations for 37 biological modules and seven mutually exclusive categories: Crown (n=2), Flower (n=2), Latex (n=6), Leaf (n=8), Petiole (n=3), Root (n=11), and Seed/floral pool (n=4). Asterisks denote BH adjusted q < 0.05 across all 259 module–tissue tests. The turquoise module contains the six displayed rubber-pathway loci and is latex-associated. (B) Fixed 16-node, 34-edge undirected topological-overlap network from the same 36-library analysis. Node colors are 36-library module assignments. Edges are the two strongest topological overlap measure (TOM) links per displayed node plus core-locus pairs with TOM ≥ 0.27; *g12582* and *g261* are not directly connected under this rule. Edges encode neither direction nor causality.

The filtered network view connected these six loci with transcription factors from the ERF, MYB, DOF, NAC, and SPL families, as well as genes associated with jasmonate and ethylene biosynthesis (Figure 5B). Seven prespecified promoter motifs showed no significant enrichment, and the broader JASPAR analysis likewise recovered no enriched promoter motif. Significant adjusted P values among the prespecified motifs reflected depletion rather than enrichment (Supplemental Figure S10). These transcription factors and hormone-associated genes therefore remain candidates based on co-expression rather than demonstrated regulator-target relationships. The separately fitted 12-library network served as a sensitivity analysis; all module assignments, tissue correlations, and network edges reported in Figure 5 come from the primary 36-library analysis.

All six latex libraries came from public datasets, making latex status confounded with study, genotype, and protocol. The module association cannot separate these effects completely or establish regulatory direction. Even with that limitation, three lines of evidence converge on the same working model: rubber-pathway families are not uniquely enlarged in *T. kok-saghyz*, several focal loci are preferentially expressed in underground tissues, and the central transferase and rubber-particle genes are coordinately expressed in a latex-associated module. Together, these results support coordinated pathway deployment, rather than species-specific family expansion, as the working hypothesis for rubber production in this crop.

### The mappable genome defines the accessible space for genotyping and association

In this repeat-rich genome, reliable genotyping required separating allelic variation from reads that could map to related copies. Across the eight chromosome-scale scaffolds, which span 1,168,509,718 bp, GenMap classified 573,113,065 bp (49%) as uniquely mappable at k = 150 with two allowed errors. RepeatMasker intervals covered 900,614,544 bp (77.1%) of the same chromosome span, leaving 267,895,174 bp of nonrepeat sequence. Intersecting the uniquely mappable and nonrepeat sets defined a conservative mappable genome of 210,385,031 bp, equivalent to 18% of Chr1–Chr8 and 17.6% of the complete nuclear assembly. The uniquely mappable fraction varied modestly among chromosomes, from 46.9% on Chr6 to 50.7% on Chr5. The conservative mappable fraction was 18.3, 18.0, 19.9, 16.6, 16.8, 17.9, 19.0, and 17.0% for Chr1 through Chr8, respectively, corresponding to 20.1 to 32.7 Mb per chromosome. This intersection contained 78.5% of the nonrepeat sequence but only 36.7% of the uniquely mappable sequence; 362.7 Mb of uniquely mappable sequence overlapped RepeatMasker annotations. The difference reflects the distinct criteria of the two methods: GenMap measures whether a 150-bp sequence can be placed uniquely, whereas RepeatMasker classifies sequence by similarity to repeat families. We therefore treat the GenMap set as a permissive boundary and its intersection with nonrepeat sequence as the conservative boundary used for allele-level analyses (Supplemental Table S10, Panel A).

Gene accessibility was evaluated for the 34,610 protein-coding genes located on Chr1–Chr8; the 27 genes on unplaced nuclear sequences were excluded from this chromosome-only analysis. After CDS segments were merged across isoforms within each gene, 32,117 genes (92.8%) had at least one CDS base within the conservative mappable genome, 28,003 (80.9%) had at least 50% of their CDS accessible, and 14,302 (41.3%) had all CDS bases accessible. Conversely, 2,493 genes (7.2%) had no CDS overlap with the conservative mappable genome. Across the chromosomes, 31.2 of 37.0 Mb of nonredundant CDS sequence, or 84.3%, fell within the conservative mappable genome. Coding sequence was therefore enriched 4.68-fold relative to the 18.0% chromosome-wide coverage of this space (Supplemental Table S10, Panel B). Thus, the conservative mask excludes most of the repetitive genome while retaining most annotated coding sequence.

Across 186 samples with complete consensus genotypes, caller agreement followed by minor-allele-frequency and missingness filters retained 2,076,193 biallelic allele records. Paralog filtering reduced this set to 1,117,924 records at 1,109,407 unique positions. Restricting the data to the 181-member analysis panel and reapplying the sample and variant filters retained 1,099,707 allele records, equivalent to approximately one association allele record per 191 bp of conservative mappable sequence (Table 2).

**Table 2.** Cohorts and marker sets used for population association and F2 mapping. Counts correspond to successive analysis stages. Allele records can exceed unique positions after multiallelic decomposition. For population association, cross-model support required the same allele record to pass its model-specific genome-wide threshold in at least two of three models. LOD is the full-minus-null log likelihood divided by ln(10); thresholds are the 95th percentiles of 1,000 permutation maxima.

| Analysis stage | Samples | Phenotype classes | Records/markers and result |
| --- | --- | --- | --- |
| Sequenced / complete consensus | 187 / 186 | Not applicable | Sample flow |
| Consensus after minor-allele-frequency and missingness filtering | 186 | Not applicable | 2,076,193 biallelic allele records |
| Paralog-filtered consensus | 186 | Not applicable | 1,117,924 allele records at 1,109,407 unique positions |
| Association panel | 181 | Endpoint-specific subsets below | 1,099,707 allele records tested in association models |
| Filled-seed population endpoint | 166 | 84 filled seed / 82 no filled seed; 15 unscored | 92,651 LD-pruned markers for $F_{ST}$; no genome-wide significant marker in any model |
| Herbicide population endpoint | 177 | 145 tolerant / 32 susceptible; 4 unscored | Two genome-wide significant markers in GEMMA, none in GCTA or PLINK2, and no exact marker supported by at least two models |
| $F_2$ filled-seed scan | 83 | 48 filled seed/ 35 no filled seed | Chr1 LOD 4.59 > 4.42 |
| $F_2$ herbicide scan | 83 | 53 tolerant / 30 susceptible | Chr2 LOD 3.08 < 4.53 |
| Full linkage map | 83 | Not applicable | 7,624 ParentCall2 markers 4,474 ordered markers; 621.75 cM |
| Marey/recombination subset | 83 | Not applicable | 4,466 markers; 619.34 cM; 528 rate-bearing windows and 61 unavailable windows |

The population analysis first examined the filled-seed endpoint as an operational proxy for selfing capacity. Plants were classified according to whether bagged capitula produced filled seed without a directed cross. This screen is relevant to self-compatibility, although it does not establish the reproductive mechanism. Among the 181 panel members, 166 were scored: 84 produced filled seed and 82 did not, while 15 lacked a score. The two phenotypic groups largely overlapped in the PCA, which included all 181 individuals. Across the 166 scored plants, 92,651 LD-pruned single-copy markers produced a mean per-marker FST of 0.009645 and a maximum of 0.325581 (Supplemental Figure S11), consistent with little genome-wide differentiation although individual-marker estimates reached 0.325581. The herbicide analysis included 177 scored individuals, comprising 145 tolerant and 32 susceptible plants; this imbalance reflects survival selection before genotyping. No marker reached genome-wide significance in any of the three filled-seed models. For herbicide response, GEMMA identified two markers above its Bonferroni threshold, led by Chr6:124,035,144 (P = 6.76 × 10⁻⁹), but neither marker passed its corresponding threshold in GCTA or PLINK2. Consequently, no exact allele record passed its model-specific threshold in at least two models for either endpoint. Model-specific marker counts, thresholds, genomic-inflation factors, and leading markers are reported in Supplemental Table S11, with Manhattan and quantile-quantile plots in Supplemental Figure S14. Thus, population association did not identify a cross-model-supported locus for either endpoint in this related panel.

### F2 mapping detects a chromosome 1 QTL for selfing-related seed production

Filled-seed production from bagged capitula was used as a practical proxy for selfing ability in 83 independent F2 plants. Forty-eight plants produced filled seed, whereas 35 did not. The scan detected a QTL at Chr1:80,117,130 with LOD 4.59, exceeding the experiment-wise permutation threshold of 4.42. Only the peak marker formed the connected component above the peak-minus-1.5-LOD cutoff, so the interval boundaries were estimated by interpolation to the nearest flanking markers below the cutoff. This placed the support interval at Chr1:80,072,004 to 80,146,944, a span of 74.94 kb. The interval overlaps a single predicted protein-coding gene, *g7174*, which encodes a chloroplast-targeted phospholipase A1-like protein with a class 3 lipase domain. The peak marker is intergenic, 43.9 kb downstream of *g7174* and 77.2 kb upstream of *g7175*. Other members of the broader phospholipase A1 and class 3 lipase families contribute to jasmonate-dependent pollen maturation, anther dehiscence, and floral development in other plant species, making *g7174* a positional candidate for functional testing (Ishiguro et al., 2001; Li et al., 2009; Cai et al., 2014). In the same F2 population, the herbicide-response scan included 53 tolerant and 30 susceptible plants. Its maximum LOD of 3.08 at Chr2:121,982,779 remained below the permutation threshold of 4.53, and no significant herbicide-response QTL was detected (Supplemental Figure S12A,B; Supplemental Table S9).

The F2 genotype set also produced a chromosome-scale linkage map. Marker selection began with 10,737 biallelic allele records at 10,617 unique founder-heterozygous positions. ParentCall2 retained 7,624 markers, and 4,474 ordered markers formed nine linkage groups spanning 621.75 cM. These groups represented the eight physical chromosomes, with Chr1 divided between LG8 and LG9 (Supplemental Figure S12C). Removing eight markers assigned outside their dominant physical chromosomes produced a chromosome-dominant Marey subset of 4,466 markers spanning 619.34 cM, equivalent to a genome-wide average of 0.530 cM/Mb.

Recombination rates were estimated in 528 of 589 nonoverlapping 2-Mb windows; the remaining 61 lacked marker-span coverage and were retained as unavailable rather than assigned a zero rate. Repeat fraction and recombination rate showed a descriptive inverse relationship across the 528 rate-bearing windows (Spearman ρ = -0.6227), consistent with reduced recombination in repeat-rich regions (Supplemental Figure S13). The chromosome 1 QTL therefore narrows follow-up to *g7174* and the adjacent intergenic region. Because the support interval was interpolated around a single connected peak marker, validation in larger families, together with allele and reproductive-tissue expression analyses, will be needed to determine whether this locus influences pollen function, fertilization, or subsequent seed development.

## Discussion

The improved continuity of OSU TarKok 3.0 provides a common framework for examining chromosome reorganization, rubber-pathway regulation, and domestication traits in a large, repeat-rich genome. The comparative analyses reveal strong chromosome conservation within *Taraxacum* and multicomponent relationships with lettuce, while the copy-number and expression results support coordinated pathway deployment rather than unique expansion of rubber-related gene families. The same reference defines the short-read mappable space, supports chromosome-scale linkage analysis, and has enabled identification of a chromosome 1 QTL for selfing-related filled-seed production. Together, these results turn previously disconnected observations into specific hypotheses that can be tested across accessions, tissues, and breeding populations.

### Within-chromosome contiguity expands the analytical value of the reference

The principal advance of OSU TarKok 3.0 is continuity within chromosome scaffolds rather than chromosome-scale organization alone. Scaffold N50 differs only modestly from TK1151 because both references are organized at chromosome scale, but internal continuity differs by almost 30-fold: contig N50 rises from 1.3 to 37.8 Mb, the largest contig from 6.0 to 117.7 Mb, and the number of gaps falls from 1,917 to 88. Because repetitive sequence occupies 77.5% of the assembly, the value of these improvements extends beyond the assembly statistics themselves. Longer uninterrupted sequence preserves gene neighborhoods and syntenic blocks, reduces interruptions within candidate intervals, provides greater context for resolving recent paralogs, and makes local mappability measurable along chromosomes. These properties are especially important when short reads from related genomic copies can be mistaken for allelic variation.

The additional sequence in OSU TarKok 3.0 is also informative. The nuclear assembly is 91.1 Mb longer than TK1151 and approximately 112 Mb larger than the GenomeScope2 estimate. Several observations are consistent with recovery of repeat-rich sequence as an important contributor to this difference: the k-mer spectrum contains an extensive high-copy tail beyond the fitted region, repeats occupy 77.5% of the primary assembly, and the length-weighted LTR Assembly Index (LAI) is 41.6. The first rubber dandelion genome reported evidence that recent LTR retrotransposon insertions contributed substantially to genome enlargement in this species (Lin et al., 2018). LAI is informative because it evaluates assembly continuity using LTR retrotransposons, which are among the most difficult regions of a plant genome to assemble, rather than gene-space completeness alone (Ou et al., 2018). It does not, however, measure the amount of repetitive sequence recovered. The contributions of biological length variation, repeat recovery, and assembly behavior cannot be partitioned from these comparisons, but the repeat profile makes improved recovery of high-copy sequence a plausible explanation for much of the apparent excess. The final nuclear gene count provides no indication that the difference instead reflects an unusual increase in gene number.

The representative proteome extends that improvement into gene-level analysis. Protein-mode BUSCO completeness rises from 73.5% in the TK1151 proteome to 97.5% in OSU TarKok 3.0, while duplicated BUSCOs fall from 28.2% to 5.4%. This result does not validate every gene model, but provides a much more complete and less redundant one-protein-per-gene set for orthology and family-size inference. That distinction matters because fragmented, fused, or duplicated predictions can create artificial gains and losses before any evolutionary model is fitted. At chromosome scale, complementary evidence supports the organization of the reference: five chromosomes carry telomeric arrays at both ends, all eight show broadly uniform HiFi depth, and chromosome correspondence is strong against an independently assembled congener. OSU TarKok 3.0 therefore provides a stronger reference for constructing a species pangenome, surveying structural variation, resolving recent paralogs, and developing reliable markers.

### The mappable genome defines the reach and limits of short-read genotyping

The repeat structure that complicates assembly also determines which variants short reads can resolve, and continuity is what makes that boundary measurable. GenMap classified 573.1 Mb, or 49% of the combined length of the eight-chromosome span as uniquely mappable, setting a permissive boundary based on sequence mappability alone. Intersecting that sequence with nonrepeat regions produced a more conservative mappable genome of 210 Mb, about 18% of the chromosome span, which was used for allele-level analyses. The 362.7-Mb difference consists of sequence that can be mapped uniquely under the GenMap parameters but is nevertheless classified as repetitive. The two filters therefore measure different properties: GenMap evaluates whether a 150-bp sequence can be placed uniquely, whereas RepeatMasker identifies similarity to known repeat families. Requiring both unique mappability and nonrepeat status provides a conservative safeguard against treating reads from related genomic copies as allelic variation, turning a normally hidden ascertainment bias into an explicit boundary for analysis.

Within the conservative mappable genome, the filtered marker set supports population analysis, linkage mapping, and targeted assay design with reduced risk of treating paralogous reads as alleles. The retained association dataset contains an average of one fitted allele record per 191 bp of mappable sequence. Regions outside this space are repetitive, nonunique, or both, and were not interrogated reliably by the present short-read design. The consequence for interpretation is direct and relevant to other repeat-rich emerging crops. The absence of a population-association signal shared by at least two models means that no locus received the prespecified cross-model support within the conservative mappable fraction of this related panel. It does not indicate that filled-seed production or herbicide response lacks genetic control. This explicitly bounded result identifies the next design priorities: larger and less related panels for the short-read mappable space, together with long-read genotyping and a multi-accession pangenome for duplicated and structurally variable regions (Della Coletta et al., 2021).

Making the mappable genome explicit also has an immediate practical use. Short-read marker and assay development can now prioritize regions where allelic interpretation is defensible, and the fraction of any candidate region falling inside or outside that boundary can be stated in advance rather than discovered after genotyping. For a species in which paralogy has repeatedly complicated marker work, these boundaries are a durable contribution independent of any single locus reported here.

### Conserved and composite chromosomes refine the descending-dysploidy hypothesis

Chromosome organization is remarkably stable within *Taraxacum*. Seven OSU chromosomes place 94.2 to 96.6% of their anchored length on one TK1151 pseudomolecule, and every OSU chromosome places 96.3 to 97.5% on one dominant *T. mongolicum* pseudomolecule. This conservation persists even as retained nucleotide identity and, more sharply, the fraction of aligning sequence declines outside the conspecific comparison. Large chromosome segments remain recognizable despite substantial sequence divergence, which is consistent with the divergence reported among *Taraxacum* genomes (Lin et al., 2022). OSU Chr4 is the principal exception, with substantial relationships with two TK1151 pseudomolecules. Chr4 showed no corresponding chromosome-scale anomaly in the available assembly checks: HiFi depth was broadly uniform, its uniquely mappable fraction was 48.3% (within the 46.9 to 50.7% range across the eight chromosomes), and its Marey curve showed no major discontinuity in physical-genetic order. Like every OSU chromosome, Chr4 also retained one dominant correspondence with *T. mongolicum*. These observations provide no evidence of a gross assembly error, although they cannot distinguish accession-specific structural variation from a localized difference between the two assemblies.

The lettuce comparison places this conserved framework in the context of chromosome-number change. Chr2, Chr4, Chr5, and Chr8 retain nearly exclusive correspondence with one lettuce chromosome, whereas Chr1, Chr3, Chr6, and Chr7 combine two or three major lettuce-affinity components. Within the anchored subset this mixture of predominantly single-component and multicomponent relationships is compatible with descending dysploidy from an x = 9-like organization to the x = 8 *Taraxacum* karyotype. This pattern is also compatible with a history more complex than single unreworked end-to-end fusion, potentially involving fusion together with translocation and other rearrangements. Cytogenetic and phylogenomic studies in *Reichardia* support x = 9 as a plausible ancestral condition in Cichorieae and show that dysploidy can involve changes in heterochromatin as well as chromosome organization (Siljak-Yakovlev et al., 2017; Cho et al., 2022). This interpretation also fits the deeper history of Asteraceae, in which paleohexaploidy, genomic exchange, fractionation, and repeated chromosome reshuffling produced mosaic modern genomes (Feng et al., 2026).

The two interstitial telomeric arrays add landmarks without resolving the history. One lies on composite Chr1, the other on Chr5, which has nearly exclusive lettuce Chr3 correspondence. Their different contexts make them candidates for retained, relocated, or independently generated telomeric sequence rather than equivalent evidence for one fusion event. The four-genome blocks anchor 19.0% of the OSU chromosome span, sufficient to identify predominant chromosome affinities within the retained blocks but insufficient to determine the direction, order, breakpoints, or mechanism of chromosome-number change. What the comparison achieves is a narrowing: a broad chromosome-count hypothesis becomes an explicit division into four largely conserved and four composite chromosomes, against which chromosome-contiguous sampling across Cichorieae, cytogenetic localization, and breakpoint-scale alignment can now test alternative histories.

### Rubber-pathway organization supports coordinated deployment rather than lineage-specific expansion

The comparative analyses distinguish two explanations for rubber production: lineage-specific expansion of pathway families and coordinated use of a broadly conserved toolkit. Uniform projections across nine Cichorieae genomes showed substantial overlap in all six surveyed families. CPT counts were higher in the three lettuce genome assemblies, *Cichorium intybus* had the largest projected SRPP complement, and

*T. officinale* had the largest projected HMGR count. The HMGR-containing orthogroup likewise did not remain significant after correction across the 24,025 CAFE5 tests. Earlier pairwise comparison of TK1151 with *T. mongolicum* identified relative expansion of CPT and SRPP families in the rubber-producing genome (Lin et al., 2022). The broader sampling here refines rather than contradicts that observation, since a family can be larger than in one comparator without being uniquely enlarged across the tribe. Copy number may still contribute to quantitative variation, but it does not provide a sufficient general explanation for rubber production across the surveyed taxa.

Genome organization and expression instead converge on pathway deployment. The curated rubber-related genes are distributed across the chromosomes, and plantiSMASH did not recover them as one coordinated biosynthetic cluster. Despite that physical dispersion, the *g12582 CPT1*/*CPT2*-like locus and *TkCPTL1*-supported *g5630* were enriched in underground tissues together with CPT *g21319* and HMGR *g12555*, and the adjacent SRPP loci *g260*, *g261*, and *g262* showed underground-to-aerial expression ratios from 8.93 to 66.59. Five of these underground-enriched loci, *g12582*, *g5630*, *g260*, *g261*, and *g262*, together with REF *g21989*, were the six displayed rubber-pathway loci in the turquoise module of the 36-library network. The module eigengene was strongly associated with latex samples (r = 0.9185, q = 3.81 × 10^−13^). A local SRPP cluster therefore participates in a broader co-expression pattern linking genes involved in rubber-chain elongation with rubber-particle structure across dispersed loci.

Independent experiments make such regulation plausible and give the pattern mechanistic context. *TkCPT1* and *TkCPT2* form complexes with *TkCPTL1* that produce extra-long-chain polyisoprenes (Müller et al., 2025); CRISPR disruption of *TkCPT1* reduces rubber production and joint disruption of *TkCPT1* and *TkCPT2* blocks rubber biosynthesis and rubber-particle formation (Xu et al., 2025); SRPP paralogs differ in expression and interaction partners (Wolters et al., 2024); *TkMYC2* overexpression increases rubber production and SRPP/REF expression (Wu et al., 2024); *TkMYC2* binds the promoters of *CPT2* and several SRPP genes (Fantini et al., 2025); and *TkDof20* directly activates *TkSRPP7*, increasing laticifer area and rubber accumulation (Fang et al., 2026). The underground-tissue contrast was based on bulk crown and root samples and therefore does not establish root or laticifer specificity. The present network did not recover significant promoter-motif enrichment, and its latex libraries are confounded with study, genotype, and protocol, so it does not identify the regulator of these loci. What it does provide is convergence across family size, genomic distribution, tissue enrichment, co-expression, and independent functional biology. That convergence makes coordinated deployment of a dispersed pathway the strongest working model produced by this study.

For germplasm development, the practical implication is to evaluate regulatory variation and expression differences alongside gene copy number. Expression variation among accessions and transcription factors co-expressed with the focal loci are candidates for functional testing, but the present data do not establish that they predict rubber yield. Because the *g12582* prediction combines *CPT1*-like and *CPT2*-like isoforms, targeted long-read transcripts and paralog-specific assays remain necessary before the two components can be separated in this accession.

### The recombination landscape structures where mapping resolution is achievable

The F2 marker set produced a genetic map with 4,474 ordered loci spanning 621.75 cM. Most physical chromosomes were represented by one dominant linkage group, while Chr1 remained divided between LG8 and LG9. This map complements the earlier high-density *T. kok-saghyz* map developed for rubber-content QTLs by extending trait mapping to reproductive and herbicide-response phenotypes on OSU TarKok 3.0 coordinates (Yang et al., 2023). It also reveals a structured recombination landscape. Across 528 rate-bearing 2-Mb windows, recombination rate showed an inverse association with repeat fraction (Spearman ρ = -0.6227). The association is descriptive, but is consistent with a broad flowering-plant pattern in which crossovers are concentrated toward gene-rich, repeat-poor chromosome regions (Brazier & Glémin, 2022). High-resolution analysis in soybean likewise found suppressed recombination in repeat-rich and pericentromeric regions (Ma et al., 2023). In rubber dandelion, the same heterogeneity indicates where linkage may facilitate or limit QTL resolution and the separation of favorable alleles.

### A chromosome 1 locus connects reproductive biology to a testable candidate

Within-family segregation revealed a chromosome 1 locus for filled-seed production from bagged capitula, the study’s operational endpoint for selfing-related seed production under restricted cross-pollination. The peak at Chr1:80,117,130 exceeded the experiment-wise permutation threshold, and interpolation around the single connected peak marker defined a 74.94-kb support interval whose left boundary overlaps 1,248 bp of the 3′ end of *g7174*. The peak marker itself is intergenic, 43.9 kb downstream of *g7174*. The family result complements the population analysis in a way that is instructive for outcrossing crops generally: segregation within a single F₂ reduces the broad population structure that dominates a mixed panel and can expose an allele that is rare or family-specific, which is precisely the situation expected when reproductive variation is maintained at low frequency in an outcrossing species.

The annotation suggests a specific and testable reproductive hypothesis. *g7174* encodes a predicted chloroplast-targeted phospholipase A1-like protein with a class 3 lipase domain. Arabidopsis *DAD1* is a chloroplast-localized phospholipase A1 that initiates jasmonate biosynthesis in flowers and synchronizes pollen maturation, anther dehiscence, and flower opening (Ishiguro et al., 2001), while in rice, the putative class 3 lipase *EG1* regulates spikelet development and jasmonate signaling controls floral patterning through the *OsMYC2*-*OsMADS1* pathway (Ishiguro et al., 2001; Cai et al., 2014). These analogies connect the predicted localization and domain of *g7174* to lipid signaling and reproductive development, without making it the *Taraxacum* ortholog of a characterized fertility gene. However, they also point to an important distinction.

A variant near *g7174* could affect pollen maturation, anther function, or floral development without altering the self-incompatibility recognition system itself. Rubber dandelion has a complex reproductive baseline: classical embryology established sexual development with strong summer self-sterility and increased self-fertility under some late-season conditions (Warmke, 1943). Microscopy of modern material demonstrated a sporophytic self-incompatibility response that restricts self-pollen progression (Wollenweber et al., 2021). Germplasm screening recovered variation in the degree of self-incompatibility and individuals producing selfed seed while cytological analysis confirmed predominantly regular sexual sporogenesis and gametogenesis (Ponciano et al., 2026; Schuchovski et al., 2026). In Asteraceae more broadly, sporophytic self-incompatibility involves *S*-locus dominance interactions and unlinked modifiers of cross-compatibility and selfing rate, so partial and quantitative variation in selfing is expected rather than anomalous (Hiscock et al., 2003; Brennan et al., 2011). The filled-seed endpoint cannot distinguish among these mechanisms. The next steps are to genotype progeny to confirm selfing, compare *g7174* and the surrounding intergenic sequence between the parents, assay expression and pollen behavior during controlled self-and cross-pollination, and replicate the effect in larger families. If its association with selfed seed production is validated, a locus conferring predictable selfed seed set would address one of the two central obstacles to line development in this crop.

### An informative null for herbicide response

The herbicide-response scan in the 83-plant F₂ family did not cross its permutation threshold, with a maximum LOD of 3.08 at Chr2:121,982,779 against a threshold of 4.53. We interpret this result within the limits of the design, rather than as evidence against genetic control, for three reasons. First, the endpoint records emergence and survival through the first true leaf after a sulfentrazone was applied at a single rate and therefore does not capture partial injury or dose response. Measurements across replicated doses could recover quantitative variation that the present binary score discards. Second, the scan comprised 53 tolerant and 30 susceptible F₂ plants, and this sample size limits power to detect modest genetic effects (Beavis, 1998; Xu, 2003). Because both endpoints were scored on the same 83 plants using the same genotype data, marker set, and family structure, the contrast between a detected chromosome 1 locus and the non-significant herbicide scan cannot be attributed to differences in sample composition or marker coverage. Instead, it may reflect differences in genetic architecture, effect size, phenotype resolution, or sampling variation between the two endpoints. Third, and most generally, the scan interrogated only founder-heterozygous, single-copy markers within the short-read mappable space. A causal variant in duplicated or repeat-associated sequence was not directly assayed and might not be captured through linkage to an accessible marker. Herbicide tolerance remains a priority breeding objective for this crop, because it could expand selective chemical control during establishment, which is one of the few management levers available during the period when the crop is most vulnerable to weed competition (Iaffaldano et al., 2018; Pannwitt et al., 2025), and the design requirements for detecting it are now explicit.

### Current limitations define priorities for genomics-assisted domestication

Several constraints define the scope of these conclusions. OSU TarKok 3.0 was generated from the same accession and underlying sequence data as OSU TarKok 2.0, using a different assembly pipeline. Chromosome-scale ordering used the existing Omni-C-supported framework rather than a newly generated long-range library, and a secondary reference-guided placement step assigned approximately 1% of the assembly using TK1151 as a guide. Read depth, telomeric arrays, and the independent *T. mongolicum* comparison provide complementary but not equivalent support. Expression measurements came from bulk crown and root samples, which establish enrichment in underground tissues but do not distinguish root cell types or demonstrate laticifer specificity. All latex libraries in the co-expression analysis were public and therefore confounded with study, genotype, and protocol, so the module-to-latex association requires validation in a balanced design. The association panel is related and modest in size, the F₂ family comprises 83 plants, and both trait endpoints are binary operational scores rather than mechanistic measurements. The chromosome 1 support interval was interpolated around a single connected peak marker and requires replication before it is treated as a fine-mapped assignment.

Within those bounds, the results replace several disconnected genomic observations with a connected and testable framework for improving an outcrossing crop. The reference establishes what fraction of the genome current markers can interrogate, how recombination varies with repeat content and may influence mapping resolution, which pathway loci show coordinated expression and therefore merit testing for regulatory variation alongside copy number, and where selfing-related filled-seed variation segregates within one family. The immediate priorities that follow are a multi-accession pangenome to characterize sequence diversity and structural variation outside the mappable genome, long-read genotyping of duplicated regions, larger and less related population panels for both domestication endpoints, and progeny testing to determine whether the chromosome 1 locus acts through self-compatibility, pollen function, or subsequent seed development. Rubber dandelion has been studied primarily as a rubber biosynthesis system; the framework presented here is intended to support its development as a crop.

## Conclusions

OSU TarKok 3.0 turns the repeat-rich rubber dandelion genome into a practical framework for biological discovery and crop development. Its internal continuity and improved representative proteome support chromosome comparison, gene-family analysis, expression profiling, and repeat-aware genotyping on the same coordinates. Macrosynteny preserves chromosome identity within *Taraxacum* while identifying four composite relationships with lettuce that refine a complex descending-dysploidy hypothesis. Rubber-related families are broadly shared across Cichorieae, but central transferase and rubber-particle loci converge through underground enrichment and latex-associated co-expression. Together with recent functional studies, this evidence favors coordinated deployment of a dispersed biosynthetic system over unique family expansion as the main working model for rubber production. The reference also defines approximately 210 Mb of reliable short-read genotyping space, anchors a chromosome-scale linkage and recombination map, and localizes a QTL for selfing-related seed production to a 74.94-kb interval whose left boundary overlaps *g7174*. These results replace several disconnected genomic observations with testable hypotheses about chromosome history, pathway regulation, and reproductive variation, providing a clearer path from reference assembly to rubber dandelion improvement.

## Abbreviations

BUSCO: Benchmarking Universal Single-Copy Orthologs
CitBP: citrate-binding protein
CPT: *cis*-prenyltransferase
CPTL/RTA: *cis*-prenyltransferase-like/rubber transferase activator
FST: fixation index
HiFi: high fidelity
LOD: logarithm of odds
QTL: quantitative trait locus
REF: rubber-elongation factor
SRPP: small-rubber-particle protein
TPM: transcripts per million
WGCNA: weighted gene co-expression network analysis.

## Data Availability

Sequence reads analyzed in this study are available in the NCBI Sequence Read Archive under BioProject PRJNA792143 and SRA study SRP731533. These comprise PacBio HiFi reads from OH-1004 (SRR40405429), the Omni-C reads from the same accession (SRR40405428), 12 RNA-seq libraries (SRR40404971–SRR40404982), and seqWell low-pass whole-genome genotyping reads from 187 individuals (SRR40406659–SRR40406866). The OSU TarKok 3.0 nuclear genome assembly and annotation have been submitted to GenBank under WGS accession JAJTUU000000000; processing and public release are pending. The reference-guided OH-1004 chloroplast consensus has been deposited in GenBank under accession PZ899947. Software versions, scientific parameters, and cohort definitions are provided in the Supplemental Information and the GitHub repository: https://github.com/Fresnedo-Lab/OSU-TarKok-3.0. The OSU TarKok 3.0 nuclear assembly and associated data files are available at https://www.rubber-dandelion-genomics.org and permanent Zenodo repository: https://doi.org/10.5281/zenodo.23585451. These resources include the unmasked and soft-masked nuclear genome FASTAs; BRAKER3 and UTR-augmented structural annotations; CDS, all-isoform protein, and representative-protein FASTAs; coordinates of mappable genome BED file, gene–transcript mappings; integrated and tool-specific functional annotations from InterProScan, eggNOG-mapper, iTAK, dbCAN, KofamScan, and TargetP; ncRNA, RepeatMasker, EDTA, and plantiSMASH annotations; chloroplast FASTA, GFF3 and CDS, gene, protein, rRNA, and tRNA FASTAs; and OrthoFinder orthogroup assignments, gene-count tables, single-copy ortholog lists, the species tree, and pairwise OSU–TK1151 orthology relationships, and all corresponding MD5 checksums.

## Author Contributions

P.V.M. Writing – original draft; Visualization; Formal analysis; Data curation; Writing – review & editing; Software; Investigation. C.G.M Investigation; Resources; Methodology; Writing – review. E.S.A Investigation; Resources; Methodology; Writing – review & editing. D:R:W Resources. J:V:A Resources. J.F.R. Funding acquisition; Conceptualization; Supervision; Writing – review & editing. All authors read and approved the submitted version.

## Funding

This work was supported by the Agriculture and Food Research Initiative-Foundational and Applied Science Program, project award no. 2020-67013-30876, from the U.S. Department of Agriculture’s National Institute of Food and Agriculture.

Hatch program, project award no. OHO01510, from the U.S. Department of Agriculture’s National Institute of Food and Agriculture.

U.S. NSF Engineering Research Center for the Transformation of American Rubber through Domestic Innovation for Supply Security (TARDISS): This material is based upon work supported by the U.S. National Science Foundation under Cooperative Agreement No. 2330145. Any opinions, findings and conclusions or recommendations expressed in this material are those of the author(s) and do not necessarily reflect the views of the U.S. National Science Foundation.

## Acknowledgments

The authors thank the Molecular and Cellular Imaging Center (MCIC, CFAES Analytical Services, The Ohio State University) for library preparation, sequencing, and read processing, and the Ohio Supercomputer Center for computational resources. The authors also thank Debbie Zaborski, research assistant, for her assistance in the early stages of development of the inbred line OH-1004.

## Conflict of Interest

The authors declare no competing interests.

## Declaration of Generative AI and AI-Assisted Technologies

During the preparation of this work, the authors used OpenAI Codex (GPT-5.6) to assist with language organization and reproducible document assembly. The authors reviewed and edited the output as needed to ensure accuracy and take full responsibility for the content of the published article.

## Supplemental Material

Supplemental Methods, Supplemental Figures S1–S14, and Supplemental Tables S1–S11 are provided in the separate Supplemental Information.

## Supplemental Information

### From chromosome-scale assembly to the mappable genome of rubber dandelion: chromosome reorganization, rubber-pathway coordination, and domestication traits

Pablo Vargas-Mejía, Camila Gutiérrez-Manriquez, Elizabeth S. Anderson, Daniel R. Williams, Julio Vega-Arreguín, and Jonathan Fresnedo-Ramírez

### Contents

Supplemental Methods; Supplemental Figures S1-S13; Supplemental Tables S1-S8; References.

### Supplemental Methods

These Supplemental Methods provide software versions, parameters, filters, cohort definitions, and assembly-stage definitions supporting the main text. Analyses used either the 857-sequence primary assembly before organellar filtering or the 659-sequence, nuclear subset obtained after organellar partitioning, or the final 658-sequence nuclear assembly after removal of one contaminant, as specified below. Genotype filters were designed to reduce read misassignment between paralogous regions, and such misassignment was not treated as evidence of assembly redundancy.

### Plant materials and growth conditions

The sequenced material comprised 187 *Taraxacum kok-saghyz* individuals. Complete consensus genotypes were obtained for 186 individuals., HTS_138 was the only sequenced individual without a complete consensus genotype. The 181-member final panel excluded HTS_060, HTS_097, HTS_110, Juliett, and HTS_174 because they have no row in the master trait table. The final panel contains HTS_006, 83 F2, 77 F3, and 20 non-family accessions. The QTL scans used only the 83 independent F2.

Seeds were sown in 200-cell flats filled with a 50:50 mixture of Promix FLX and field soil. Once the fifth true leaf had unfolded in more than half of the plants, seedlings were transplanted into 10 cm wide by 35 cm long tree pots filled with pre-moistened Promix BX Mycorrhizae. Plants were grown under standard greenhouse conditions at the Ohio Agricultural Research and Development Center (Wooster, OH): a day temperature setpoint of 18 to 24 degrees Celsius and a night setpoint of 14 to 17 degrees Celsius, relative humidity of 30 to 60 percent, 12 hours of supplemental lighting (06:00 to 18:00), fertilization with 20-20-20 NPK, and biological pest control with *Orius* spp., *Neoseiulus cucumeris*, *Chrysoperla rufilabris*, *Encarsia formosa,* and *Aphidius colemani*.

The mapping family derives from a cross between lines with contrasting bagged-capitulum filled-seed endpoints and herbicide responses. Pollen was transferred by rubbing donor and recipient capitula, and crossed heads were bagged to exclude foreign pollen. HTS_006, which had the filled-seed and herbicide-tolerant endpoints, founded the family analyzed here. Accession provenance and the mapping-family pedigree are given in Supplemental Table S1.

### Filled-seed endpoint phenotyping

The binary score records the bagged-capitulum filled-seed endpoint. It does not resolve self-compatibility, the reproductive mechanism, partial fertility, or temporal variation in fertility.

Capitula were bagged at anthesis without directed pollination. Endpoint = 1 was assigned to the filled-seed endpoint; endpoint = 0 was assigned to the no-filled-seed endpoint under the same assay. Among the final-panel individuals, 166 were scored: 84 had endpoint = 1 (filled-seed endpoint) and 82 had endpoint = 0 (no-filled-seed endpoint); 15 were unscored.

### Herbicide-tolerance phenotyping

Genotyped individuals were classified as herbicide-tolerant or susceptible for binary mapping on the basis of survival following a single pre-emergence application of the protoporphyrinogen oxidase (PPO) inhibitor sulfentrazone (Spartan 4F) at 280 g active ingredient per hectare, delivered with a Generation 4 Research Track Sprayer at the rate used to verify the tolerant parents. Seed was sprayed two days after sowing; individuals that germinated and produced healthy seedlings were classified tolerant, and those that failed to germinate or died before the first true leaf expanded were classified susceptible. The susceptible line OH-1004 “Zaborski” was included in every screen as a control. Because a single labeled rate was applied, the score records survival and does not separate full tolerance from partial injury.

Among the individuals analyzed by association, 177 of the 181 panel members had an herbicide-response phenotype: 145 tolerant and 32 susceptible, with four unscored (main-text Table 2). The imbalance toward tolerance reflects survival selection before genotyping because susceptible plants were less likely to be advanced and sampled, which reduces power for this endpoint.

### DNA extraction and genome sequencing

Genomic DNA was extracted from *Taraxacum kok-saghyz* OH-1004 and sequenced with PacBio high-fidelity (HiFi) chemistry, yielding approximately 43 to 44 Gb of HiFi reads (mean aligned read depth about 36×) from two single-molecule real-time sequencing cells.

### The OSU TarKok 2.0 assembly

The OSU TarKok 2.0 assembly of *Taraxacum kok-saghyz* (GenBank GCA_047496595.1) is a chromosome-scale assembly produced by Dovetail Genomics. PacBio HiFi contigs from the same library were scaffolded with Dovetail Omni-C in vitro proximity-ligation data. It served as the primary scaffolding reference and as a comparator.

### De novo genome assembly, scaffolding, gap filling, and polishing

A primary contig assembly was produced from the HiFi reads with Verkko v2.3.2 in HiFi-only mode (Rautiainen et al., 2023). The Verkko contigs were polished and scaffolded with RagTag v2.1.0 (Alonge et al., 2022) against a polished version of the OSU TarKok 2.0 assembly and polished for two rounds with Racon v1.5.0 (Vaser et al., 2017) using the HiFi reads aligned with minimap2 v2.30 (map-hifi preset) (Li, 2018), producing a scaffold backbone. Contigs left unplaced on this backbone were then re-placed onto the existing chromosomes with RagTag against the polished OSU TarKok 2.0 assembly (asm5 preset) and, as a secondary reference, against the chromosome-scale *T. kok-saghyz* TK1151 assembly (asm10 preset) (Lin et al., 2022), with the OSU TarKok 2.0 placements weighted twice the TK1151 placements. The two placements were reconciled with the RagTag merge module while the eight chromosomes were held intact. Through this step, approximately 13.5 Mb (about 1% of the assembly) of remaining contigs were successfully assigned to chromosomes, leaving the TK1151 reference used exclusively for this reconciliation without altering the core backbone. Remaining gaps were patched with sequence from an independent hifiasm v0.25.0 assembly (Cheng et al., 2021) and from the OSU TarKok 2.0 reference. The assembly received a final two rounds of Racon polishing with the HiFi reads, which removed forty-eight short zero-coverage contigs. No new long-range data were generated for the present assembly, so its chromosome-scale organization derives from the Omni-C proximity-ligation scaffolding of the same HiFi library in OSU TarKok 2.0, carried across by reference-guided scaffolding rather than re-measured; the additional structural support is described in the Results. The resulting assembly is referred to as OSU TarKok 3.0; its nuclear chromosome-scale scaffolds are named Chr1 to Chr8 in order of decreasing length.

### Assembly evaluation and organellar screening

Assembly statistics were calculated separately for the 857-sequence primary assembly (before organellar filtering), the 658-sequence organellar-filtered nuclear assembly, and the 198-sequence separated organellar set. The nuclear and organellar subsets exactly reconstruct the sequence count, length, N-runs, and N bases of the primary assembly before organellar filtering. Continuity curves were calculated from the 658-sequence organellar-filtered nuclear assembly and its 746 maximal non-N segments as shown in main-text Figure 1. Genome-mode BUSCO v6.0.0 was rerun on the final 658-sequence nuclear assembly using eudicotyledons_odb12 (n = 2,805). It recovered C:98.4% (2,759) [S:91.6% (2,569), D:6.8% (190)], F:1.1% (30), and M:0.6% (16).

Protein-mode BUSCO v6.0.0 was run on the final OSU TarKok 3.0 nuclear proteome of 34,637 representative proteins, one per final nuclear gene model, against eudicotyledons_odb12 (n = 2,805; Manni et al., 2021). It recovered C:97.5% (2,734) [S:92.1% (2,583), D:5.4% (151)], F:1.3% (37), and M:1.2% (34). Jellyfish and GenomeScope2 were used for k-mer profiling (Marçais & Kingsford, 2011; Ranallo-Benavidez et al., 2020).

### Reference-guided chloroplast consensus and annotation

The 151,316-bp product is a reference-guided chloroplast consensus. PacBio HiFi reads from the two accession-OH-1004 libraries were aligned to OR990541.1 with minimap2 map-hifi (Li, 2018); primary alignments were retained with SAMtools (Li et al., 2009), and read names were deduplicated with SeqKit rmdup (Shen et al., 2016). This process retained 72,408 recruited reads. The recruited reads were remapped to OR990541.1 and randomly subsampled at fraction 0.043 to approximately 300-fold depth. BCFtools mpileup used -d 600, -q 0, and -Q 13; haploid variants were called with call --ploidy 1 -m -v, filtered at QUAL ≥ 30 and DP ≥ 50, and applied with BCFtools consensus (Danecek et al., 2021). The 213 retained substitutions and zero indels produced a 151,316-bp consensus with coverage at all positions.

A BLAST+ self-alignment (Camacho et al., 2009) defined the longest reverse-complement match as the inverted repeat. The 83,900-bp LSC, 18,538-bp SSC, and two 24,439-bp inverted repeats sum to 151,316 bp. The artificial end-to-start junction was spanned by 6,143 HiFi alignments, and complete tiling was consistent with circular closure of the reference-guided consensus. PGA, run in circular mode with the OR990541.1 GenBank record as the principal annotation reference, provided the feature calls (Qu et al., 2019). An independent BLAST+ inventory recovered 80 CDS, 27 tRNAs, and four rRNAs (Camacho et al., 2009).

### Repeat, gene, and non-coding-RNA annotation

Repetitive sequence was modeled de novo with RepeatModeler2 v2.0.8 (Flynn et al., 2020), the family library was reclassified with TEsorter against the REXdb viridiplantae database (Zhang et al., 2022; Neumann et al., 2019) and RepeatClassifier, and the genome was masked with RepeatMasker v4.2.3 (Smit et al., 2013–2015). A structural transposable-element annotation was produced independently with EDTA v2.2 (Ou et al., 2019), and the assembly-wide LTR Assembly Index was computed with LTR_retriever (Ou & Jiang, 2018). Repeat occupancy (77.5%), transposable-element occupancy (59.8%), and LAI (41.6) were calculated on the 857-sequence primary assembly before organellar partitioning. Protein-coding genes were predicted with BRAKER3 v3.0.8 (Gabriel et al., 2024), integrating RNA-seq evidence aligned with HISAT2 v2.2.2 (Kim et al., 2019) and a 264-proteome UniProt eudicot reference set (The UniProt Consortium, 2023). The final nuclear annotation contained 34,637 protein-coding genes and 39,326 transcripts. A representative proteome containing one protein per gene was generated by selecting the longest translation-valid model with complete coding boundaries when available, using transcript ID for deterministic tie breaking. Nine documented partial-gene exceptions retained the longest translation-valid partial model. Expression was not used for representative-protein selection. Proteins were functionally annotated with InterProScan v5.78-109.0 and eggNOG-mapper v2.1.12, and non-coding RNA genes with tRNAscan-SE v2.0.12, barrnap v0.9, and Infernal v1.1.5 (Jones et al., 2014; Cantalapiedra et al., 2021; Chan & Lowe, 2019; Nawrocki & Eddy, 2013). Non-coding RNA summaries were restricted to predictions retained on the final 658-sequence nuclear assembly. Abundant rRNA-model matches were retained as uncurated prediction rows and were not interpreted as a biological rRNA gene count.

### Comparative synteny

Whole-genome macrosynteny was computed with ntSynt v1.0.8 using minimizers of length 24 and window 1000, OSU TarKok 3.0 as target, and orientation normalization, followed by ntSynt-viz (Coombe et al., 2025). Comparisons used TK1151, *T. mongolicum* TM5, and *Lactuca sativa* RefSeq GCF_002870075.5 (Lin et al., 2022; Reyes-Chin-Wo et al., 2017). Panels A and B of main-text Figure 2 display all retained blocks shared among OSU TarKok 3.0, TK1151, and *T. mongolicum*; no 100-kb plotting threshold was applied. Panel C displays the retained four-genome blocks that also include lettuce. The lettuce comparison anchors 222,337,000 bp, or 19.03%, of the eight OSU chromosome scaffolds, and correspondence was summarized as shares of each OSU chromosome’s anchored syntenic length. Orientation was normalized for visualization and was not interpreted as inversion evidence. Multichromosomal correspondence is compatible with rearrangement or fusion but does not establish direction, event order, or mechanism.

### Single-copy phylogeny and genome divergence

The nine-genome phylogeny used 1,480 complete single-copy eudicotyledons_odb12 BUSCO proteins present in all genomes and the 857-sequence OSU primary assembly before organellar filtering. Per-ortholog MAFFT alignments were trimmed with trimAl and concatenated to 769,094 amino-acid columns (Katoh & Standley, 2013; Capella-Gutiérrez et al., 2009). IQ-TREE v3.1.2 used ModelFinder, which selected JTT+F+I+R7 by BIC, and 1,000 ultrafast bootstrap replicates (Wong et al., 2026; Kalyaanamoorthy et al., 2017; Hoang et al., 2018).

Comparative nucleotide identity was calculated from the best minimap2 alignments of 5-kb fragments from each comparator query assembly to the eight OSU chromosome-scale scaffolds, retaining aligned spans of at least 2 kb (Li, 2018). Identity is length-weighted and conditional on retained alignments; aligned fraction is retained aligned query length divided by the corresponding comparator assembly length (Supplemental Table S4).

### Gene-family copy number across the tribe

Rubber-pathway families were first reconciled against characterized proteins and mapped uniformly to all nine assemblies with Miniprot (Li, 2023), retaining protein identity at least 0.60 and coverage at least 0.50. CPT, CPTL/RTA, CitBP, SRPP, REF, and HMGR were kept as distinct families in the exported count table. Exact projected counts for the six families across all nine assemblies are provided in Supplemental Table S8, Panel A.

The structural prediction *g12582* contains *TkCPT1*-and *TkCPT2*-like isoforms and is retained as an unresolved locus for expression. Full-length protein identity supports *g5630* as *TkCPTL1*/*TbRTA*; *g9503* remains a probable divergent CPTL paralog. No paralog-specific TPM was derived from the combined *g12582* prediction.

### Orthology inference and gene-family evolution

OrthoFinder v3.1.5 (Emms et al., 2026; Emms & Kelly, 2019) analyzed eight one-protein-per-gene proteomes. All eight inputs were verified as one-protein-per-gene FASTAs. The OSU input contained 34,637 proteins, one per nuclear GFF gene. Comparator inputs were generated by selecting the longest protein associated with each annotated gene or locus identifier. The eight inputs contained 309,333 proteins representing 309,333 distinct genes. Actual input counts and the corresponding isoform-level source counts are reported separately in Supplemental Table S2. Similarity search, clustering, multiple alignment, and gene trees used DIAMOND, MCL, FAMSA, and FastTree, respectively (Buchfink et al., 2021; Enright et al., 2002; Deorowicz et al., 2016; Price et al., 2010). The rooted species tree was made ultrametric with root age 100. Twenty-seven families with more than 100 copies in any taxon were excluded. CAFE5 v5.1.0 received 33,822 families and filtered 9,797 inferred absent at the root, leaving 24,025 tests. The reported model used a single global birth-death rate, family P-value calculation, 32 threads, and no error model (Mendes et al., 2020); family P values were adjusted across all 24,025 tests by the Benjamini–Hochberg method (Benjamini & Hochberg, 1995). Branch probabilities were unadjusted descriptive outputs conditional on the fitted family model. An alternative error-model fit was discarded because it estimated a negative error probability and produced invalid probability values; none of its outputs were used. These orthogroup assignments and family counts are release-bound to OrthoFinder v3.1.5 and the eight stated one-protein-per-gene proteomes.

### RNA extraction and sequencing

RNA was extracted from leaf, flower, crown, root, and a pooled floral and seed sample of *Taraxacum kok-saghyz*. Tissue was collected from greenhouse-grown plants, flash-frozen in liquid nitrogen and stored at -80 degrees Celsius until extraction. RNA was extracted from all tissues using the high-salt CTAB method described in Wong et al. (2014). DNAse treatment was then performed using the DNA-free™ DNA Removal Kit (ThermoFisher Scientific) according to manufacturer’s instructions. RNA concentration was measured using a Qubit™ 4 and Qubit™ RNA High Sensitivity (HS) Assay Kit (ThermoFisher Scientific), then RNA integrity threshold (RIN) was measured using a TapeStation (Agilent, Santa Clara, CA, USA). Samples with a RIN ≥7 were then used to make RNA libraries using the NEBNext® Ultra™ II RNA Library Prep Kit for Illumina® (New England BioLabs® Inc.) with two biological replicates per tissue. The concentration and quality of the twelve resulting libraries were assessed by fluorometry and electrophoresis as above, then they were sequenced on an Illumina NovaSeq 6000 platform as 2 x 151 bp paired-end reads.

### Rubber-pathway expression, co-expression networks, and biosynthetic gene clusters

The in-house experiment comprised 12 libraries. The contrast compared underground tissues (crown and root; n = 4 libraries) with aerial tissues (leaf, flower, and pooled floral/seed; n = 8 libraries). Reads were aligned with HISAT2, featureCounts provided raw counts, TPM was used for display, and DESeq2 tested differential expression (Kim et al., 2019; Liao et al., 2014; Love et al., 2014). The prespecified criterion for underground-tissue enrichment was an underground:aerial ratio >1 and DESeq2 adjusted P <0.05.

The primary co-expression analysis used the combined 36-library signed WGCNA network (27,721 expressed genes; 37 biological modules after excluding grey). Module eigengenes were correlated with seven mutually exclusive categories: crown (n = 2), flower (n = 2), latex (n = 6), leaf (n = 8), petiole (n = 3), root (n = 11), and seed/floral pool (n = 4). Benjamini-Hochberg adjustment was applied once across all 259 module–tissue P values. The 12-library network was inferred separately as a sensitivity analysis (43 biological modules after excluding grey; leaf, flower, crown, and root n = 2 each and seed/floral pool n = 4; 215 tests). Main-text Figure 5B used the 36-library signed TOM at power 16 for a fixed 16-node set. Each node contributed its two strongest finite TOM links; unordered duplicates were collapsed, gene ID provided a deterministic tiebreaker, and all rubber-core pairs with TOM ≥0.27 were added, yielding 34 undirected edges. The *g12582*–*g261* pair is not an edge.

plantiSMASH v2.0.4 under Python 3.8.19 (Kautsar et al., 2017) analyzed the 857-sequence primary assembly before organellar filtering together with its corresponding 34,737-gene prepartition BRAKER gene/mRNA/CDS annotation. The analysis used the plants taxon, nucleotide input, eukaryotic mode, supplied gene models without additional gene finding, CD-HIT clustering, and no result limit; ClusterBLAST, KnownClusterBLAST, and fullhmmer were not enabled. The analysis identified 42 candidate clusters comprising 706 gene–cluster assignments involving 686 unique genes; 20 genes were assigned to two overlapping clusters. All 686 genes are retained in the final nuclear annotation.Predicted cluster classes and their overlap with rubber-related loci are summarized in Supplemental Figure S7 and Supplemental Table S7.

For Supplemental Figure S10, strand-aware 1-kb promoters were clipped at the nearest upstream gene boundary. Seven IUPAC elements were tested by two-sided Fisher exact tests with Benjamini-Hochberg adjustment. AME v5.5.9 in the MEME Suite (McLeay & Bailey, 2010; Bailey et al., 2015) tested 656 JASPAR CORE 2022 plant matrices (Castro-Mondragon et al., 2022) at E <1.

### seqWell library processing and variant calling

Genomic DNA libraries were prepared using seqWell multiplexed library-preparation technology. Paired-end reads from these libraries were re-synchronized with BBMap repair (Bushnell, 2014) and processed with nf-core/sarek v3.5.1 on Nextflow v24.10.4 (Di Tommaso et al., 2017; Ewels et al., 2020; Garcia et al., 2020). The two-caller consensus used GATK HaplotypeCaller v4.6.2.0 (McKenna et al., 2010) and BCFtools v1.21 (Danecek et al., 2021). GenMap v1.3.0 (Pockrandt et al., 2020) was indexed against the 857-sequence primary assembly and run with k = 150 and E = 2 allowed errors. The eight analysis chromosomes are sequence-identical and equal in length to Chr1-Chr8 of the final 658-sequence nuclear reference, so chromosome coordinates were unchanged by organellar partitioning and removal of the unplaced contaminant. BedGraph values ≥0.999999 were treated as uniquely mappable and merged within Chr1-Chr8. RepeatMasker intervals were independently restricted and merged on the same chromosomes; their complement defined nonrepeat sequence, and its intersection with the uniquely mappable set defined the conservative mappable genome used for variant filtering (Quinlan & Hall, 2010). For gene accessibility, CDS intervals were merged across transcripts within each chromosome-anchored gene before intersection with the conservative mappable genome; chromosome-wide CDS intervals were additionally merged across genes to prevent double counting (Supplemental Table S10).

### Variant filtering, association, and family-based QTL mapping

Four passing-site lists were intersected. The HDplot implementation used H <0.60 and D <7.0 for the singleton class (McKinney et al., 2017). VCFtools --hardy (Danecek et al., 2011) retained sites with P_HET_EXCESS ≥1× 10⁻⁶ or a missing excess P value. Mean depth was calculated over called panel genotypes; the upper cutoff was median + 3 × 1.4826 × MAD, with no lower-depth cutoff. For the selfed HTS_006 family, a homozygous-founder site failed when the progeny Mendelian inconsistency rate exceeded 0.05. For a heterozygous or missing founder, a site failed when progeny heterozygosity exceeded 0.85.

PLINK2 v2.0.0-a.6.9LM used --maf 0.05, --geno 0.5, --mind 0.9, --snps-only, and --max-alleles 2 (Chang et al., 2015). LD pruning used --indep-pairwise 50 5 0.2; ten PCs were calculated, and PC1-PC5 entered the association models. Variant identifiers were set from chromosome, position, reference, and alternate alleles.

Filled-seed endpoint = 1 and no-filled-seed endpoint = 0; missing scores remained NA. Association models were GEMMA v0.98.5 linear mixed models (Zhou & Stephens, 2012), GCTA v1.94.1 MLMA-LOCO (Yang et al., 2011), and PLINK2 v2.0.0-a.6.9LM logistic regression with Firth fallback using --glm firth-fallback (Firth, 1993; Chang et al., 2015). PC1-PC5 were included in all models. GEMMA and GCTA used a Bonferroni threshold of P = 0.05/1,099,707. PLINK2 used phenotype-specific 5% genome-wide thresholds obtained from 1,000 phenotype permutations: P = 2.61345 × 10⁻⁶ for filled seed and P = 5.62516 × 10⁻⁶ for herbicide response. Cross-model support required the same chromosome:position:reference:alternate allele record to pass its model-specific threshold in at least two models. Genomic-inflation factors were calculated from all valid P values in each primary model output. Model-specific results are summarized in Supplemental Table S11 and Supplemental Figure S14.

The family scan used R 4.3.3 (R Core Team, 2024) for additive logistic-regression likelihood-ratio tests at 10,737 biallelic allele records from 10,617 unique founder-heterozygous positions; decomposition of multiallelic sites accounts for the 120 additional allele records. Missing genotypes were omitted per marker; markers required at least 20 complete F2 and nonzero genotype variance. LRT = 2(logLik(full)−logLik(null)) and LOD = (logLik(full)−logLik(null))/ln(10). Recorded marker P values were converted with the chi-square(df = 1) inverse survival function. The alpha = 0.05 threshold was converted through the same monotone function from the 95th percentile of the maximum −log10(P) in 1,000 phenotype permutations. The connected peak component at LODpeak−1.5 was located in the complete physical-order Chr1 LOD vector, and each boundary was linearly interpolated between the inner marker and its below-cutoff flanking marker. Lep-MAP3 ParentCall2 used removeNonInformative = 1; Filtering2 used dataTolerance = 0.0001; SeparateChromosomes2 evaluated lodLimit = 8, 10, 12, and 15, selecting the map closest to eight groups with at least 50 markers. The selected map contained nine linkage groups for eight chromosomes because Chr1 remained split into LG8 and LG9. OrderMarkers2 used useKosambi = 1, sexAveraged = 1, numThreads = 4, and randSeed = 42 (Rastas, 2017; Kosambi, 1944). ParentCall2 retained 7,624 markers, OrderMarkers2 ordered 4,474, and the chromosome-dominant Marey subset retained 4,466 after removing eight off-dominant markers.

For Supplemental Figure S13, OrderMarkers2 positions were oriented to increasing physical position and fitted within each linkage group by equal-weight pool-adjacent-violators isotonic regression; fitted genetic position between markers was obtained by linear interpolation. Physical windows were 2,000,000 bp with a 2,000,000-bp step (zero overlap), with the terminal window truncated at the chromosome end. For each window, the linkage group with greatest marker-span overlap was used, and rate was the fitted cM difference divided by the marker-covered physical span in Mb. Of 589 genome windows, 61 without marker-span overlap were NA and 528 had a rate; fitted zero rates were retained as zero. Repeat fraction was the RepeatMasker-covered fraction from bedtools coverage (Quinlan & Hall, 2010). Thus n = 528 is the number of non-NA rate windows used for the descriptive Spearman correlation, not the total number of physical windows.

### Supplemental Figures

**Supplemental Figure S1.**
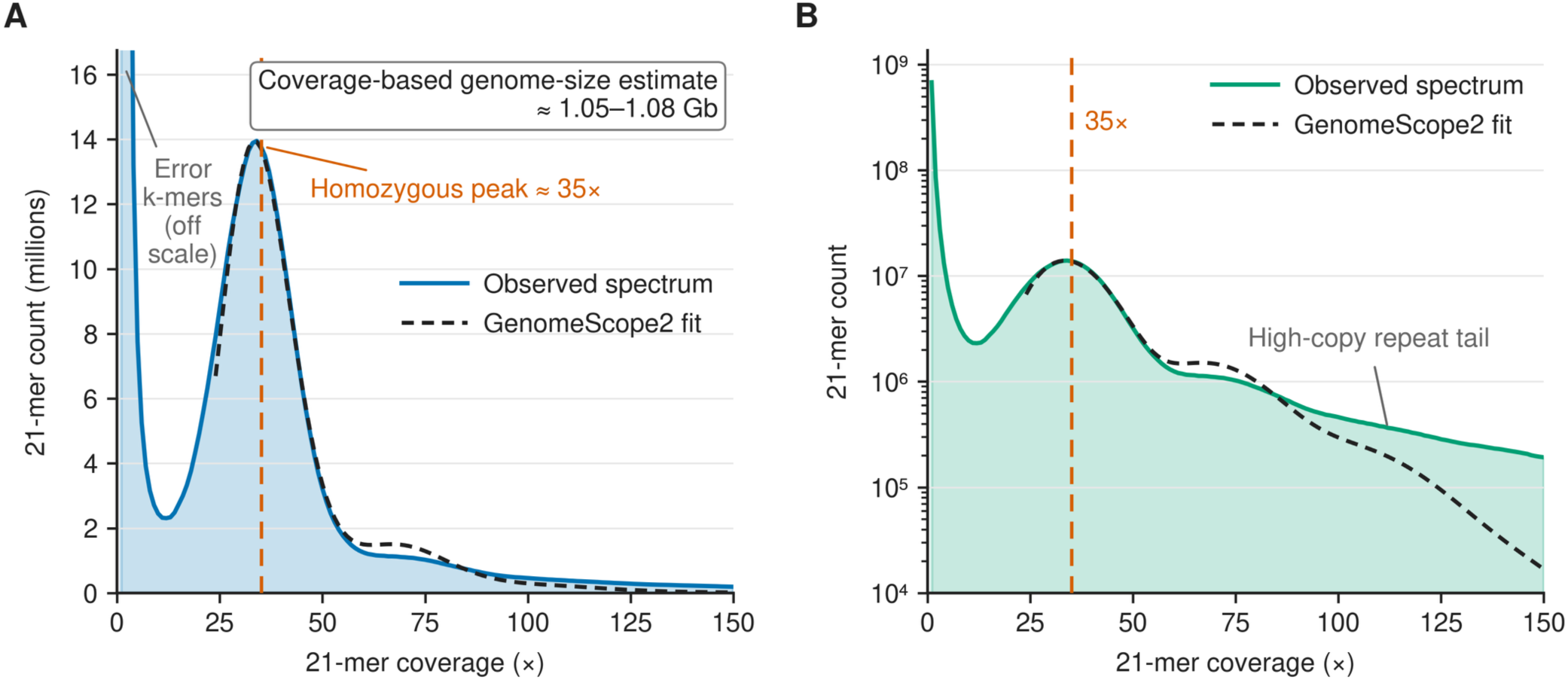
HiFi 21-mer spectrum and coverage-based genome-size estimate. (A) Linear-scale 21-mer spectrum with the GenomeScope2 fit and a homozygous coverage peak near 35×. The direct GenomeScope2 fit did not capture the extensive high-copy repeat tail, so genome size was estimated independently from total k-mer abundance divided by peak coverage at approximately 1.08 Gb. (B) The same spectrum on a logarithmic y-axis, showing the high-copy repeat tail that extends beyond the principal fitted region. The approximately 1.08-Gb value is a coverage-based estimate and not the direct GenomeScope2 fitted interval.

**Supplemental Figure S2.**
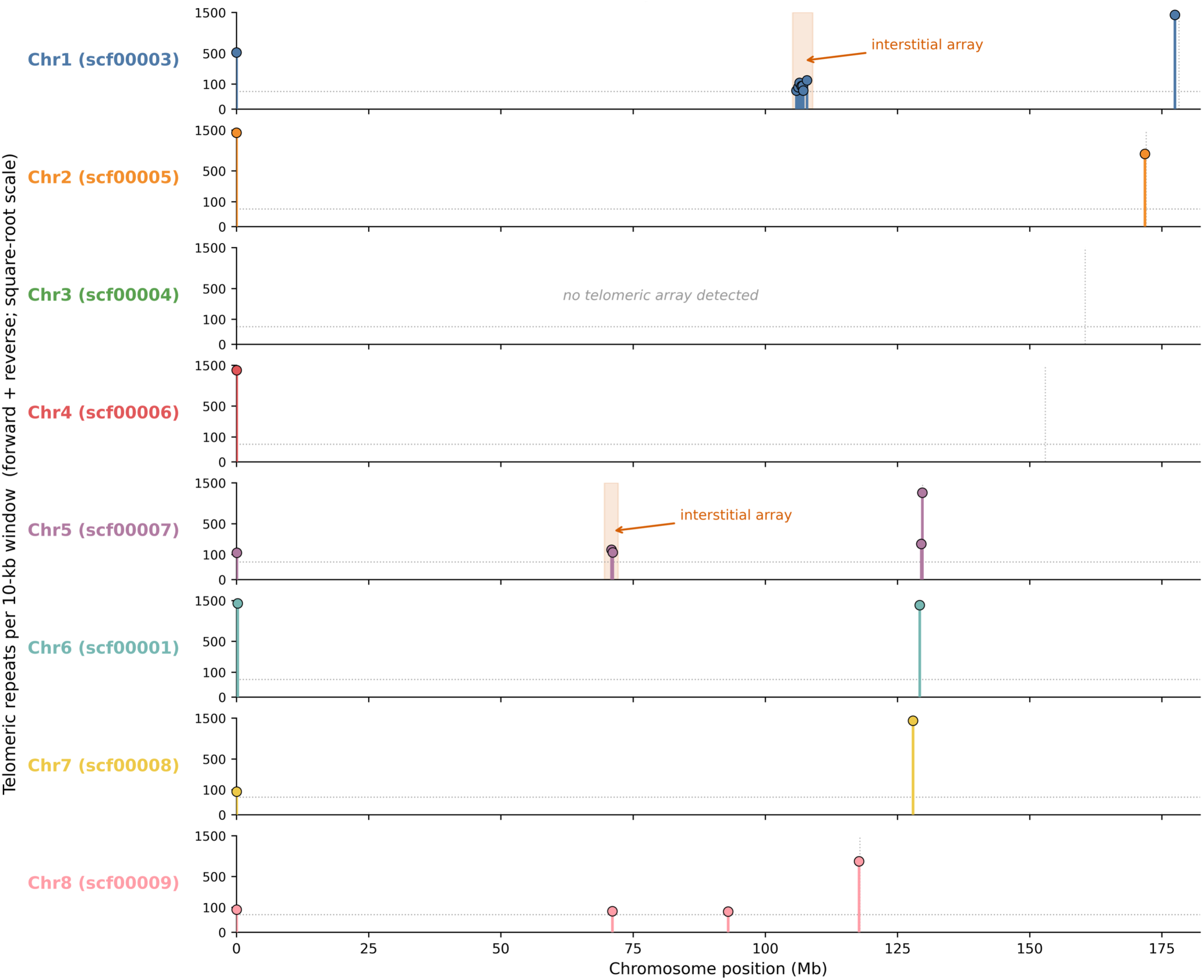
Genome-wide telomeric-repeat (AAACCCT) density, showing terminal arrays at 12 of 16 chromosome ends and interstitial arrays on two chromosomes. The interstitial arrays are compatible with candidate fusion regions but do not establish a specific chromosome-fusion history. Colored markers = telomeric arrays (>50 repeats/1 kb): at chromosome ends = terminal telomeres; red shaded bands = interstitial arrays on Chr1 and Chr5 (candidate chromosome-fusion sites)

**Supplemental Figure S3.**
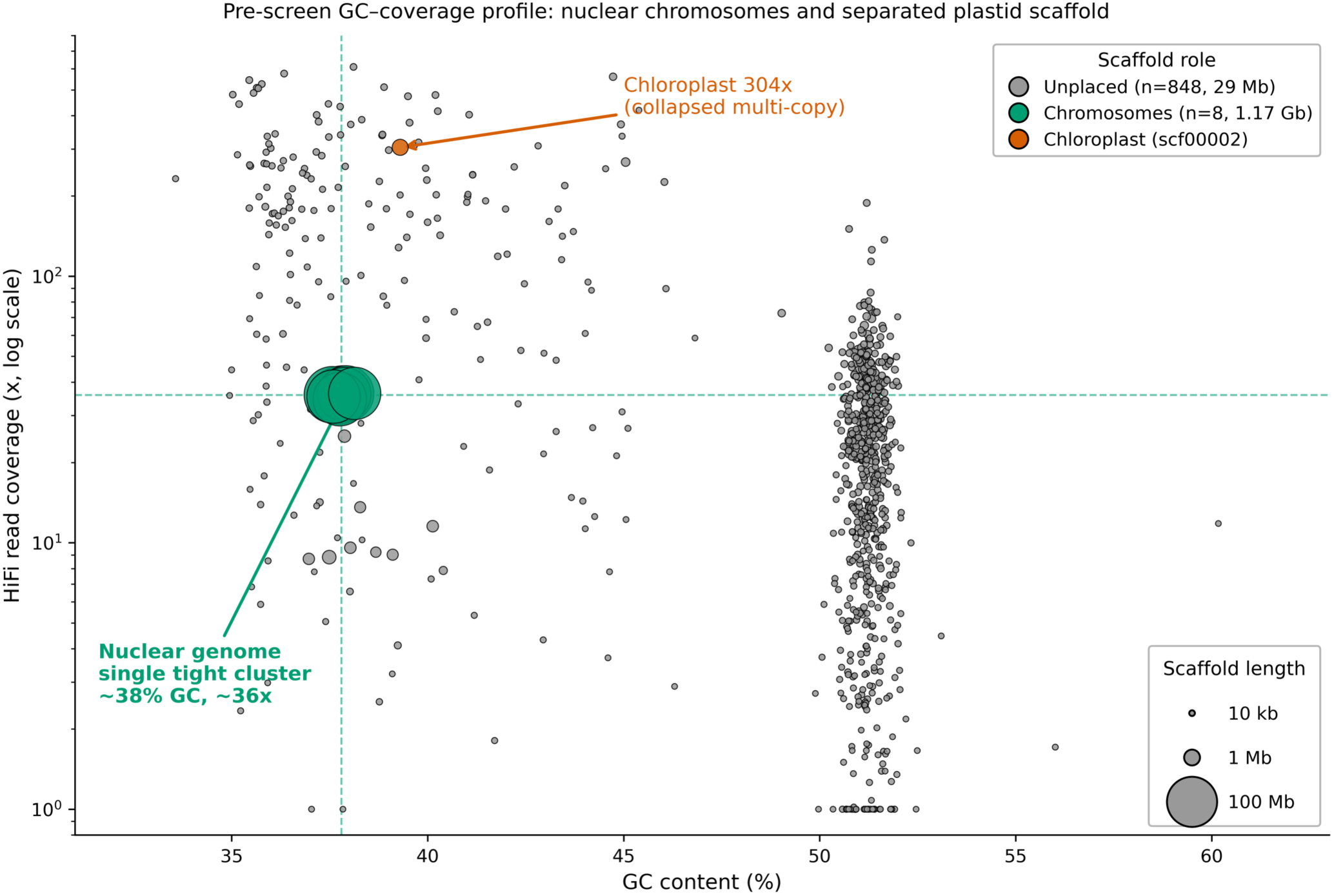
GC-versus-coverage blob plot for the 857-sequence primary assembly before organellar filtering. The separated high-copy, high-GC group contains sequences assigned to the 198-sequence organellar set; the 1,051,501-bp plastid scaffold shown here is absent from the 659-sequence organellar-filtered nuclear assembly. Aside from the separated organellar group, no additional high-abundance cluster was evident. Screening against the NCBI UniVec database returned zero hits at ≥95% identity over ≥30 bp; these screens are not a taxonomically exhaustive assay.

**Supplemental Figure S4.**
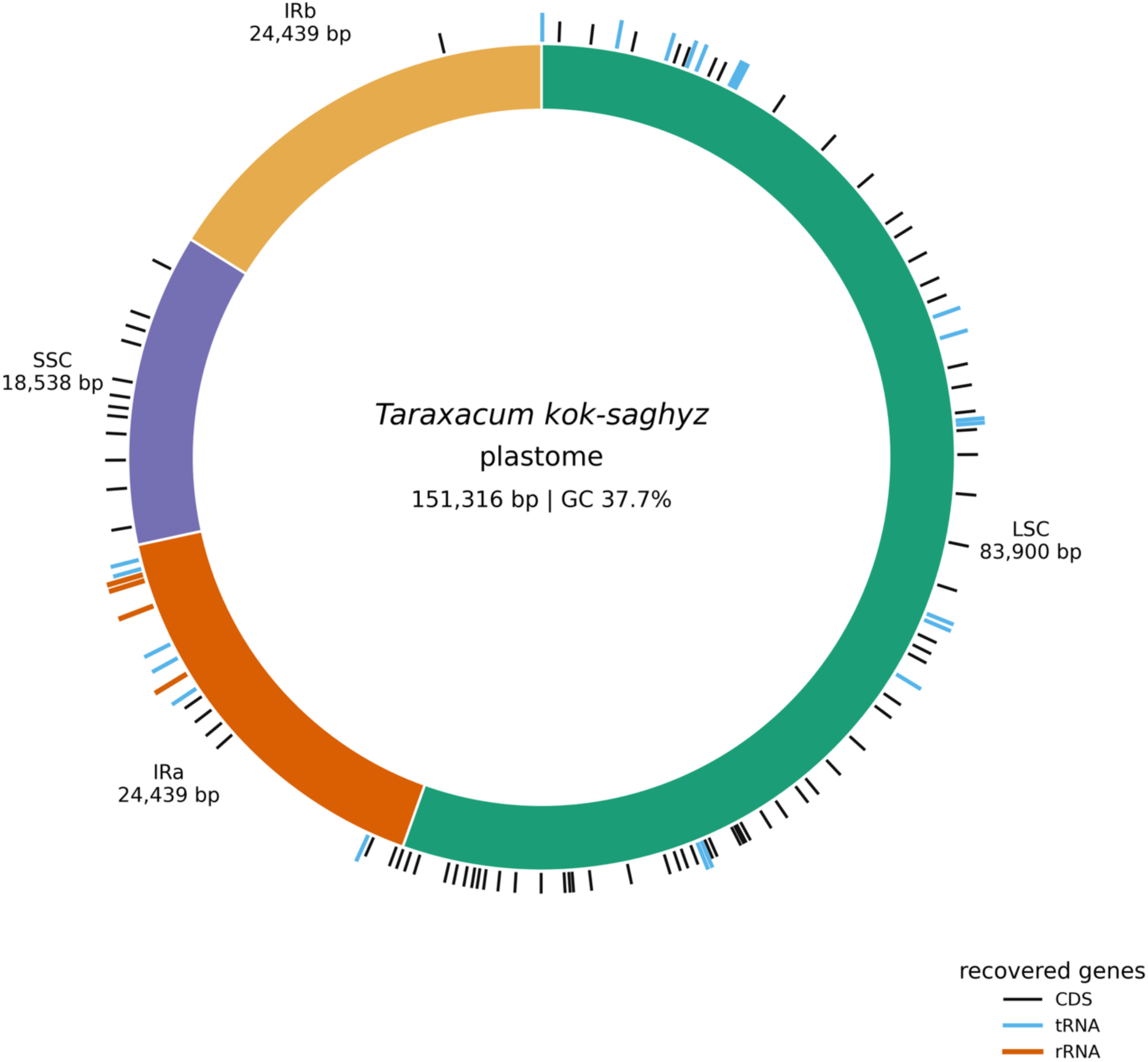
Structure of the 151,316-bp reference-guided chloroplast consensus. The large single-copy region is 83,900 bp, the small single-copy region is 18,538 bp, and each inverted repeat is 24,439 bp. Consensus generation used HiFi reads recruited and aligned to OR990541.1; PGA provided the feature annotation.

**Supplemental Figure S5.**
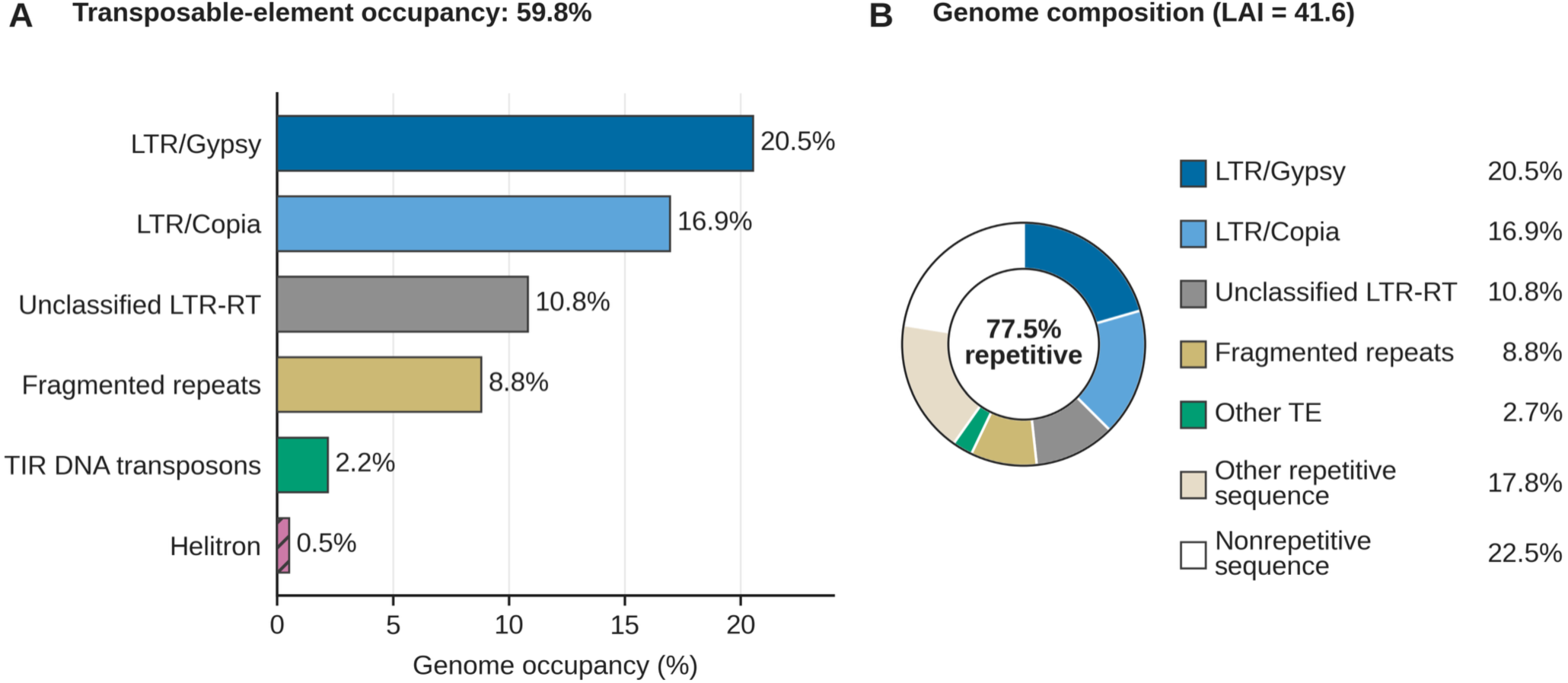
Transposable-element composition calculated on the 857-sequence primary assembly before organellar partitioning. Genome occupancy by transposable-element superfamily is shown with the total repetitive fraction (77.5%), transposable-element fraction (59.8%), and LTR Assembly Index (41.6). Repeat and transposable-element occupancy were calculated on the primary assembly before organellar partitioning. The length-weighted LAI of 41.6 was calculated across the eight chromosome-scale scaffolds.

**Supplemental Figure S6.**
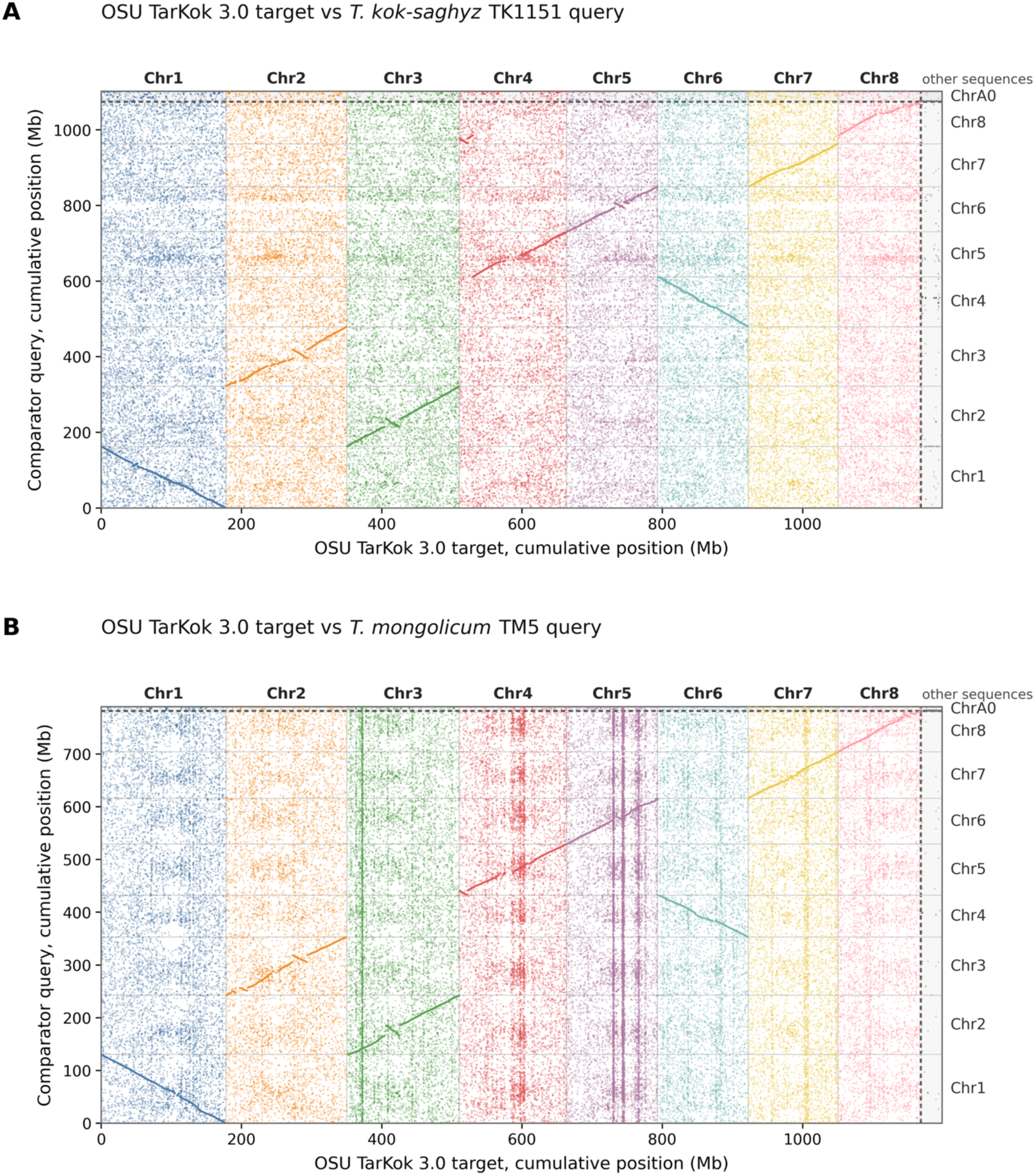
Whole-assembly dot plots using the OSU nuclear assembly as query and TK1151 or *T. mongolicum* as comparator queries. The plots provide the alignment source for the retained 5-kb-fragment identity summaries; aligned-fraction denominators are the corresponding comparator query assemblies (Supplemental Table S4).

**Supplemental Figure S7.**
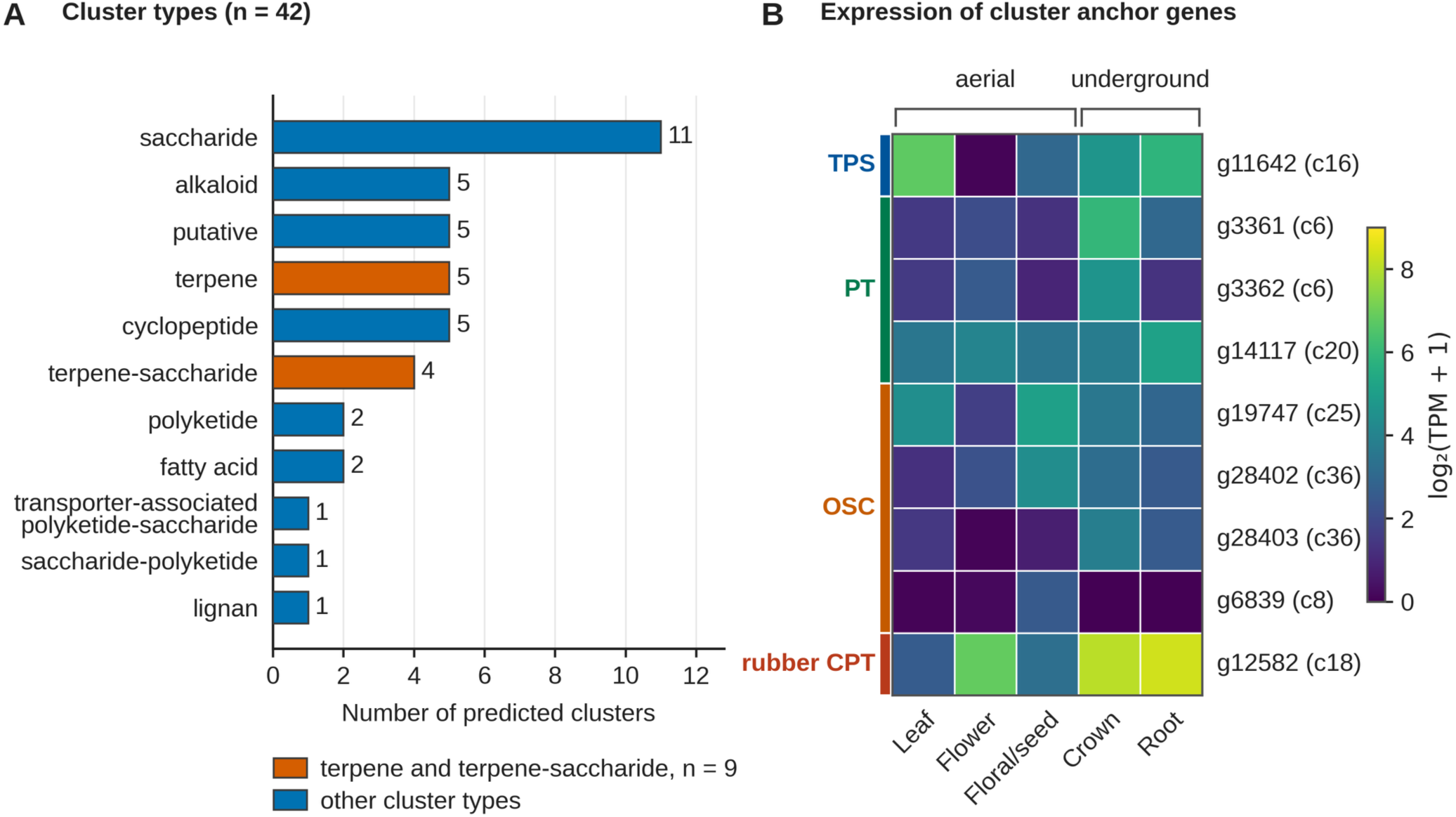
plantiSMASH biosynthetic-gene-cluster predictions and their relationship to rubber and terpenoid genes. Rubber-pathway loci are predominantly dispersed. Only the *g12582* locus overlapped a predicted plantiSMASH cluster, classified as a lignan cluster; no other curated rubber-related locus occurred within a predicted cluster.

**Supplemental Figure S8.**
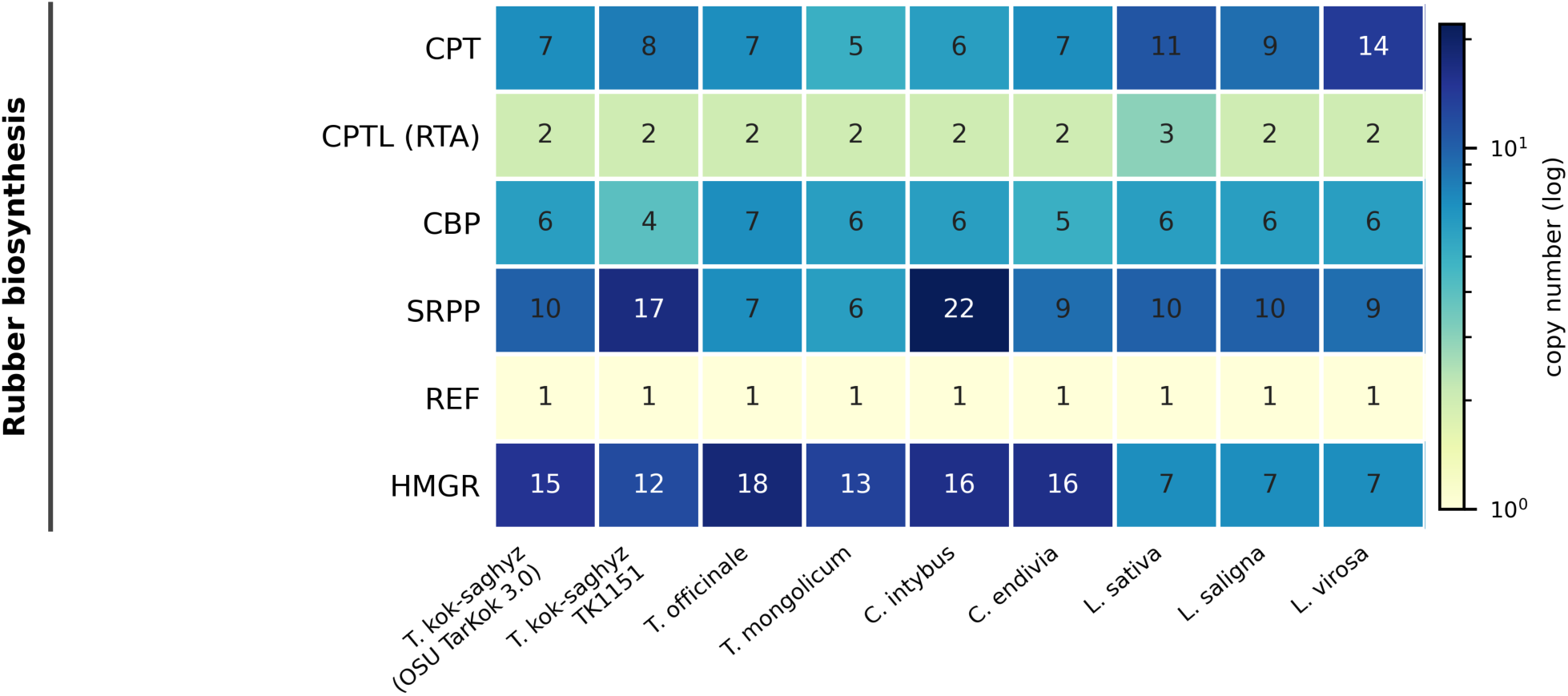
Total projected CPT, CPTL/RTA, citrate-binding protein (CitBP), SRPP, REF, and HMGR homology-cluster counts from uniform Miniprot mapping across nine Cichorieae assemblies. These broad projections are distinct from curated expression loci and from the individual global orthogroups tested by CAFE5. Total projected HMGR-family size is not uniquely elevated in *T. kok-saghyz*; HMGR-containing OG0000471 is evaluated separately in Supplemental Figure S9. Exact projected counts for all six families and nine assemblies are provided in Supplemental Table S8, Panel A.

**Supplemental Figure S9.**
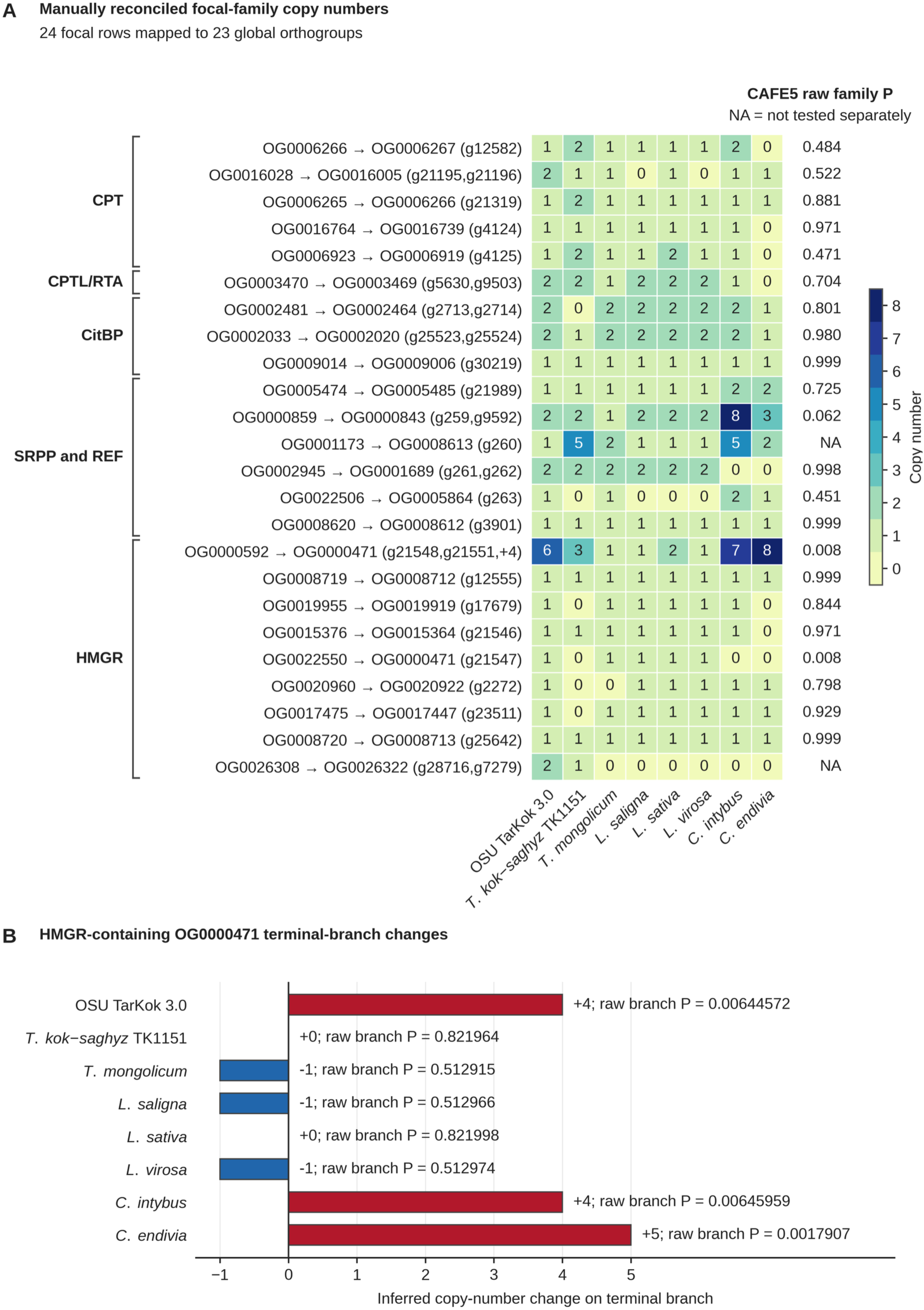
OrthoFinder/CAFE5 results using the final OSU representative protein set of 34,637 proteins. (A) Manually reconciled copy-number vectors for 24 focal rows mapped to 23 global orthogroups. Displayed family P values are raw and carry no significance symbols. (B) Inferred terminal-branch changes for HMGR-containing OG0000471. Bar color (Blue or red) indicates direction only, not significance. The family result is raw P = 0.008; family BH q = 0.160, and the inferred OSU change is +4 with raw branch probability = 0.00644572. Branch probabilities are unadjusted descriptive outputs conditional on the fitted model; BH was applied only to family P values. These statistics do not describe the total projected HMGR-family count.

**Supplemental Figure S10.**
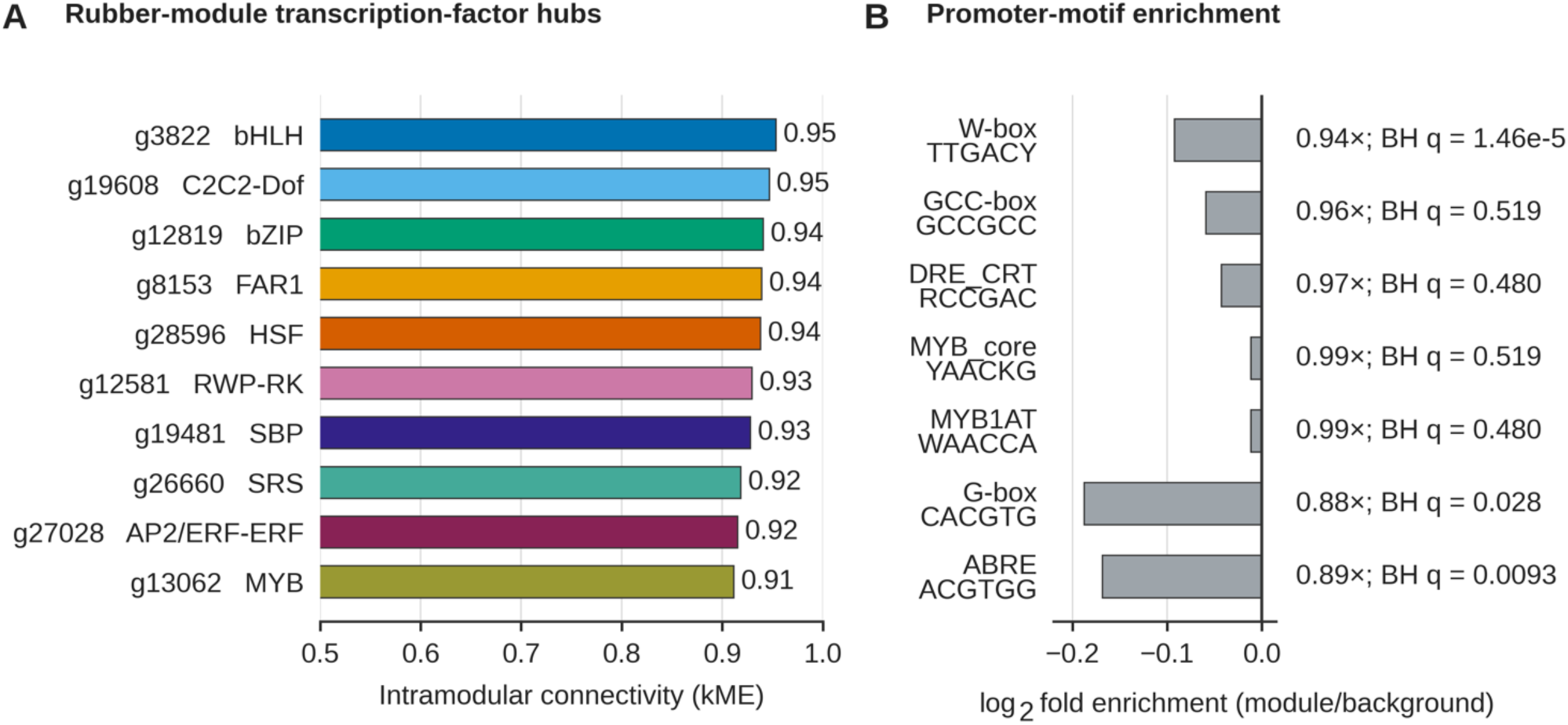
Transcription-factor hubs and promoter-motif tests for the combined 36-library rubber module. (A) The ten sequence-specific transcription factors with the highest intramodular connectivity (kME). (B) Strand-aware, neighbor-clipped 1-kb promoters were scanned for seven prespecified IUPAC motifs; two-sided Fisher exact tests were adjusted across motifs by Benjamini-Hochberg. No significant enrichment was detected and AME against JASPAR Plants returned zero motifs at E <1. Significant adjusted P values corresponded to motif depletion rather than enrichment.

**Supplemental Figure S11.**
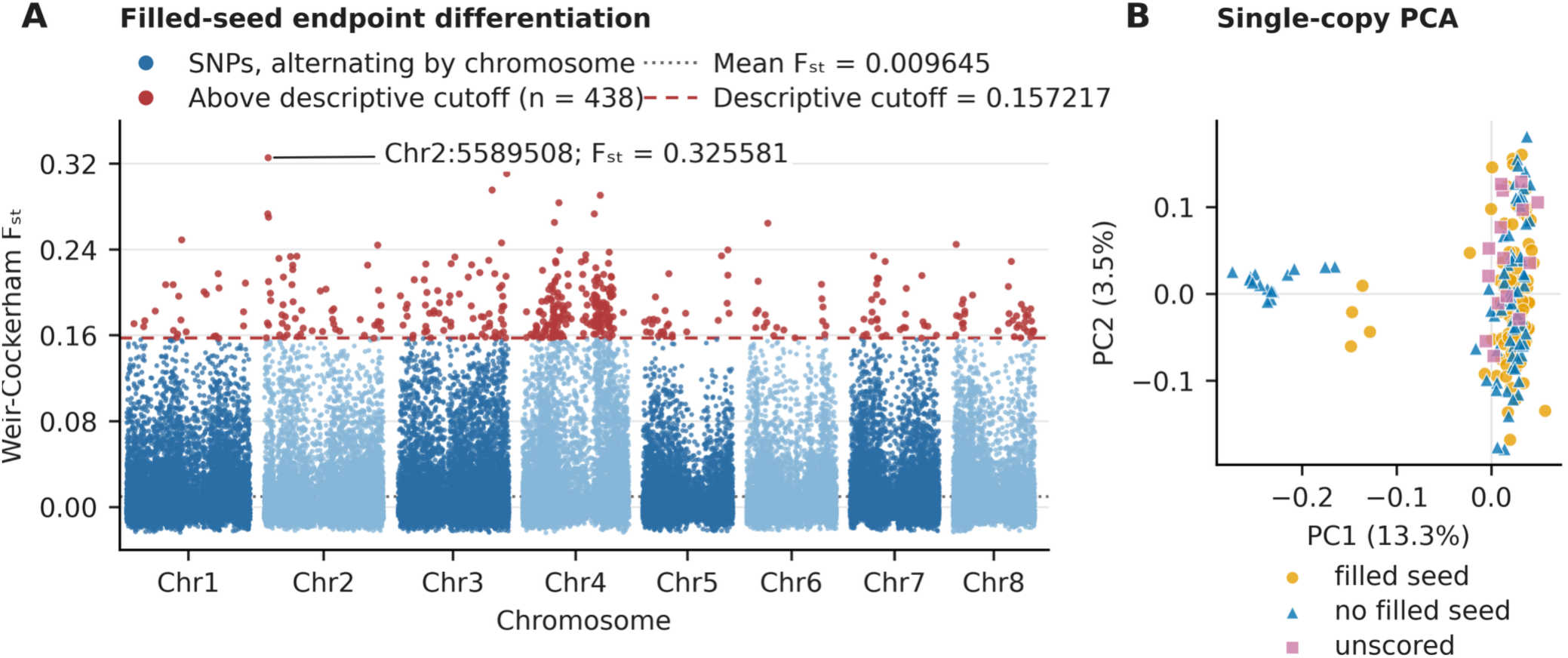
Callset-specific filled-seed versus no-filled-seed endpoint differentiation. (A) PLINK2 per-SNP Weir–Cockerham FST from 92,651 LD-pruned single-copy markers in 166 scored individuals (84 endpoint = 1; 82 endpoint = 0; mean 0.009645; maximum 0.325581). (B) Principal components for all 181 single-copy panel samples, including the 15 unscored individuals.

**Supplemental Figure S12.**
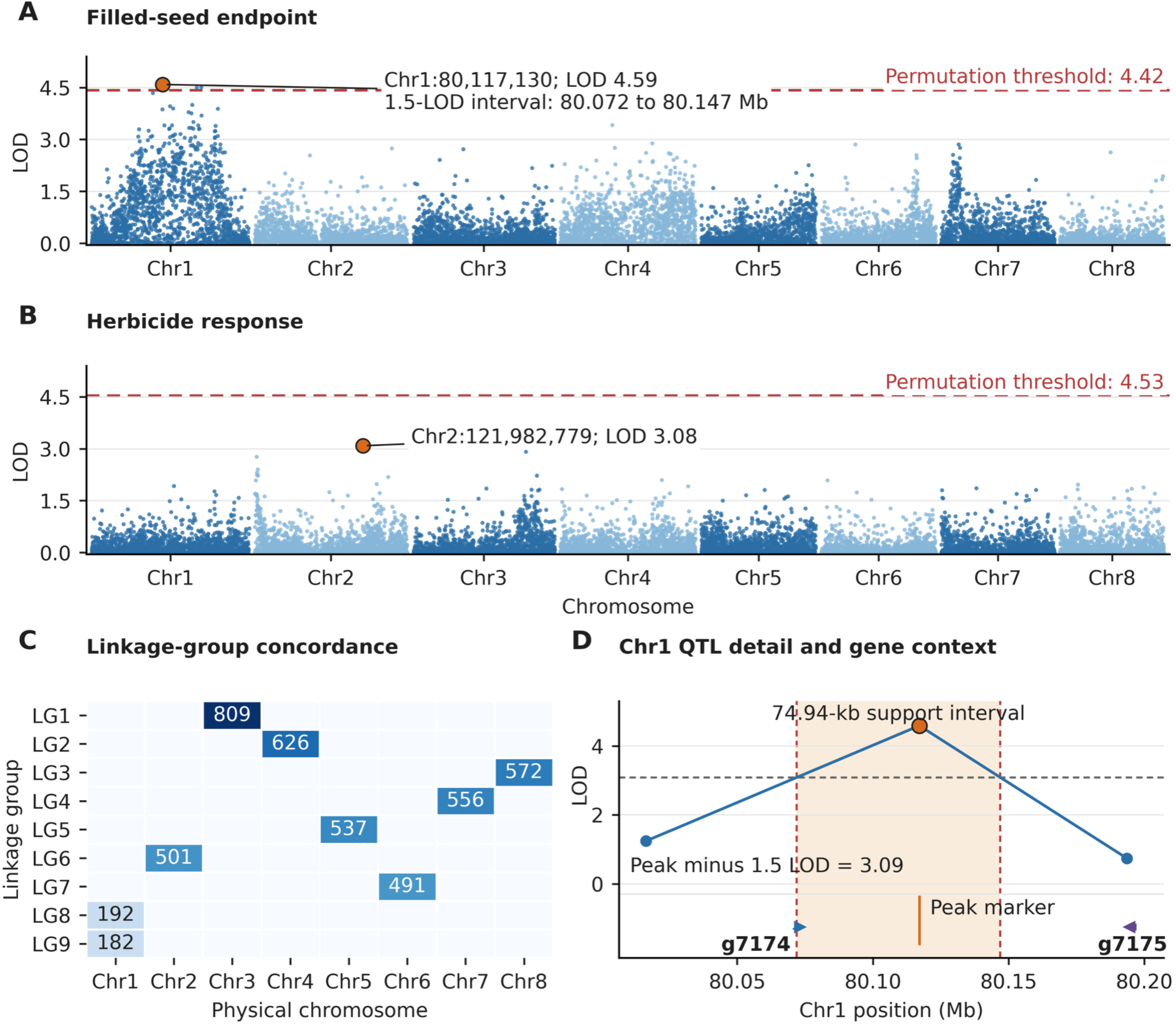
Likelihood-ratio LOD scans linkage-map correspondence, and local context of the chromosome 1 filled-seed QTL in 83 independent F2 plants. (A) The selfing-related filled-seed endpoint, scored as filled-seed production from bagged capitula in 48 filled-seed and 35 no-filled-seed plants, peaked at Chr1:80,117,130 with LOD 4.59, exceeding the 1,000-permutation threshold of 4.42. (B) Herbicide response in 53 tolerant and 30 susceptible plants peaked at Chr2:121,982,779 with LOD 3.08, below the permutation threshold of 4.53. (C) Physical-chromosome correspondence for the 4,466-marker chromosome-dominant Marey subset. The full ordered map contained 4,474 markers in nine linkage groups, with Chr1 divided between LG8 and LG9. (D) Detail of the complete physical-order Chr1 marker vector surrounding the filled-seed peak. The peak-minus-1.5-LOD cutoff was 3.09, and interpolation to the nearest flanking markers below the cutoff placed the support interval at Chr1:80,072,004 to 80,146,944, a span of 74.94 kb. The interval overlaps a single predicted protein-coding gene, *g7174*, which encodes a chloroplast-targeted phospholipase A1-like protein with a class 3 lipase domain. The peak marker is intergenic, 43.9 kb downstream of *g7174* and 77.2 kb upstream of *g7175*.

**Supplemental Figure S13.**
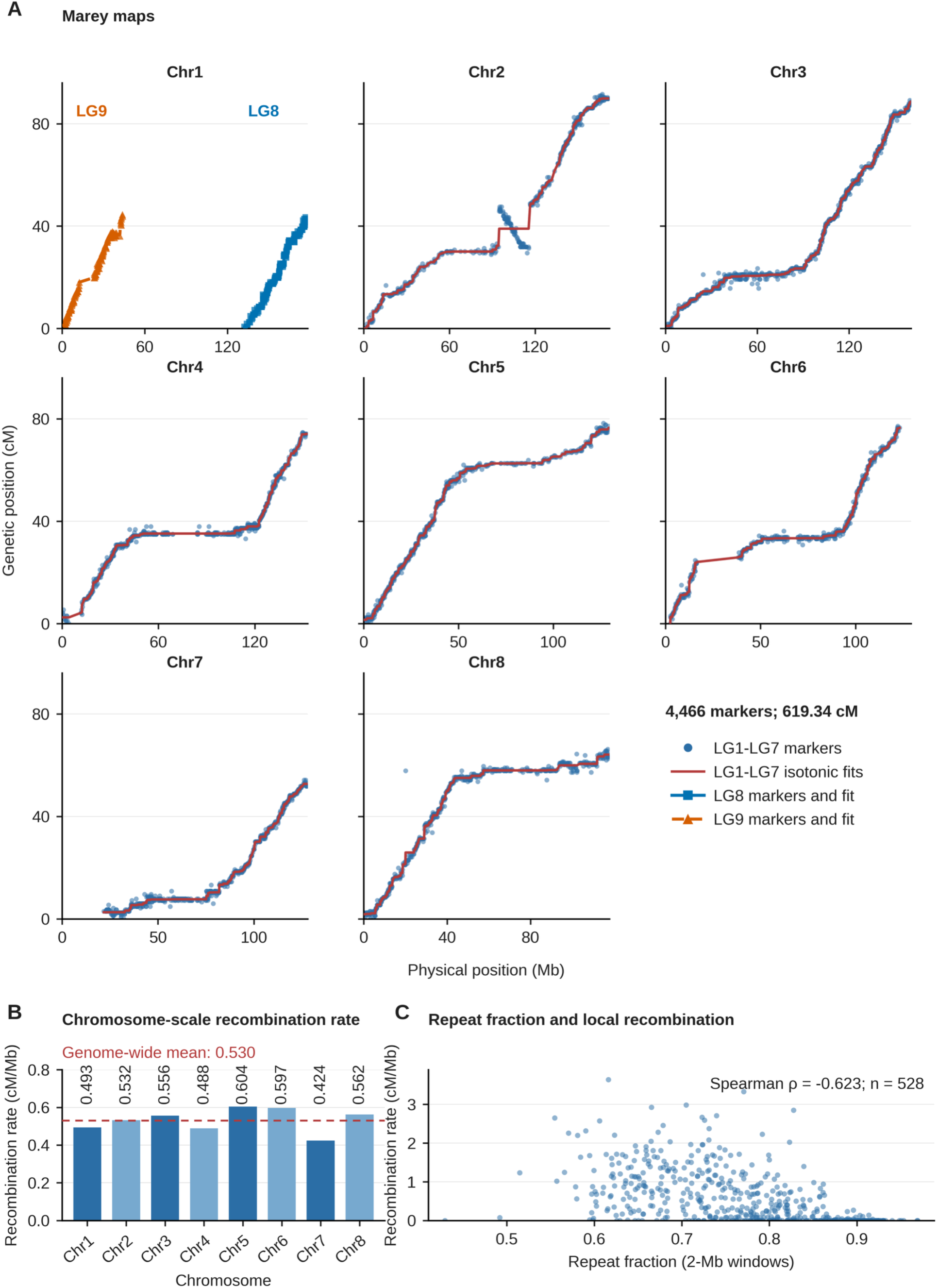
Recombination landscape from the F2-only single-copy map. Marey curves use the 4,466-marker chromosome-dominant subset (619.34 cM; 0.530 cM/Mb), whereas the full ordered map contains 4,474 markers and spans 621.75 cM. Physical windows are 2 Mb with a 2-Mb step and therefore do not overlap; 528 of 589 windows overlap a fitted marker span, including zero-rate windows, and 61 no-marker windows are NA. Repeat fraction and recombination rate have descriptive Spearman rho = -0.6227 across the 528 rate-bearing windows; no window-level inferential test was prespecified.

**Supplemental Figure S14.**
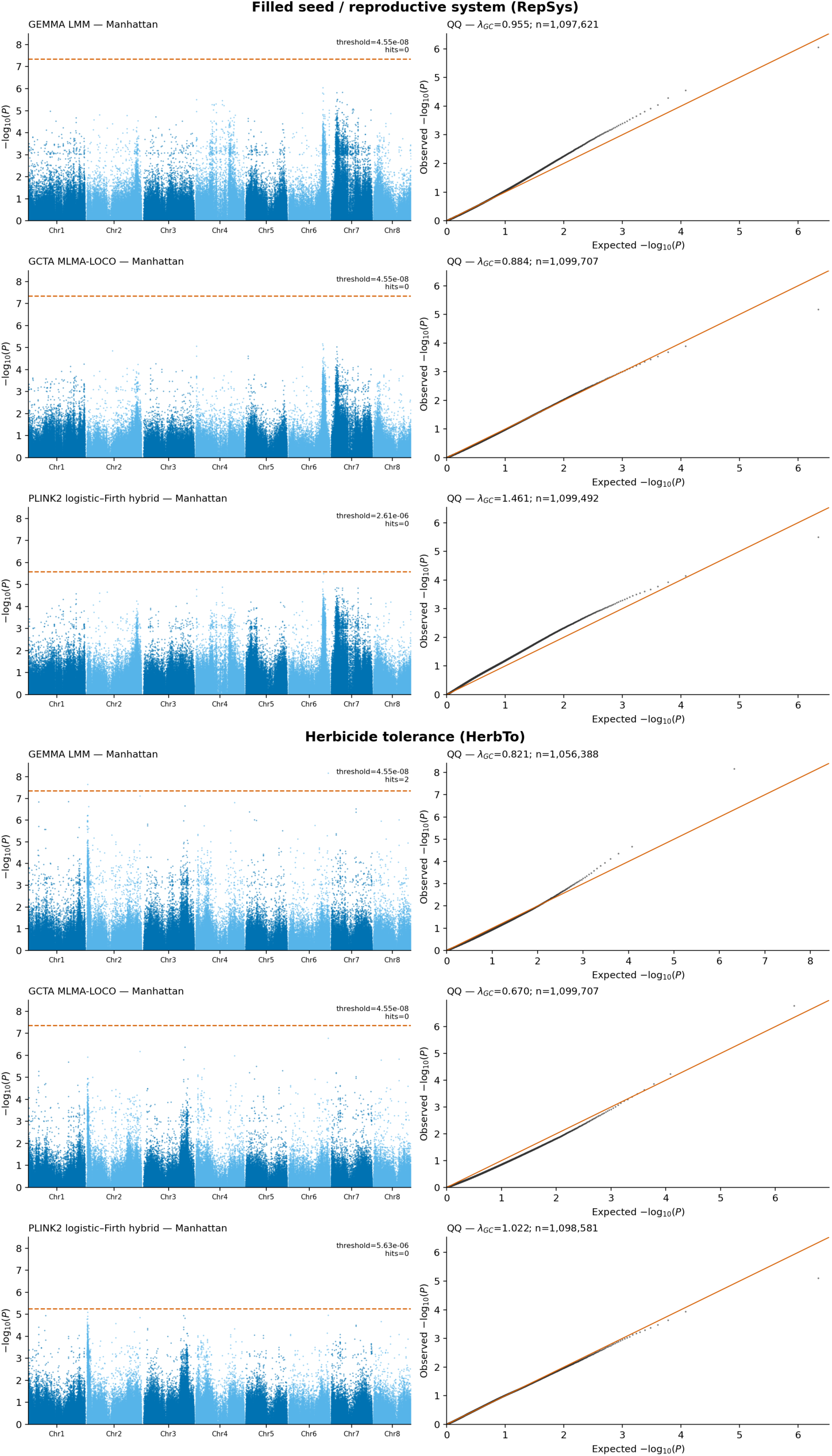
Multi-model association diagnostics for the operational filled-seed and herbicide-response endpoints. Manhattan and quantile-quantile plots are shown for GEMMA v0.98.5 linear mixed models, GCTA v1.94.1 MLMA-LOCO, and PLINK2 v2.0 logistic regression with Firth fallback, each including PC1-PC5. Dashed lines denote the model-specific genome-wide threshold. GEMMA and GCTA used a Bonferroni threshold of P = 0.05/1,099,707, whereas PLINK2 used endpoint-specific 5% genome-wide thresholds obtained from 1,000 phenotype permutations. Manhattan background points were thinned deterministically for display, while all markers with P ≤ 0.001, all leading markers, and all threshold crossings were retained. Quantile-quantile plots, λGC estimates, leading-marker summaries, and threshold counts use the complete results. Two herbicide-response markers passed the GEMMA threshold, but neither passed the corresponding GCTA or PLINK2 threshold; no exact marker was genome-wide significant in at least two models.

### Supplemental Tables

**Supplemental Table S1.** Sample and pedigree flow by ID. The first seven rows partition the 187 sequenced individuals into one sample without a complete consensus genotype, five with complete genotypes outside the final panel, and 181 final-panel members. The final row contains five unsequenced pedigree-only F₂ parents and does not contribute to the sequenced total. The table therefore lists 192 named entries, comprising 187 sequenced individuals and five additional pedigree-only nodes.

| Mutually exclusive |  |  |  |
| --- | --- | --- | --- |
| category | n | Sample IDs | Definition |
| Sequenced, no complete consensus genotype | 1 | HTS_138 | sequenced; no complete consensus genotype |
| Complete genotype, no master-trait row | 5 | HTS_060; HTS_097; HTS_110; HTS_174; Juliett | complete genotype; absent from trait table |
| Final panel: founder | 1 | HTS_006 | final panel; family founder |
| Final panel: F<sub>2</sub> | 83 | HTS_037; HTS_041; HTS_043; HTS_044; HTS_045; HTS_046; HTS_048; HTS_049; HTS_050; HTS_054; HTS_056; HTS_057; HTS_059; HTS_061; HTS_062; HTS_063; HTS_066; HTS_067; HTS_068; HTS_070; HTS_071; HTS_076; HTS_077; HTS_079; HTS_080; HTS_081; HTS_082; HTS_083; HTS_085; HTS_086; HTS_087; HTS_091; HTS_093; HTS_094; HTS_095; HTS_096; HTS_098; HTS_100; HTS_106; HTS_109; HTS_111; HTS_112; HTS_116; HTS_123; HTS_130; HTS_133; HTS_134; HTS_139; HTS_140; HTS_142; HTS_143; HTS_144; HTS_151; HTS_152; HTS_153; HTS_157; HTS_167; HTS_168; HTS_169; HTS_170; HTS_171; HTS_172; HTS_173; HTS_180; HTS_183; HTS_184; HTS_194; HTS_195; HTS_199; HTS_200; HTS_204; HTS_209; HTS_210; HTS_212; HTS_222; HTS_225; HTS_226; HTS_227; HTS_228; HTS_230; HTS_233; HTS_235; HTS_236 | final panel; independent F<sub>2</sub> used for QTL |
| category | n | Sample IDs | Definition |
| Final panel: F3 | 77 | HTS_035_4; HTS_048_1; HTS_048_2; HTS_048_3; HTS_062_1; HTS_071_1; HTS_093_1; HTS_093_2; HTS_093_3; HTS_093_4; HTS_093_5; HTS_093_8; HTS_093_9; HTS_108_1; HTS_108_10; HTS_108_2; HTS_108_3; HTS_108_4; HTS_108_5; HTS_109_1; HTS_109_2; HTS_109_3; HTS_109_4; HTS_111_1; HTS_111_2; HTS_111_3; HTS_111_4; HTS_111_5; HTS_112_1; HTS_112_2; HTS_112_4; HTS_112_5; HTS_130_1; HTS_130_11; HTS_130_2; HTS_130_3; HTS_130_4; HTS_130_9; HTS_140_1; HTS_140_2; HTS_142_2; HTS_167_1; HTS_167_2; HTS_169_1; HTS_169_2; HTS_195_2; HTS_195_3; HTS_195_4; HTS_195_5; HTS_195_9; HTS_196_1; HTS_196_2; HTS_200_1; HTS_204_1; HTS_204_2; HTS_214_1; HTS_214_10; HTS_214_11; HTS_214_12; HTS_214_2; HTS_214_3; HTS_214_4; HTS_214_5; HTS_214_6; HTS_214_8; HTS_214_9; HTS_215_1; HTS_215_2; HTS_215_3; HTS_215_4; HTS_215_5; HTS_222_1; HTS_236_1; HTS_236_10; HTS_236_18; HTS_236_3; HTS_236_8 | final panel; related F3 excluded from QTL |
| Final panel: other accessions | 20 | Bravo; ES_1; ES_2; ES_3; ES_4; HTS_017_2; HTS_017_3; HTS_017_5; IS_1; IS_2; IS_3; IS_4; IS_5; IS_6; IS_7; IS_8; OH_0061; OH_1004; VB_2; VB_3 | final panel; non-family accessions |
| Pedigree-only unsequenced F<sub>2</sub> parent | 5 | HTS_035; HTS_108; HTS_196; HTS_214; HTS_215 | pedigree node; not sequenced |

**Supplemental Table S2.** Comparative analyses. Using the eight OSU chromosome-scale scaffolds as the target for aligned-fragment comparisons and the 857-sequence OSU primary assembly before organellar filtering for the BUSCO phylogeny and uniform Miniprot projections. The final 658-sequence nuclear assembly and its representative protein set of 34,637 proteins were used for assembly-continuity statistics and OrthoFinder/CAFE5, respectively. Repeat and transposable-element occupancy were calculated on the 857-sequence primary assembly before organellar partitioning, whereas the length-weighted LAI was calculated across the eight chromosome-scale scaffolds. All eight OrthoFinder inputs contained one representative protein per gene; the protein counts reported in the table correspond to the actual OrthoFinder inputs rather than the complete isoform-level source proteomes. *T. officinale* lacked a public annotated proteome and was used only through its assembly. Genome and proteome sources were Genome Warehouse, NCBI RefSeq, or NCBI GenBank, as identified by row (Chen et al., 2021; O’Leary et al., 2016; Sayers et al., 2024). The nine-taxon BUSCO supermatrix contains 769,094 amino-acid columns, and ModelFinder selected JTT+F+I+R7 by BIC.

| Species and accession/version | Input assembly/proteome and source | Analyses |
| --- | --- | --- |
| *T. kok-saghyz* OSU; OSU TarKok 3.0 | Primary assembly before organellar filtering (857 sequences) | Aligned-fragment comparison; BUSCO phylogeny; Miniprot total-family projections |
| *T. kok-saghyz* OSU; OSU TarKok 3.0 | Organellar-filtered nuclear assembly (658 sequences) and representative proteome (34,637 proteins; one protein per gene, selected from 39,326 annotated isoforms); BRAKER3 | Assembly statistics; gene-model reconciliation; OrthoFinder/CAFE5 |
| *T. kok-saghyz* TK1151;<br>GWHBCHF | Genome and representative proteome (38,517 proteins; one longest protein per Gene = identifier, selected from 47,427 source protein records);<br>CNCB-GWH | Aligned-fragment comparison; synteny;<br>BUSCO phylogeny; Miniprot;<br>OrthoFinder/CAFE5 |
| *T. mongolicum*; GWHBCHG | Genome and representative proteome (38,214 proteins; one longest protein per Gene = identifier, selected from 54,708 source protein records);<br>CNCB-GWH | Aligned-fragment comparison; synteny;<br>BUSCO phylogeny; Miniprot;<br>OrthoFinder/CAFE5 |
| *T. officinale*; GCA_950097615 | Genome; NCBI GenBank; no annotated proteome used in OrthoFinder/CAFE5 | Aligned-fragment comparison; BUSCO phylogeny; Miniprot |
| *L. sativa*; GCF_002870075.5 | Genome and representative proteome (35,323 proteins; one longest protein per annotated gene, selected from 49,518 source protein records);<br>NCBI RefSeq | Aligned-fragment comparison; synteny;<br>BUSCO phylogeny; Miniprot;<br>OrthoFinder/CAFE5 |
| *L. saligna*; GCF_052625555.1 | Genome and representative proteome (31,321 proteins; one longest protein per annotated gene, selected from 41,179 source protein records);<br>NCBI RefSeq | Aligned-fragment comparison; BUSCO phylogeny; Miniprot; OrthoFinder/CAFE5 |
| *L. virosa*; GCF_052849685.1 | Genome and representative proteome (35,719 proteins; one longest protein per annotated gene, selected from 47,227 source protein records);<br>NCBI RefSeq | Aligned-fragment comparison; BUSCO phylogeny; Miniprot; OrthoFinder/CAFE5 |
| *C. intybus*; GCA_023525715.1 | Genome and representative proteome (43,721 proteins; one protein per annotated gene; the source proteome was already gene-representative);<br>NCBI GenBank | Aligned-fragment comparison; BUSCO phylogeny; Miniprot; OrthoFinder/CAFE5 |
| *C. endivia*; GCA_023376185.1 | Genome and representative proteome (51,881 proteins; one protein per annotated gene; the source proteome was already gene-representative);<br>NCBI GenBank | Aligned-fragment comparison; BUSCO phylogeny; Miniprot; OrthoFinder/CAFE5 |

**Supplemental Table S3.** Structural, functional, non-coding RNA, and BUSCO completeness summary for OSU TarKok 3.0.

Panel A — Gene models
| Metric | Value | Source / note |
| --- | --- | --- |
| Protein-coding genes | 34637 | Final nuclear BRAKER3 annotation |
| Transcripts (mRNAs) | 39326 | Final nuclear BRAKER3 annotation |
| Mean transcripts per gene | 1.14 | Computed as 39326 / 34637 |
| Genes on the eight chromosome-scale scaffolds (n; %) | 34610;<br>99.92% | The eight scaffolds contain 98.0% of the assembled nuclear sequence |
| **Genes on unplaced sequences (n; %)** | 27;<br>0.08% | 650 unplaced nuclear sequences in the final 658-sequence assembly |
| **Mean / median gene length (bp)** | 2129.45<br>/ 1370 | Inclusive locus span from first to last annotated coordinate |
| **Mean / median CDS length (bp)** | 1082.35<br>/ 885 | Summed coding sequence per transcript across 39326 transcripts |
| **Mean / median representative-protein length (aa)** | 354.96<br>/ 288 | 34637 representative proteins; one selected protein per gene |
| **Mean coding exons per transcript** | 4.67 | Number of CDS features per transcript |
| **Mean / median intron length (bp)** | 308.69<br>/ 94 | 144335 annotated introns |
| **Monoexonic transcripts (n; %)** | 9940;<br>25.28% | Descriptive annotation statistic; monoexonic status alone does not demonstrate gene-model fragmentation |
| **Transcripts with any annotated UTR (n; %)** | 19314;<br>49.11% | UTR prediction depends on RNA-seq depth; calculated after intersecting the UTR-augmented annotation with the final transcript set |
| **Transcripts with annotated 5' and 3' UTRs (n; %)** | 17251;<br>43.87% | Transcripts containing both UTR feature types |
| **Mean gene density on the eight chromosome-scale scaffolds (genes Mb<sup>-1</sup>)** | 29.62 | Computed as 34610 genes / 1168.509718 Mb; provides context for the mappable-genome analysis |

Panel B — Functional annotation
| **Category** | **n** | **Proteins, n (%)** | **Tool / definition** |
| --- | --- | --- | --- |
| **At least one InterPro annotation** | 28035 | 80.94% | InterProScan v5.78-109.0; at least one InterPro entry |
| **Pfam domain assignment** | 26171 | 75.56% | InterProScan; at least one Pfam hit |
| **At least one Gene Ontology term** | 24132 | 69.67% | Union of InterProScan- and eggNOG-derived GO assignments |
| **eggNOG orthologous-group assignment** | 30902 | 89.22% | eggNOG-mapper v2.1.12 |
| **KEGG orthology or pathway assignment** | 17071 | 49.29% | Union of eggNOG KO, eggNOG pathway, and significant Kofam assignments |
| **Any core functional assignment** | 31128 | 89.87% | Union of InterPro, Pfam, GO, eggNOG orthologous-group, and KEGG assignments |
| **No core functional assignment** | 3509 | 10.13% | Complement of the core functional-assignment union |
| **Any verified functional evidence** | 32494 | 93.81% | Core union plus verified best-hit accessions, dbCAN, iTAK, plantiSMASH, targeted classifications, or positive TargetP localization |
| **No verified functional evidence** | 2143 | 6.19% | Complement of the inclusive verified-evidence definition |
| **Predicted transcription factors** | 1826 | 5.27% | iTAK; provides annotation context for the transcription-factor nodes displayed in Figure 5B |

**Panel C — Non-coding RNA**
| Feature class | n | Tool / note |
| --- | --- | --- |
| tRNA predictions, non-pseudogene | 698 | tRNAscan-SE v2.0.12 |
| tRNA pseudogene predictions | 124 | tRNAscan-SE v2.0.12 |
| rRNA-model predictions: 5S / 5.8S / 18S or eukaryotic SSU / 28S or eukaryotic LSU | 7691 / 2681 / 2812 / 3084 | barrnap v0.9 and Infernal/Rfam; uncurated computational prediction rows |
| Other rRNA-model predictions | 38 | barrnap and Infernal/Rfam; uncurated computational prediction rows |
| Total rRNA-model predictions | 16306 | Not interpreted as a curated biological rRNA gene count because the abundance indicates extensive repeat-associated matches |
| snRNA predictions | 175 | Infernal v1.1.5 / Rfam |
| snoRNA predictions | 920 | Infernal v1.1.5 / Rfam |
| miRNA predictions | 98 | Infernal v1.1.5 / Rfam |
| Other structured Rfam-family predictions | 309 | Infernal v1.1.5 / Rfam |
| Total additional structured non-rRNA predictions | 1502 | Sum of snRNA, snoRNA, miRNA, and other Rfam-family predictions |

**Panel D — Completeness of the annotated gene set**
| Metric | Value | Note |
| --- | --- | --- |
| **Protein-mode BUSCO complete** | 97.5% (2734) | eudicotyledons_odb12, n = 2805; final 34637-protein representative proteome |
| **single-copy** | 92.1% (2583) | Reported in Supplemental Methods and Table 1 |
| **duplicated** | 5.4% (151) | Protein-mode BUSCO |
| **fragmented** | 1.3% (37) | Protein-mode BUSCO |
| **missing** | 1.2% (34) | Protein-mode BUSCO |
| **Genome-mode BUSCO complete** | 98.4%<br>(2759) | eudicotyledons_odb12, n = 2805; final 658-sequence nuclear assembly |
| **single-copy** | 91.6%<br>(2569) | Genome-mode BUSCO; assembly rather than annotation |
| **duplicated** | 6.8%<br>(190) | Genome-mode BUSCO; assembly rather than annotation |
| **fragmented** | 1.1%<br>(30) | Genome-mode BUSCO; assembly rather than annotation |
| **missing** | 0.6%<br>(16) | Genome-mode BUSCO; assembly rather than annotation |
| **RNA-seq evidence used for gene prediction** | 12 libraries | BRAKER3 integrated RNA-seq evidence; a validated final per-gene support percentage was not calculated and is therefore not reported |

**Supplemental Table S4.** Length-weighted nucleotide identity among retained best alignments of 5-kb comparator-query fragments to the eight OSU chromosome-scale scaffolds target before organellar filtering. Alignments shorter than 2 kb were excluded. Identity is conditional on retained alignments; aligned fraction is retained aligned query length divided by the corresponding comparator assembly length.

| Comparison | Aligned-fragment |  |  |  |
| --- | --- | --- | --- | --- |
|  | identity | Aligned fraction | Fragment size | Minimum aligned |
| *T. kok-saghyz* TK1151 | 98.74% | 78.20% | 5 kb | 2 kb |
| *T. mongolicum* | 95.22% | 49.28% | 5 kb | 2 kb |
| *T. officinale* | 95.36% | 47.17% | 5 kb | 2 kb |

**Supplemental Table S5.** Predominant lettuce-chromosome components of anchored syntenic length. Percentages are shares of each OSU chromosome’s anchored length; residual components are retained. The classifications summarize correspondence and do not define a unique rearrangement or fusion history.

| OSU |  |  |
| --- | --- | --- |
| chromosome | Lettuce shares of anchored length | Predominant-component classification |
| Chr1 | Chr5 62.697%; Chr6 37.303% | two components |
| Chr2 | Chr4 99.856%; Chr1 0.144% | ≥99.3% on one chromosome |
| Chr3 | Chr2 75.112%; Chr9 17.907%; Chr6 4.365%; Chr5 2.617% | two major components plus residuals |
| Chr4 | Chr8 99.332%; Chr2 0.471%; Chr3 0.062%; Chr9 0.052%; Chr4 0.043%; Chr1 0.041% | ≥99.3% on one chromosome |
| Chr5 | Chr3 100.000% | one chromosome |
| Chr6 | Chr9 69.633%; Chr7 30.101%; Chr8 0.266% | two major components plus residual |
| Chr7 | Chr7 52.383%; Chr6 34.451%; Chr5 13.166% | three components |
| Chr8 | Chr1 100.000% | one chromosome |

**Supplemental Table S6.** Rubber-pathway locus expression in the 12-library in-house dataset. TPM values are locus-level; *g12582* combines *TkCPT1*-like and *TkCPT2*-like isoforms and is not paralog-specific. *g5630* is supported as *TkCPTL1*, *g9503* is a probable CPTL/RTA paralog, five loci are citrate-binding protein homologs, and *g21989* is REF. Exactly three SRPP loci meet the prespecified underground-tissue-enrichment criterion of underground:aerial ratio >1 and DESeq2 adjusted P <0.05: *g260*, *g261*, and *g262*. *g259* has adjusted P = 0.171 and does not meet the criterion. Differential-expression adjusted P values compare underground and aerial tissues; latex is not present in this dataset.

| Locus | Family | Leaf | Flower | Seed/floral | Crown | Root | Underground: |  |
| --- | --- | --- | --- | --- | --- | --- | --- | --- |
|  |  |  |  |  |  |  | aerial ratio | DESeq2 padj |
| *g12582* | CPT | 5.00 | 112.51 | 8.47 | 270.37 | 325.43 | 6.95 | 0.00971 |
| *g21319* | CPT | 10.71 | 9.52 | 11.70 | 12.21 | 16.22 | 1.31 | 0.0128 |
| *g4125* | CPT | 19.80 | 5.54 | 1.94 | 1.18 | 1.63 | 0.24 | 0.0146 |
| *g21195* | CPT | 3.99 | 0.10 | 0.05 | 0.10 | 0.10 | 0.46 | 0.497 |
| *g4124* | CPT | 0.21 | 0.03 | 0.15 | 0.02 | 0.00 | 0.89 | 0.0515 |
| *g21196* | CPT | 0.00 | 0.00 | 0.00 | 0.00 | 0.00 | 1.00 | not tested |
| *g5630* | CPTL/<br>RTA | 94.49 | 10.72 | 22.58 | 291.68 | 170.09 | 5.32 | 0.0244 |
| *g9503* | CPTL/<br>RTA | 11.96 | 10.54 | 11.87 | 11.35 | 16.63 | 1.20 | 0.0536 |
| *g259* | SRPP | 669.74 | 146.58 | 140.53 | 1509.17 | 424.26 | 3.02 | 0.171 |
| *g262* | SRPP | 21.14 | 0.72 | 1.70 | 542.55 | 634.76 | 66.59 | 0.0274 |
| *g261* | SRPP | 23.47 | 0.53 | 12.40 | 718.85 | 387.45 | 42.20 | 9.6e-08 |
| *g260* | SRPP | 35.42 | 2.82 | 6.80 | 163.57 | 120.35 | 8.93 | 9.7e-04 |
| *g3901* | SRPP | 875.40 | 125.18 | 151.55 | 123.71 | 134.16 | 0.34 | 0.115 |
| *g263* | SRPP | 1030.94 | 2.31 | 5.83 | 76.89 | 30.42 | 0.16 | 0.0382 |
| *g9592* | SRPP | 36.84 | 0.00 | 14.15 | 0.32 | 0.05 | 0.07 | 0.00351 |
| *g21989* | REF | 10.62 | 0.15 | 0.19 | 69.26 | 33.82 | 11.29 | 0.0555 |
| *g25523* | CitBP | 0.00 | 0.00 | 0.01 | 0.48 | 0.00 | 1.24 | 0.119 |
| *g2713* | CitBP | 0.02 | 0.45 | 0.09 | 0.24 | 0.19 | 1.02 | 0.628 |
| *g2714* | CitBP | 0.00 | 0.00 | 0.00 | 0.01 | 0.00 | 1.00 | not tested |
| *g25524* | CitBP | 0.00 | 0.00 | 0.00 | 0.00 | 0.00 | 1.00 | not tested |
| *g30219* | CitBP | 0.00 | 0.00 | 1.05 | 0.00 | 0.00 | 0.74 | not tested |
| *g21546* | HMGR | 722.04 | 0.95 | 452.56 | 1563.19 | 940.55 | 3.19 | 0.316 |
| *g21555* | HMGR | 49.30 | 4.24 | 85.26 | 58.71 | 197.15 | 2.73 | 0.0675 |
| *g12555* | HMGR | 4.29 | 33.53 | 29.23 | 85.52 | 63.50 | 3.23 | 0.0133 |
| *g21556* | HMGR | 38.82 | 0.27 | 47.76 | 36.87 | 98.66 | 2.30 | 0.233 |
| *g25642* | HMGR | 15.11 | 4.38 | 52.80 | 56.57 | 23.16 | 1.63 | 0.338 |
| *g21551* | HMGR | 1.85 | 0.04 | 27.03 | 2.82 | 10.92 | 0.74 | 0.0654 |
| *g2272* | HMGR | 3.74 | 26.05 | 107.42 | 3.15 | 7.22 | 0.13 | 0.00169 |
| *g23511* | HMGR | 1.98 | 4.31 | 20.99 | 3.12 | 1.45 | 0.33 | 0.0245 |
| *g17679* | HMGR | 0.01 | 0.02 | 1.56 | 0.22 | 0.76 | 0.97 | 0.0168 |
| *g21548* | HMGR | 0.00 | 0.00 | 1.62 | 0.06 | 0.87 | 0.95 | 3.8e-04 |
| *g21553* | HMGR | 13.07 | 0.00 | 0.73 | 0.23 | 0.38 | 0.23 | 0.0343 |
| *g21547* | HMGR | 6.45 | 0.05 | 1.36 | 0.12 | 0.04 | 0.30 | 1.8e-04 |
| *g28716* | HMGR | 0.02 | 0.03 | 0.15 | 0.02 | 0.00 | 0.95 | not tested |
| *g21554* | HMGR | 5.33 | 0.00 | 2.68 | 0.01 | 0.01 | 0.27 | 8.2e-05 |
| *g7279* | HMGR | 0.00 | 0.00 | 5.41 | 0.00 | 0.00 | 0.36 | not tested |

**Supplemental Table S7.** plantiSMASH biosynthetic-gene-cluster predictions and overlap with curated rubber-pathway loci.

Panel A
| Cluster type | Clusters | Genes |
| --- | --- | --- |
| Saccharide | 11 | 137 |
| Cyclopeptide | 5 | 215 |
| Terpene | 5 | 73 |
| Putative | 5 | 71 |
| Alkaloid | 5 | 75 |
| Terpene-saccharide | 4 | 56 |
| Fatty acid | 2 | 25 |
| Polyketide | 2 | 17 |
| Lignan | 1 | 10 |
| Saccharide-polyketide | 1 | 7 |
| Transporter-associated polyketide-saccharide | 1 | 20 |
| Total | 42 | 706* |

Panel B
| Rubber-related locus | Family | Predicted cluster type | Interpretation |
| --- | --- | --- | --- |
| *g12582* | CPT | Lignan | Only curated rubber-related locus overlapping a predicted plantiSMASH cluster |

\* Gene counts are gene-cluster assignments; 20 genes belong to two clusters, so the 706 assignments involve 686 unique genes.

**Supplemental Table S8.** Rubber-related family projections and OrthoFinder/ CAFE5 results. The OrthoFinder analysis using the final OSU representative protein set assigned 32,795 OSU proteins to 33,849 orthogroups and left 1,842 unassigned. After excluding 27 families with more than 100 copies, CAFE5 tested 24,025 of 33,822 input families after filtering 9,797 inferred absent at the root.

Panel A
| Gene family | OSU<br>TarKok<br>3.0 | TK1151 | T.<br>mongolicu<br>m | T.<br>officinal<br>e | C.<br>intybu<br>s | C.<br>endivi<br>a | L.<br>sativa | L.<br>saligna | L.<br>virosa |
| --- | --- | --- | --- | --- | --- | --- | --- | --- | --- |
| cis-Prenyltransferase (CPT) | 7 | 8 | 5 | 7 | 6 | 7 | 11 | 9 | 14 |
| cis-Prenyltransferase-like/rubber transferase activator (CPTL/RTA) | 2 | 2 | 2 | 2 | 2 | 2 | 3 | 2 | 2 |
| Citrate-binding protein (CitBP) | 6 | 4 | 6 | 7 | 6 | 5 | 6 | 6 | 6 |
| Small rubber particle protein (SRPP) | 10 | 17 | 6 | 7 | 22 | 9 | 10 | 10 | 9 |
| Rubber elongation factor (REF) | 1 | 1 | 1 | 1 | 1 | 1 | 1 | 1 | 1 |
| 3-Hydroxy-3-methylglutaryl-CoA reductase (HMGR) | 15 | 12 | 13 | 18 | 16 | 16 | 7 | 7 | 7 |

Note. Counts represent distinct homology clusters recovered by uniform Miniprot mapping with minimum protein identity of 0.60 and minimum coverage of 0.50. OSU TarKok 3.0 and TK1151 are assemblies of Taraxacum kok-saghyz. These broad projections are distinct from the curated OSU expression inventory and from the global orthogroups tested with CAFE5 in Panel B. No surveyed family has a uniquely highest count in OSU TarKok 3.0.

Panel B
| Quantity | Result |
| --- | --- |
| OSU proteome | 34,637 proteins; one representative per nuclear GFF gene |
| OrthoFinder | 33,849 orthogroups; 32,795 OSU proteins assigned and 1,842 unassigned |
| CAFE5 input | 33,822 families after excluding 27 large families; 9,797 root-absent families filtered; 24,025 tested |
| CAFE5 global model | single global lambda = 0.0025869305101934; no error model; -lnL = 215099 |
| Family tests | 1,929 with raw family P <0.05; 891 with family-level BH q <0.05 across 24,025 tests |
| HMGR-containing OG0000471 | OSU change +4; raw family P = 0.008; BH q = 0.160; raw OSU branch probability = 0.00644572 |
| Focal-family reconciliation | 24 manually reconciled rows mapped to 23 global orthogroups |

**Supplemental Table S9.** Full-vector support for the connected, marker-interpolated 1.5-LOD interval in the 83-F2 logistic likelihood-ratio scan. Prose values are rounded; the table retains full precision for marker LOD values, cutoff, and crossings.

| Entry | Marker/coordinate | LOD or criterion | Interpolated result |
| --- | --- | --- | --- |
| Filled-seed endpoint peak | scf00003:80117130_T | 4.5868413968128 | Chr1:80,117,130 |
| Permutation threshold | 1,000 permutations | 4.42393420637542 | significant |
| 1.5-LOD cutoff | 1,652-marker complete Chr1 vector | 3.0868413968128 | connected peak component |
| Left exterior marker | scf00003:80016532_C (80016532 bp) | 1.24291661737745 | below cutoff |
| Left interior marker | scf00003:80117130_T (80117130 bp) | 4.5868413968128 | crossing 80072004.2839767 bp |
| Right interior marker | scf00003:80117130_T (80117130 bp) | 4.5868413968128 | crossing 80146943.8412337 bp |
| Right exterior marker | scf00003:80193587_A (80193587 bp) | 0.740121375029608 | below cutoff |
| Connected interval | Chr1:80,072,004-80,146,944 | marker-interpolated 1.5-LOD | 74939.5572569966 bp |
| Interval-overlapping gene | *g7174*; Chr1:80,071,643–80,073,251 (+) | n.a. | Predicted chloroplast-targeted phospholipase A1-like protein with a class 3 lipase domain; overlaps 1,248 bp of the left interval boundary |
| Peak genomic context | Chr1:80,117,130 | intergenic | 43,879 bp downstream of *g7174* and 77,177 bp upstream of *g7175* |
| Herbicide response | Chr2:121,982,779 | LOD 3.08434689751284; threshold 4.53430762448663 | not significant |

**Supplemental Table S10:**
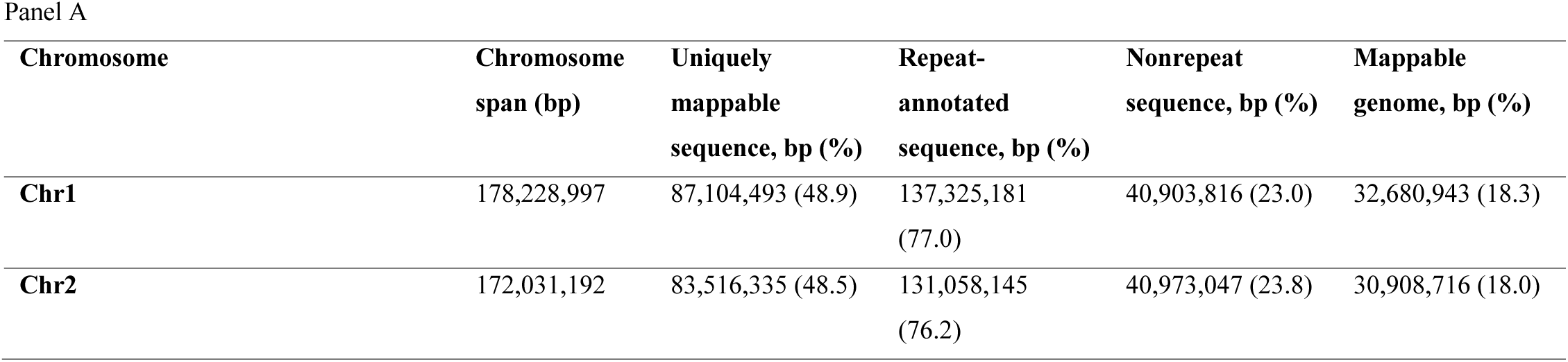
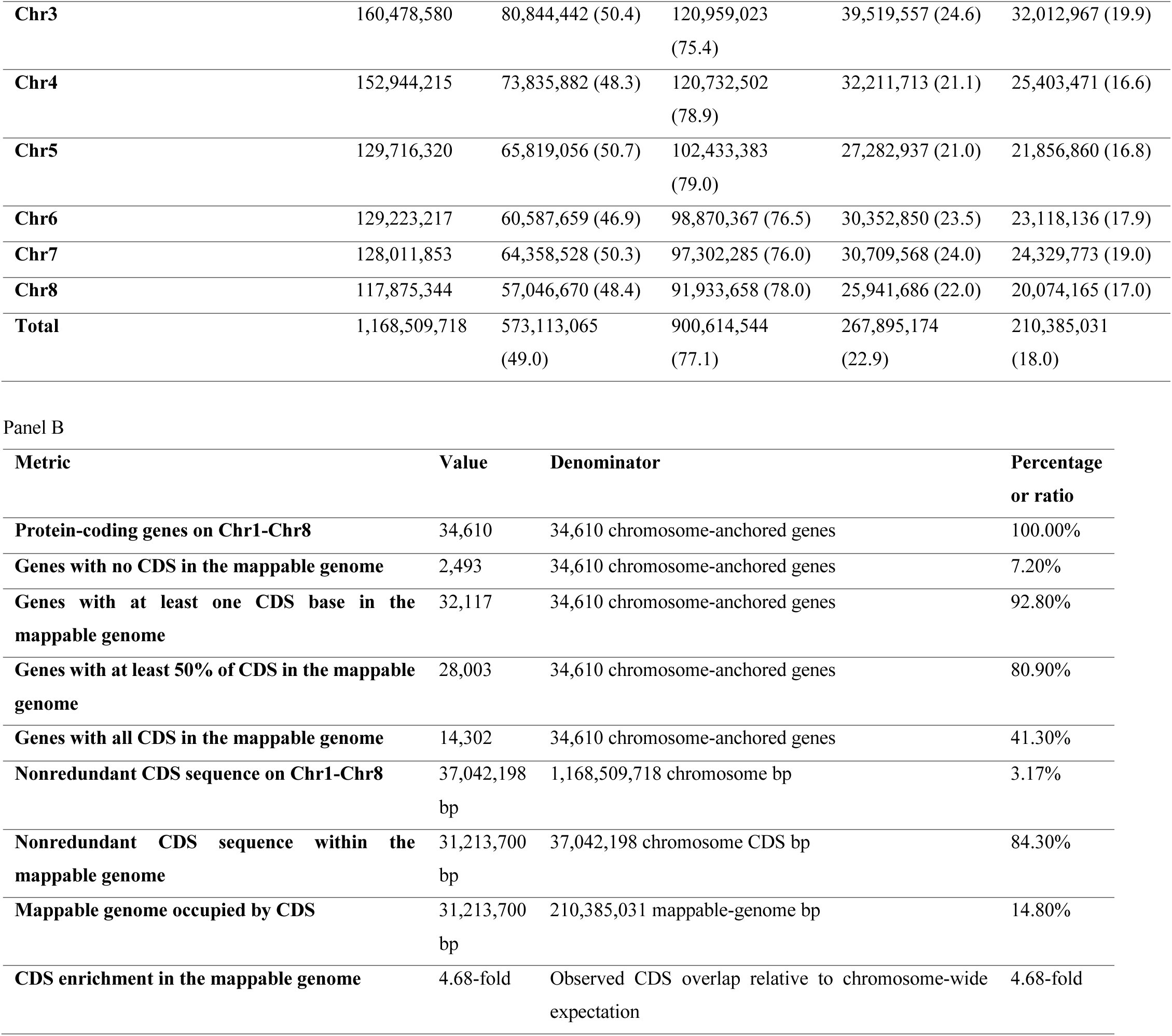
Chromosome mappability and coding-sequence accessibility in OSU TarKok 3.0. Panel A reports sequence classified as uniquely mappable by GenMap, repeat-annotated by RepeatMasker, nonrepeat, and part of the mappable genome. The mappable genome was defined as sequence that was both uniquely mappable and nonrepeat. Panel B summarizes the accessibility of coding sequence within this genome. Gene percentages use the 34,610 protein-coding genes located on Chr1-Chr8 as their denominator; the 27 genes on unplaced sequences were excluded.

**Supplemental Table S11.** Multi-model population-association summary for the operational filled-seed and herbicide-response endpoints. . GEMMA v0.98.5 linear mixed models, GCTA v1.94.1 MLMA-LOCO, and PLINK2 v2.0 logistic regression with Firth fallback included PC1-PC5. GEMMA and GCTA used a Bonferroni threshold of P = 0.05/1,099,707, whereas PLINK2 used endpoint-specific 5% genome-wide thresholds obtained from 1,000 phenotype permutations. The genomic-inflation factor, λGC, was calculated from every valid P value in each primary output. Cross-model support required the same chromosome:position:reference:alternate allele record to pass its model-specific threshold in at least two models.

| **Endpoint** | **Model** | **n** | **Markers with valid P** | **λGC** | **Genome-wide threshold** | **Lead marker** | **Lead effect** | **Lead P** | **Genome-wide hits** |
| --- | --- | --- | --- | --- | --- | --- | --- | --- | --- |
| **Filled seed** | GEMMA linear mixed model | 166 | 1,097,621 | 0.955 | Bonferroni P = 4.5467 × 10<sup>-8</sup> | Chr6:107,995,347 G>A | β = -0.271244 | 8.9343 × 10<sup>-7</sup> | 0 |
| **Filled seed** | GCTA MLMA-LOCO | 166 | 1,099,707 | 0.884 | Bonferroni P = 4.5467 × 10<sup>-8</sup> | Chr6:106,968,545 A>C | β = 0.290115 | 6.6695 × 10<sup>-6</sup> | 0 |
| **Filled seed** | PLINK2 logistic regression with Firth fallback | 166 | 1,099,492 | 1.461 | Permutation P = 2.61345 × 10<sup>-6</sup> | Chr6:109,760,720 G>A | OR = 0.214734 | 3.1669 × 10<sup>-6</sup> | 0 |
| **Herbicide response** | GEMMA linear mixed model | 177 | 1,056,388 | 0.821 | Bonferroni P = 4.5467 × 10<sup>-8</sup> | Chr6:124,035,144 T>G | β = -0.400286 | 6.7563 × 10<sup>-9</sup> | 2 |
| **Herbicide response** | GCTA MLMA-LOCO | 177 | 1,099,707 | 0.67 | Bonferroni P = 4.5467 × 10<sup>-8</sup> | Chr6:124,035,144 T>G | β = -0.361516 | 1.6579 × 10<sup>-7</sup> | 0 |
| Herbicide<br>response | PLINK2<br>logistic<br>regression with<br>Firth fallback | 177 | 1,098,581 | 1.022 | Permutation P = 5.62516 × 10<sup>-6</sup> | Chr2:3,099,459 G>A | OR = 0.176071 | 7.9604 × 10<sup>-6</sup> | 0 |

## Notes

https://www.rubber-dandelion-genomics.org

https://github.com/Fresnedo-Lab/OSU-TarKok-3.0

https://doi.org/10.5281/zenodo.23585451

### Competing Interest Statement

The authors have declared no competing interest.

